# Biochemical Profiling and Structural Basis of ADAR1-Mediated RNA Editing

**DOI:** 10.1101/2025.01.02.631069

**Authors:** Xiangyu Deng, Lina Sun, Min Zhang, Rashmi Basavaraj, Jin Wang, Yi-Lan Weng, Yang Gao

## Abstract

ADAR1 is a pivotal regulator in RNA-induced immune responses by catalyzing the conversion of adenosine to inosine on double-stranded RNA. Mutations on ADAR1 are associated with human autoimmune disease, and targeting ADAR1 has been proposed for cancer immunotherapy. However, the molecular mechanisms governing ADAR1-mediated RNA editing remain enigmatic. Here, we provide detailed biochemical and structural characterizations of human ADAR1. Our biochemical profiling reveals that ADAR1 editing is both sequence and RNA duplex length-dependent, but can well tolerate mismatches near the editing site. Moreover, our high-resolution structures of ADAR1-RNA complexes, coupled with mutagenesis studies, revealed the molecular basis for RNA binding, substrate selection, dimerization, and the crucial role of the RNA-binding domain 3 for ADAR1 editing. The ADAR1 structures also help explain the potential defects of disease-associated mutations, where biochemical and RNA-sequencing analysis further indicate some of the mutations preferentially impact the editing of RNAs with short duplex. Our findings illustrate the molecular mechanism of ADAR1 editing and provide clues for deciphering its role in immune regulation and drug targeting.

**HIGHLIGHTS:**

- Biochemical profiling of ADAR1 RNA substrate preference
- Atomic resolution structures of ADAR1 with two physiological RNA substrates
- Disease-related mutations of ADAR1 preferentially impact RNA editing with short dsRNA.
- RNA-binding domain 3 is essential for ADAR1 RNA capture and editing

## Introduction

Adenosine deaminase acting on dsRNA 1 (ADAR1) is an essential immune surveillance factor that catalyzes the post-transcriptional conversion of Adenosine (A) to Inosine (I) on double-stranded (ds)RNA^1,2^. ADAR1 editing of dsRNA effectively counteracts the activation of various nucleic acid sensors, thereby averting aberrant immune response^3^. Notably, ADAR1 knockout in mouse models results in embryonic lethality, while mutations in ADAR1 are linked to Aicardi–Goutières syndrome (AGS), a severe interferonopathy characterized by the heightened interferon responses^4–8^. Intriguingly, ADAR1 knockdown significantly enhances the susceptibility of tumor cells to immune checkpoint blockade treatment^9–11^, making ADAR1 a promising target for bolstering cancer immunotherapy. Additionally, biotechnological tools leveraging the specific RNA editing of ADAR1 have been developed for RNA base editing, sensing, and disease treatment^12–14^.

ADAR1 exists in two isoforms: the interferon-inducible full-length ADAR1 (ADAR1-p150) and the constitutively expressed shorter isoform (ADAR1-p110). ADAR1-p150 contains two left-handed Z-nucleic acid (Z-RNA) binding domains (ZBD-𝘢, and ZBD-β), and three dsRNA binding domains (RBD-I, II, III) besides the deaminase domain, whereas the ADAR1-p110 lacks the ZBD-α and a nuclear export signal (NES)^15^. Additionally, there exists ADAR1 homolog ADAR2, which contains a deaminase domain, two dsRBDs, but no ZBDs^15^. Notably, ADAR1-p150 has a distinct substrate specificity from ADAR1-p110 or ADAR2 and exerts dominant effects in modulating innate immunity^3,5,6,16,17^. Previous studies suggested that the deaminase domain of ADAR1 plays a major role in defining its RNA selection^18–21^. In addition, the architecture of RBDs, intracellular localization of ADAR1, and the unique ZBDs have also been proposed to contribute to ADAR1’s substrate specificity^20,22–24^. Deep RNA sequencing has identified a large number of endogenous RNAs that are possibly edited by ADAR1^16,25–30^, and multiple lines of research have suggested that both the RNA sequence and secondary structure are important for regulating ADAR editing^15,21,31–36^. However, the precise sequence and structural determinants of RNA substrates governing ADAR1 recognition and editing are not well defined. Moreover, the structural basis of ADAR1 action and the molecular basis of ADAR1 disease-related mutations remain poorly understood.

In this study, we performed comprehensive biochemical profiling to elucidate the substrate preference of ADAR1, revealing its dependence on both RNA sequence and duplex length, alongside a remarkable tolerance of mismatches near the editing site. In addition, we presented atomic resolution structures of ADAR1 with two different RNA substrates, unveiling residues crucial for ADAR1 RNA binding, editing, and dimerization. Our results also highlighted the critical role of RBD3 in ADAR1 RNA binding and editing. Furthermore, our biochemical characterizations and RNA-sequencing (RNA-Seq) analysis of AGS-associated ADAR1 mutations demonstrated that many of the ADAR1 mutations preferentially affect ADAR1 editing on short duplex RNA. In summary, our work offers crucial molecular insights into ADAR1-mediated RNA editing. The results will be essential in deciphering ADAR1’s regulation of RNA-induced immune responses and promoting targeting and engineering of ADAR1 for biomedical applications.

## Results

### Biochemical profiling of ADAR1 RNA substrate preference

We aim to systematically investigate the ADAR1’s substrate preference with *in vitro* biochemical assays. However, the ADAR1-p150 is challenging to obtain due to its low yield and a strong tendency to aggregate during purification. Based on AlphaFold prediction^37^, we designed several ADAR1 N-terminal truncations by deleting the first 33, 67, 100, or 126 residues of the disordered region while keeping all the ZBDs and RBDs (Figure S1A). Truncating the first 126 residues on the N-terminus (ADAR1-S126, the ADAR1 variant used in the following text) results in the best expression and protein quality after purification (Figures S1A-S1B). Consistent with previous studies^38,39^, size-exclusion chromatography analysis confirmed that the purified ADAR1 assembles into homodimers in the absence of RNA (Figure S1C). To examine the deaminase activity of ADAR1, we developed a coupled enzymatic assay employing *Escherichia coli* Endonuclease V (EcEndoV), which specifically recognizes the deaminase product, “I”, and efficiently cleaves the RNA at the 3ʹ-side of “I”^40^. With a designed hairpin RNA substrate containing a single editable A, the cleaved product can only be observed in the presence of both EcEndoV and ADAR1, but not with either enzyme alone (Figure 1A).

**Figure 1.**
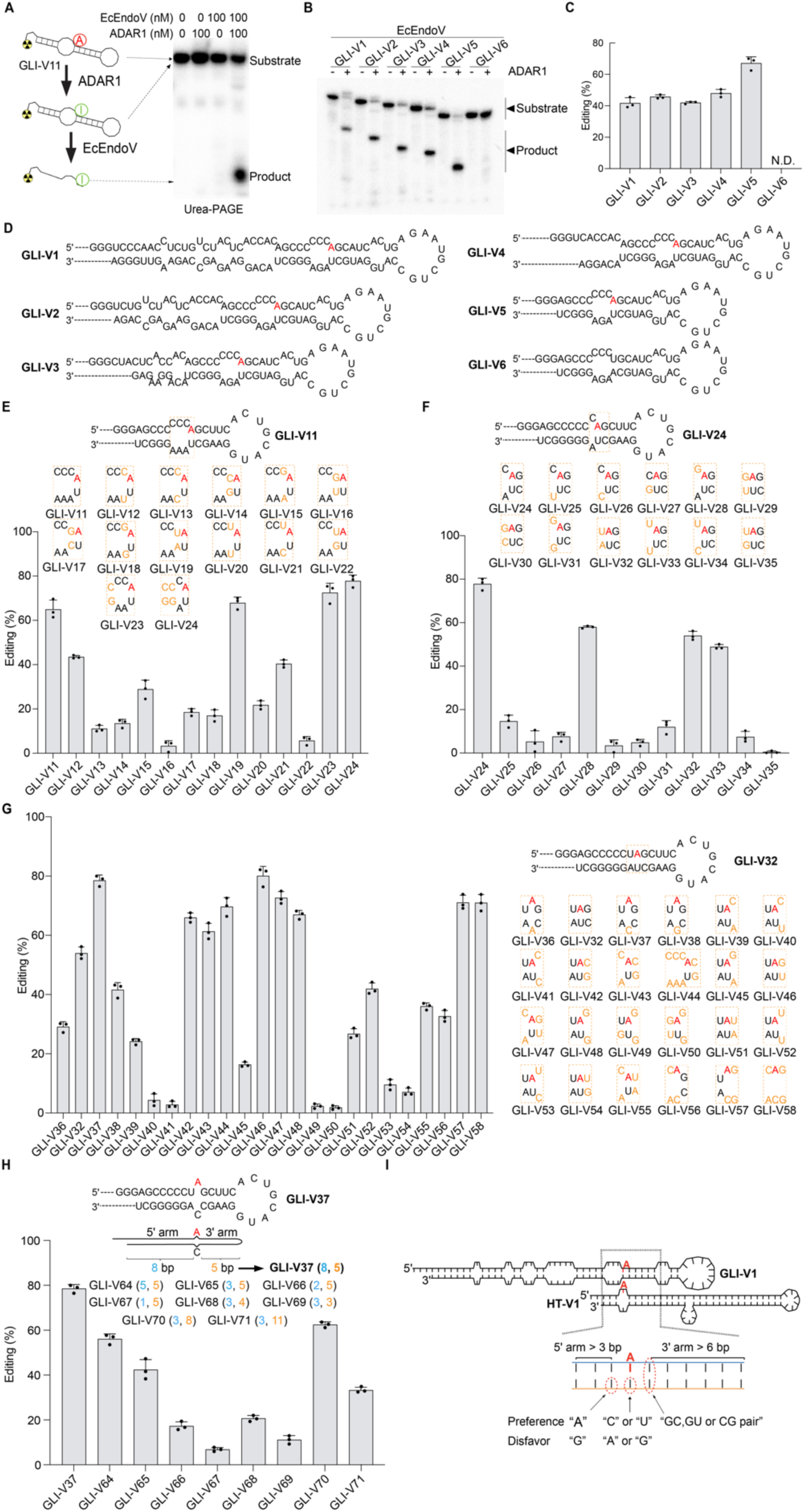
Biochemical profiling of ADAR1 RNA substrate preference on GLI RNA variants. (A) Diagram illustrating the coupled enzymatic assay utilized for ADAR1 activity assessment. Bands of substrate and cleavage products are indicated. The RNA diagram represents the secondary structure of the GLI-V11 RNA used in our experiments. (B) Representative gels showing RNA editing activities of ADAR1 against GLI RNA variants (V1-V6) (C) Normalized editing activity of (B). N.D. indicates not determined. (D) The secondary structures of RNA variants were tested in (B) with the editable A highlighted in red. (E-H) ADAR1 editing activity on GLI RNA variants with diverse sequences and lengths. In (E-G), RNA variant backbones are shown as secondary structures. The editable A sites are highlighted in red, while mutated bases are enclosed in dashed boxes and labeled in orange. In (H), lengths of the 5ʹ or 3ʹ arms of the RNA backbones are indicated, with mutated RNA variants of different lengths denoted in parentheses. (I) Summary of the RNA sequence and length preference for ADAR1 editing. The secondary structures and folds of the GLI-V1 and HT-V1 RNAs were displayed and followed by the summary of RNA sequence and length preferences for ADAR1 editing. The editing site A is highlighted in red. Optimal editing lengths for the 5ʹ or 3ʹ arm and sequence preferences surrounding editing sites are indicated. Values represent the mean of three independent experiments, with error bars indicating standard deviation. 120 nM ADAR1 or mutants and 25 nM each RNA were used. See also Figures S1 and S2.

We next systematically investigated the ADAR1 editing with RNA variants derived from Glioma-associated oncogene 1 (GLI) mRNA^41^. ADAR1 can efficiently edit the GLI (GLI-V1) substrate at a primary “A” site on the 5ʹ-strand (Figures 1B-1D). We then constructed the following modifications of GLI-V1, including sequentially shortening the GLI-V1 on the distal end of the hairpin loop (GLI-V2 to GLI-V5), deleting the smaller hairpins (GLI-V7), shortening the larger hairpins (GLI-V8 and GLI-V9), changing the 3-bp bubble from “CCC:AGA” to “CCC:AAA” (GLI-V10), or only keeping the “A” site after a 3-bp bubble on duplex (GLI-V11), and found none of those affected ADAR1 editing efficiency (Figures 1B-1D, and S1D-S1F). As a control, mutating the editable “A” site on GLI-V6 eliminated the major editing band (Figures 1B-1D). We then systematically tested the RNA sequence preference at or near the editable “A” site based on GLI-V11 (Figure 1E). On the 5ʹ-adjacent pair, the non-editing strand has a strong preference for “A”, while “G” is disfavored on either the editing or non-editing strands (editing efficiency: C:A>C:U>C:C or C:G; G:A>G:C or G:G>G:U; U:A>U:C>U:U>U:G, with RNA substrates GLI-V11 to V22; Figures 1E and S1G). Besides, having three, two, or one mismatches on the 5ʹ-side of the editable A shows similar editing efficiency (GLI-V11, V23, and V24; Figures 1E and S1G). The 5ʹ-adjacent pair preference was further confirmed with the RNA variants based on GLI-V24, which only contains one mispair at the 5ʹ-adjacent pair (GLI-V24 to V35; Figures 1F and S1G). At the editing site, the base opposing the editable A (orphan base) has a preference for “C” (GLI-V37), followed by “U” (GLI-V32), but disfavors “A” or “G” (GLI-V36 and V38) (Figures 1G and S1G). On the 3ʹ-adjacent pair, RNAs with C:G (GLI-V42, V43, and V44), G:C (GLI-V32), or G:U (GLI-V46 and V47) base pair were more efficiently edited than the U:A (GLI-V51 and V55) or U:G (GLI-V54) base pair, but no clear preference was found for different combinations of mismatches (GLI-V39 to V55; Figures 1G and S1G). Furthermore, RNA substrates having the editable A surrounded by two or three mismatches but with preferred 5ʹ and 3ʹ-adjacent sequences still retained good editing activity (GLI-V56 to V58; Figures 1G and S1G). In addition, we designed GLI-V59 to GLI-V63 by introducing C:U mismatches away from the editing site on either the 5ʹ or 3ʹ arm based on the backbone of GLI-V32, which lacks mismatches in the duplex RNA (Figure S1J). Interestingly, introducing a mismatch 2 base pairs away from the editing site on the 3ʹ arm (GLI-V62) resulted in a moderate decrease in editing activity compared to the perfect duplex (Figures S1H-S1J). We next investigated the length preference of RNA editing based on GLI-V37’s backbone (Figure 1H). With the 3ʹ-arm fixed at 5 bp, shortening the 5ʹ-arm of RNAs from 8 bp to 1 bp sequentially decreased editing activity (GLI-V37 and V64 to V67), and the most significant activity reduction occurred when the 5ʹ-arm was reduced from 3 bp (GLI-V65, 42% editing) to 2 bp (GLI-V66, 16% editing) (Figures 1H and S1K). With the 5ʹ-arm fixed at 3 bp, the GLI-V70 with 8 bp 3ʹ-arm was more efficiently edited than those with 11, 5, 4, or 3 bp (GLI-V65, V68 to V71; Figures 1H and S1K).

To validate the RNA specificity for ADAR1 editing, we selected another physiologically relevant RNA substrate but with distinct sequence and structural features, a 71 nucleotide RNA derived from 5-HT_2C_R (HT) pre-mRNA (HT-V1)^7^. ADAR1 efficiently edits the HT-V1 substrate as well as the HT-V2, which replaced the other A:U pairs near the editable A (Figures S2A-S2C). Further deleting 5 nt of the bulge (HT-V3) and replacing all A:U pairs but only keeping one editable A on the editing strand (HT-V4) retained ADAR1 editing (Figures S2A-S2C). We next verified the RNA length requirement of HT for ADAR1 editing. On the 5ʹ-arm, reducing the length from 3 (HT-V6) to 2 or 1 bp (HT-V13 or V12) almost abolished the editing, whereas increasing the length to 6, 9, or 20 bp (HT-V14 to V16) retained editing activity (Figures S2A-S2C). On the 3ʹ-arm, reducing the duplex length from 25 to 6 bp (HT-V4 to HT-V8) retained ADAR1 editing, but the editing activity was dramatically reduced when the 3ʹ-arm was shortened from 6 bp (HT-V8, 60 % editing) to 5, 4, or 3 bp (HT-V9 to V11, less than 20 % editing) (Figures S2A-S2C). These results confirmed that the ADAR1 editing is length-dependent, requiring at least 3 bp on the 5ʹ-arm and 6 bp on the 3ʹ-arm for optimal editing. We next investigated the sequence preference for editing based on HT-V6. At the editing site, an A:C mismatch is preferred over an A:U base pair, whereas an A:A or A:G mismatch significantly reduced the activity (HT-V6 and V24 to V26; Figures S2D and S2E). On the 5ʹ-adjacent pair, a U:A base pair was preferred over a G:U, U:G, G:C, or C:G base pair, again suggesting the preference for “A” on the non-editing strand (HT-V6 and V17 to V20; Figures S2D and S2E). On the 3ʹ-adjacent pair, a G:C base pair was preferred over G:U, U:A, or U:G (HT-V6 and V21 to V23; Figures S2D and S2E). Our results confirmed that HT RNA shares a similar RNA sequence and structural preference as GLI RNA (Figure 1I), even though the two RNA substrates appear to have quite different sequences and structures.

### Cryo-EM structures of ADAR1-RNA complex

We next employed cryo-electron microscopy (cryo-EM) to investigate the structural basis of ADAR1 editing. Two distinct RNAs with different sequence and structural features, the GLI-V11 and HT-V2 (Figures 1B and S2B), were selected for cryo-EM studies. To stabilize the ADAR1-RNA complex, a transition state mimic analog 8-azanebularine (8-aza)^42^ was introduced to the editable A site in both RNAs. With cryo-EM single particle reconstruction (Figures S3 and S4), the ADAR1-GLI-V11 structure was determined at 3.01 Å (ADAR1-GLI Complex) (Figures 2A, 2C, S3, and S5, and Table S1) and the ADAR1-HT-V2 structure was determined at 3.20 Å (ADAR1-HT Complex) (Figures 2B, 2D, S4, S5, and Table S1). In each cryo-EM map, a deaminase domain dimer and its associated RNA, 14 bp for GLI and 29 bp for HT, can be well traced (Figures S6A-S6J). However, although a nearly full-length ADAR1 was used, none of the ZBDs or RBDs were observed in electron density maps, possibly due to their high flexibility. In both GLI and HT structures, the deaminase domains (residues 840-1224) form an asymmetric dimer (monomers A and B), with monomer A mainly interacting with RNA (Figure 2). The overall conformations of the two monomers are similar with an averaged root-mean-square deviation (RMSD) of 0.558 Å for the 305 aligned Cα atoms, however, two active site loops (residues 1003-1010 and 1108-1124) in monomer A take different conformations in monomer B, and a loop at the dimer interface (residues 1016-1025) in monomer A is disordered in monomer B (Figures S7A). Moreover, the overall structure of ADAR1-RNA and ADAR2-RNA complexes^19^ (PDB 8E0F) aligns well with a RMSD of 1.5 Å for the 531 aligned Cα atoms (Figure S7B). Interestingly, a flexible loop in ADAR1 (residues 970-995) assumes a distinct conformation in ADAR2 (residues 454-476). Residue H988 in the loop forms a new zinc-binding site together with C1081, C1082, and H1103 (Figures S6D and S7B), consistent with previous predictions^43^. In addition, an inositol hexakisphosphate (IHP) molecule, which was reported to help protein folding of ADAR2^42,44^, was also found in each ADAR1 monomer, where they form numerous hydrogen bonds with conserved residues^44^ (Figures S6E-S6F).

**Figure 2.**
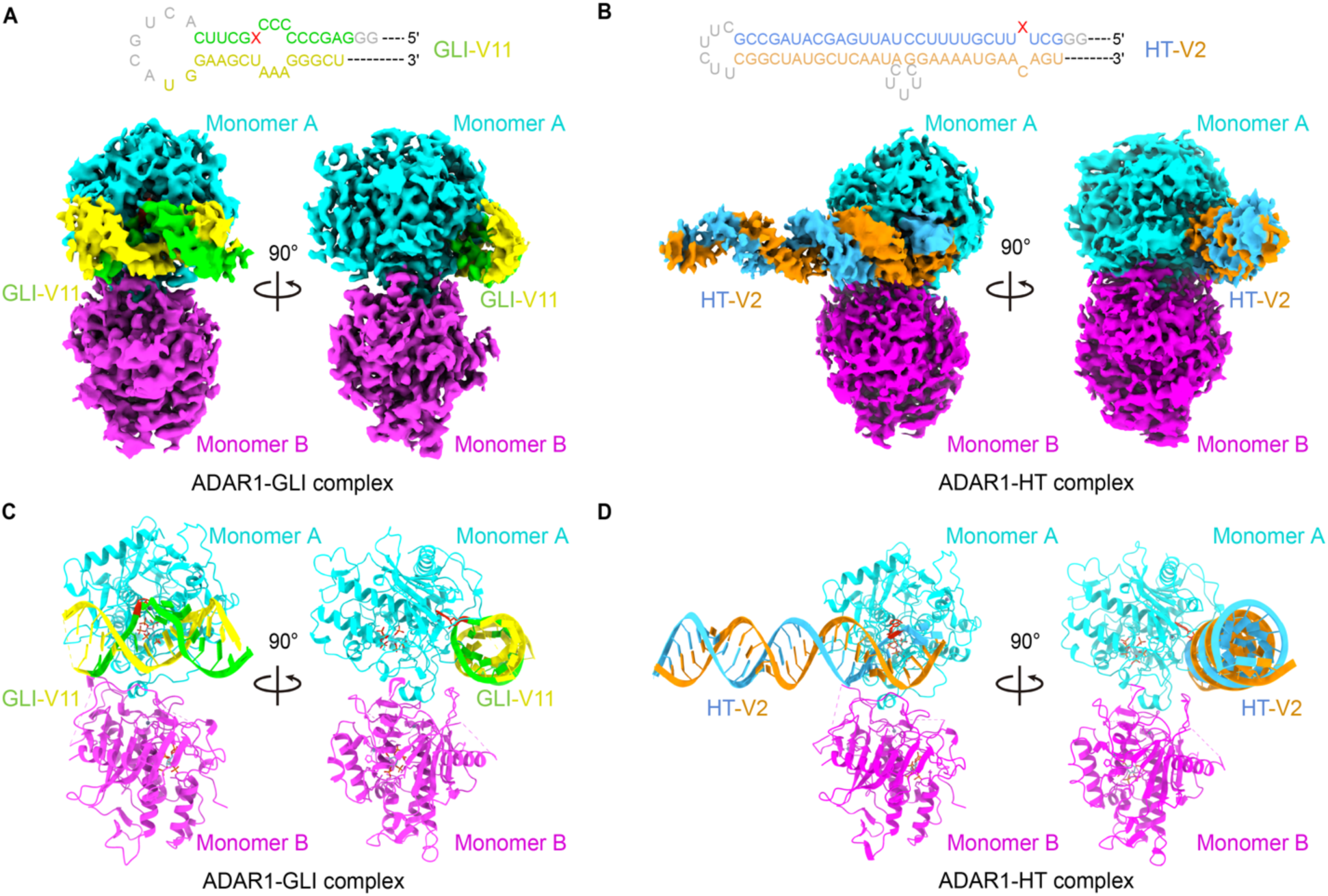
Cryo-EM structures of ADAR1 complexes with GLI or HT RNA. (A-B) Two views of the cryo-EM maps of the ADAR1-GLI (A) or ADAR1-HT (B) complex. The secondary structure of GLI-V11 or HT-V2 RNA is displayed in the top panel of (A) or (B), the modified A (8-aza) in GLI-V11 or HT-V2 RNA is denoted as "X" and highlighted in red. The editing strand of GLI-V11 or HT-V2 is colored in green or blue, respectively, while the non-editing strand of each is colored in yellow and orange, the disorder sequences in GLI-V11 or HT-V2 are colored in grey. Monomer A is colored in cyan, and monomer B is colored in magenta. (C-D) Two views of the atomic model of the ADAR1-GLI complex (C) or ADAR1-HT complex (D). The models are color-coded as in (A) and (B). See also Figures S3-S7 and Tabel S1.

### The structural basis for ADAR1 RNA binding and editing

The two high-resolution structures of the ADAR1-RNA complexes provide a great opportunity for understanding the structural basis of ADAR1 RNA binding and editing. The GLI and HT dsRNA bind in a similar conformation to ADAR1, with an RMSD of 0.955

Å for the 709 aligned Cα atoms (Figures 3A, S7C, and S7D). Consistent with the length dependence identified by biochemical assays, ADAR1 contacts the dsRNA with 3-4 bps duplex on the 5ʹ arm and 6-7 bps duplex on the 3ʹ arm on both RNAs (Figures 3B-3E). Interestingly, both the 3-bp "CCC:AAA" bubble on the 5’ arm of the GLI RNA and the short 3-bp “GCU:UGA” 5’ arm duplex of the HT RNA are stabilized by ADAR1 with distorted base pair patterns that are distinct from regular A-form dsRNA (Figures 3D, 3E, S7E and S7F). Our observations suggest that ADAR1 can tolerate RNA substrates with mismatches by remodeling the RNA structures around the editing site, which is consistent with the biochemical results (Figures 1E and 1G). In both ADAR1 structures, the R892, K895, K996, K999, R1001, R1030, and K1120 from monomer A and K1115 from monomer B are proximate to the RNA backbones (Figures 3B-3E). Consistent with the structural observation, mutating the RNA interaction residues (K895E, K996E, R1001E, R1030E, K1115E, and K1120E) significantly decreased the editing activity for GLI-V11 (Figures 3F, and S10A). However, only R1001E, R1030E, and K1120E show decreased activity for the longer HT-V2 RNA, whereas K895E, K996E, and K1115E retain similar levels of HT-V2 editing as WT (Figures 3F and S10D). Further examination revealed that all these mutants show editing defects for RNA (GLI-V11, V32, or HT-V6) with short (<10 bp) 5ʹ and 3ʹ arms, or RNA with a long (20 bp) 5ʹ-arm (HT-V16) (Figures 3F, S10A-S10C, and S10F). However, for the RNA with a long (20 bp) 3ʹ-arm (HT-V5), the K895E, K996E, and K1115E ADAR1 retain comparable activity as WT, similar as editing against HT-V2 RNA (Figures 3F, S10D, and S10E).

**Figure 3.**
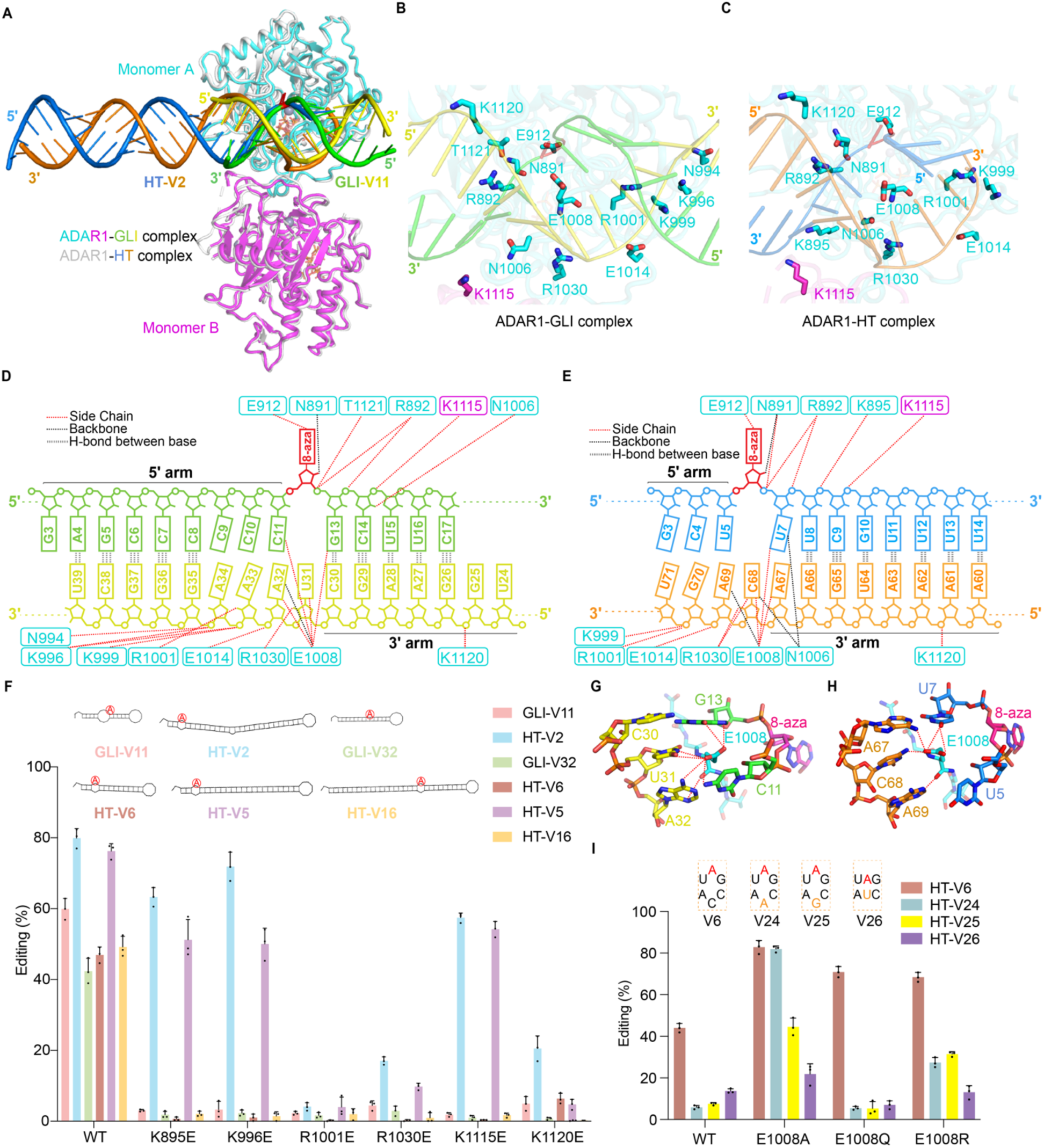
Structural insights into ADAR1 editing. (A) Structural comparison of the ADAR1-GLI and ADAR1-HT complex, with the monomer A aligned. The ADAR1-GLI complex and HT-V2 RNA in the ADAR1-HT complex are color-coded as in Figures 2C and 2D, respectively. The ADAR1 dimer of the ADAR1-HT complex was colored in grey. (B-C) The zoom-in views of the RNA recognition site in ADAR1-GLI (B) and ADAR1-HT complexes (C). (D-E) Overview of the molecular interactions between ADAR1 and GLI-V11(D) and HT-V2 (E) RNA. (I) (F) Editing activity of ADAR1 wild type (WT) or mutants with different RNA substrates. Secondary structures of RNA substrates are displayed, with the editing site A highlighted in red. (G-H) Nucleotide recognition by E1008 in the ADAR1-GLI complex (G) and the ADAR1-HT complex (H). (I) Editing activity of ADAR1 E1008 mutants with four HT-RNA variants with different sequence contexts around the editing site. (F) and (I) Values represent the mean of three independent experiments, with error bars indicating standard deviation. 100 nM ADAR1 or mutants and 25 nM each RNA were used. See also Figures S6-S10.

In both ADAR1-GLI and ADAR1-HT complexes, the modified editable A (8-aza) is flipped out of the RNA duplex and inserted into the active site pocket (Figures 3D, 3E, and S8A). The O6 group of 8-aza and the side chains of C966, C1036, and H910 coordinate the catalytic Zn^2+^ and hydrogen bonds with E912’s side chain (Figure S8A). Compared to monomer B, the conformation changes of the flexible loop in monomer A (residues 1003-1010) promoted the 8-aza flipping, resulting in a 6.7 Å shift of E1008 (Figure S7A). The E1008 in monomer A is inserted into the dsRNA minor groove to take the place of the flipped 8-aza (Figures 3G and 3H), forming hydrogen bonds with the orphan base “U31” in ADAR1-GLI or “C68” in the ADAR1-HT complex (Figures 3G and 3H). Consistent with the biochemical preference (Figure 1I), modelling of “A” or “G” at the orphan base leads to a steric clash with E1008 (Figures S8B and S8C). In addition to interacting with the orphan base, E1008 plays a pivotal role in recognizing the non-editing strand. Structural analyses reveal that E1008 interacts with the purine base (adenine) on the non-editing strand (“A32” in ADAR1-GLI or “A69” in ADAR1-HT) through hydrogen bonds formed by its side chain or main chain (Figures 3G and 3H). Replacing the 5ʹ-adjacent “A” on the non-editing strand with a smaller “C” or “U” would abolish the hydrogen bond interaction with E1008, while replacing it with “G” may cause a steric clash with the main chain of E1008 in both structures (Figures S8D and S8E). These findings further indicate that the strong sequence preference for the non-editing strand is largely mediated by E1008. To confirm the role of conserved E1008 in ADAR1 substrate selection, we created E1008A, E1008Q, and E1008R mutations and tested them against HT-RNA variants. Interestingly, all three mutants show better activity for HT-V6 (with A-C mismatch at the editing site) than the WT (Figures 3I, S10G, and S10H), similar as previously reported^18,45,46^. Furthermore, E1008A showed efficient editing regardless of the orphan base sequence, confirming its role in orphan base selection (Figures 3I, S10G, and S10H). However, WT and E1008 mutated ADAR1 displayed similar levels of editing to HT variants with non-preferred 5ʹ adjacent pairs (HT-V17 and HT-V18) (Figures S10I and S10J), possibly because the sequence preference at this position is mainly controlled by steric clashes between non-preferred 5ʹ adjacent pairs and the main chain of E1008.

The ADAR1 asymmetric dimer is mainly mediated by a conserved helix^19^ (residues 1021–1030) in monomer A and the catalytic pocket (residues 1001-1010) in monomer B (Figures S9A and S9B), with a surface area of 1115 Å^2^. Specifically, two residues in monomer A are important for dimerization: D1023 is inserted into the interface cavity of monomer B and interacts with R1001, G1009, and T1010 through a salt bridge and several hydrogen bonds; and the bulky W1022 sidechain interacts with A970 and L971 from monomer B with hydrogen bond and hydrophobic interactions (Figures S9A and S9B). Similar to our RNA binding site mutations and as previously reported for ADAR2^19^, the W1022A mutation affected the editing of short RNAs (GLI-V11, V32, or HT-V6) or RNA with a long 5ʹ arm (HT-V16), but not RNAs with a long 3ʹ arm (HT-V2 and HT-V5), whereas the D1023A mutation significantly decreased the editing activity on all RNAs (Figures S9C and S10A-S10F).

### The molecular basis of AGS-associated mutations on ADAR1

Eight ADAR1 mutations have been correlated with AGS, with seven of these mapped to the deaminase domain (A870T, L872T, R892H, K999N, G1007R, Y1112F, and D1113H), and one (P193A) residing in the ZBD-α domain^4^. Previous studies have suggested that AGS mutations on ADAR1 would affect ADAR1 RNA editing^4,47–50^, yet their substrate preferences are not well defined. In the ADAR1-GLI complex, R892 and K999 are directly involved in the dsRNA binding, and G1007 flanks E1008 to displace the editing “A” (Figure 4A). Y1112 and D1113 are located on the flexible loop (residues 1108-1124) near residues for RNA binding (K1115) and asymmetric dimerization (R1116-S1118) (Figures 4A and S9B). Lastly, A870 and L872 are located in the protein interiors, possibly important for protein folding and stability (Figure 4A). Interestingly, the RNA-binding affinities of all AGS mutants are comparable to that of the WT (Figure S11), and size-exclusion chromatography analysis indicated all purified AGS mutants eluted as homodimers with similar retention volumes as WT (Figure S16F). However, our biochemical profiling suggested that the AGS mutations can be classified into three categories: G1007R eliminates ADAR1 editing on all RNA substrates; P193A, which has been shown to affect ADAR1 binding to Z-RNA^51,52^, retains ADAR1 editing activity on our model RNAs; A870T, L872T, R892H, K999N, Y1112F, and D1113H have significantly reduced activity against short GLI and HT RNAs, but maintain comparable activity towards HT-V2 and HT-V5 with long 3ʹ arm (Figures 4B, and S10K-S10P). Moreover, Y1112F and D1113H also maintained ADAR1 activity on HT-V16 with long 5ʹ arm dsRNA (Figures 4B and S10K-S10P). Thus, our results suggested that AGS mutations on ADAR1 may affect different properties of ADAR1 binding and editing, and most of these mutations have specific defects in editing short duplex RNAs.

**Figure 4.**
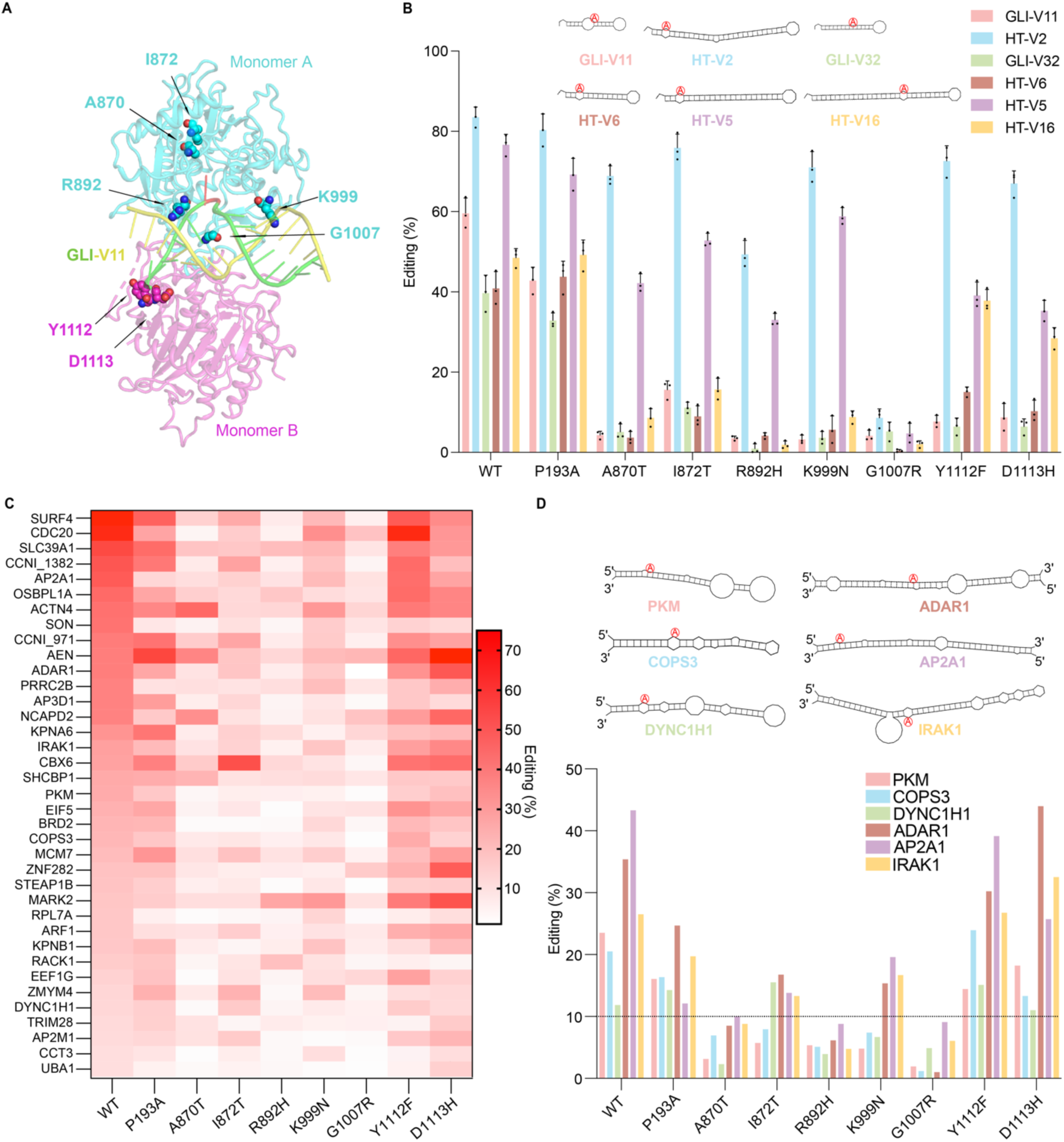
Structural, biochemical characterization and RNA-Seq analysis of ADAR1 AGS-associated mutations. (A) Mapping of AGS-associated mutations onto the ADAR1-GLI complex. (B) Editing activity of ADAR1 AGS-associated mutants with different RNA substrates. Values represent the mean of three independent experiments, with error bars indicating standard deviation. 100 nM ADAR1 or mutants and 25 nM each RNA were used. (C) Heatmap of RNA editing rates for WT and AGS-associated ADAR1 mutants across 37 selected RNAs, as determined by RNA-Seq. The name, for example, CCNI_971, indicates editing at position A971 in the CCNI mRNA. Among the tested groups, the G1007R mutant shows the most severe defects in RNA editing. (D) Editing activity of wild-type (WT) ADAR1 and AGS-associated mutants on selected dsRNAs identified from RNA-Seq data. Secondary structures of the RNAs are shown, with the editing site adenosine (A) highlighted in red. The black dotted line indicates 10% editing activity as a reference. See also Figures S10–S13.

To validate the impaired RNA editing activities of ADAR1 mutants, we performed RNA-Seq in HEK293 cells overexpressing ADAR1 WT, AGS mutants, or an empty vector control. Expression of all ADAR1 variants was confirmed by their mRNA levels (Figure S13B). Whole-transcriptome analysis showed that P193A, Y1112F, and D1113H exhibited a mild reduction in overall editing compared to WT ADAR1, whereas A870T, R892H, K999N, and G1007R exhibited significant decreases in the number of editing sites (Figure S13A; Table S5). To minimize background editing events from endogenous ADAR1 and/or ADAR2 expression (Figure S13B), we selected editing sites with at least ten reads across all groups and retained those sites with editing rates exceeding 10% in WT but no editing in the control. Applying these criteria, we identified 37 RNA substrates (Figure 4C; Table S6). The editing trends aligned with our biochemical results: G1007R displayed the most severe editing impairment, while A870T, I872T, R892H, and K999N showed significant editing reductions across most RNA substrates. In contrast, mutants P193A, Y1112F, and D1113H behaved similarly to WT (Figure 4C).

To investigate the substrate preferences of AGS mutants, the secondary structures of all selected RNAs were analyzed (Figures 4D and S13C–S13F) using the RNAfold serve^53^. Of these, 17 RNAs were excluded as their editing sites were predicted to exist outside of the duplex RNA regions, on large bulges or with branches (Figures S13C and S13D), likely due to inaccuracies in secondary structure predictions, unidentified long-range interactions or unknown complementary RNA sequences^54,55^. For the remaining RNAs, the P193A, Y1112F, and D1113H mutants showed editing activities similar to those of WT (Figures 4D and S13E). Moreover, the A870T, I872T, R892H, and K999N mutants displayed more pronounced defects in editing RNAs forming short dsRNA structures with multiple mismatches and/or large bulges (e.g., PKM, COPS3, or DYNC1H1), while being less affected for RNAs forming long dsRNA structures (e.g., ADAR1, AP2A1, or IRAK1) (Figure 4D). While most RNAs followed the general trend observed in our biochemical results, the G1007R mutant exhibited high background editing (>10%) on EEF1G, CDC20, ZNF282, NCAPD2, and AEN RNAs (Figure S13F), and the short dsRNAs (SHCBP1, ZMYM4, OSBPL1A, or CBX6) showed deviations in editing preference for two or three mutants (A870T, I872T, R892H, or K999N) (Figure S13E). This variability may be attributed to inaccuracies in secondary structure predictions, unknown factors affecting ADAR1 recruitment and editing efficiency, and/or editing associated with endogenous ADAR1/ADAR2 expressions. Overall, our RNA-seq data confirmed that AGS mutants exhibit varied defects in RNA editing activity, with particularly at editing sites located in short dsRNA regions.

It has been reported that the HT-RNA is specifically expressed in the brain^7^, while GLI-RNA is expressed in HEK293 cells^55^. However, the low expression levels of GLI RNA led to insufficient reads for reliable analysis (Figure S13B) in our transcriptome-wide profiling. To address this, we amplified GLI cDNA and performed Sanger sequencing. Our results identified four editing sites on the GLI duplex RNA (Figures S13E and S13F). Site 2, which was previously reported^41^, exhibited high background editing in the control or G1007R groups, suggesting that it may not be primarily edited by the overexpressed ADAR1. Site 1, the major editing site in our in vitro assay, along with site 3, both of which are located near the end of the RNA duplex, showed the most impaired editing in mutants A870T, R892H, K999N, and G1007R. Site 4, located in the middle of the duplex RNA, retained comparable editing activity for all ADAR1 variants except G1007R (Figures S13E and S13F). Overall, these results confirm the G1007R as the most defective mutation, while other mutants display varying levels of editing deficiencies depending on the RNA secondary structures.

### The role of RBDs in ADAR1 editing

Apart from the deaminase domain, the ZBD and RBD domains on ADAR1 may also contribute to RNA binding and editing. To test their roles, we sequentially removed the ZBD and RBD domains from the N-terminus of ADAR1 (Figure 5A). Interestingly, the editing activity on three RNAs with different arms’ lengths (HT-V5, V6, and V16) was kept at a similar level by removing ZBD-α (P110), ZBD-β (R1R2R3D), RBD1 (R2R3D) or RBD2 (R3D), indicating that these domains may only have minor roles in editing our model RNA substrates (Figures 5B, and S14A-S14C). However, the RNA binding and editing activity were both significantly reduced when only the deaminase domain (D-801 or D-833) was retained, and the editing product can only be confidently detected with higher protein concentration (to 1 μM) for both D-801 and D-833 (Figures 5B, and S14A-S14G). Furthermore, the R3D bearing the dsRNA binding deficient mutations on RBD3^56^ (R3D-EAA; containing K777E/K778A/K781A mutations) also abolished RNA binding and editing activity (Figures 5B, S14H, and S14K), while R3D maintained the dsRNA binding ability and showed a better binding for the RNA with a long 3’ arm (HT-V5) than the short HT RNA (HT-V6) or RNA with a long 5’ arm (HT-V16) (Figures S14I and S14J). These results align with previous findings that the RNA binding of RBD3 is crucial for ADAR1 editing^56,57^, but distinct from studies of ADAR2, where the deaminase domain alone is sufficient for editing^42^.

**Figure 5.**
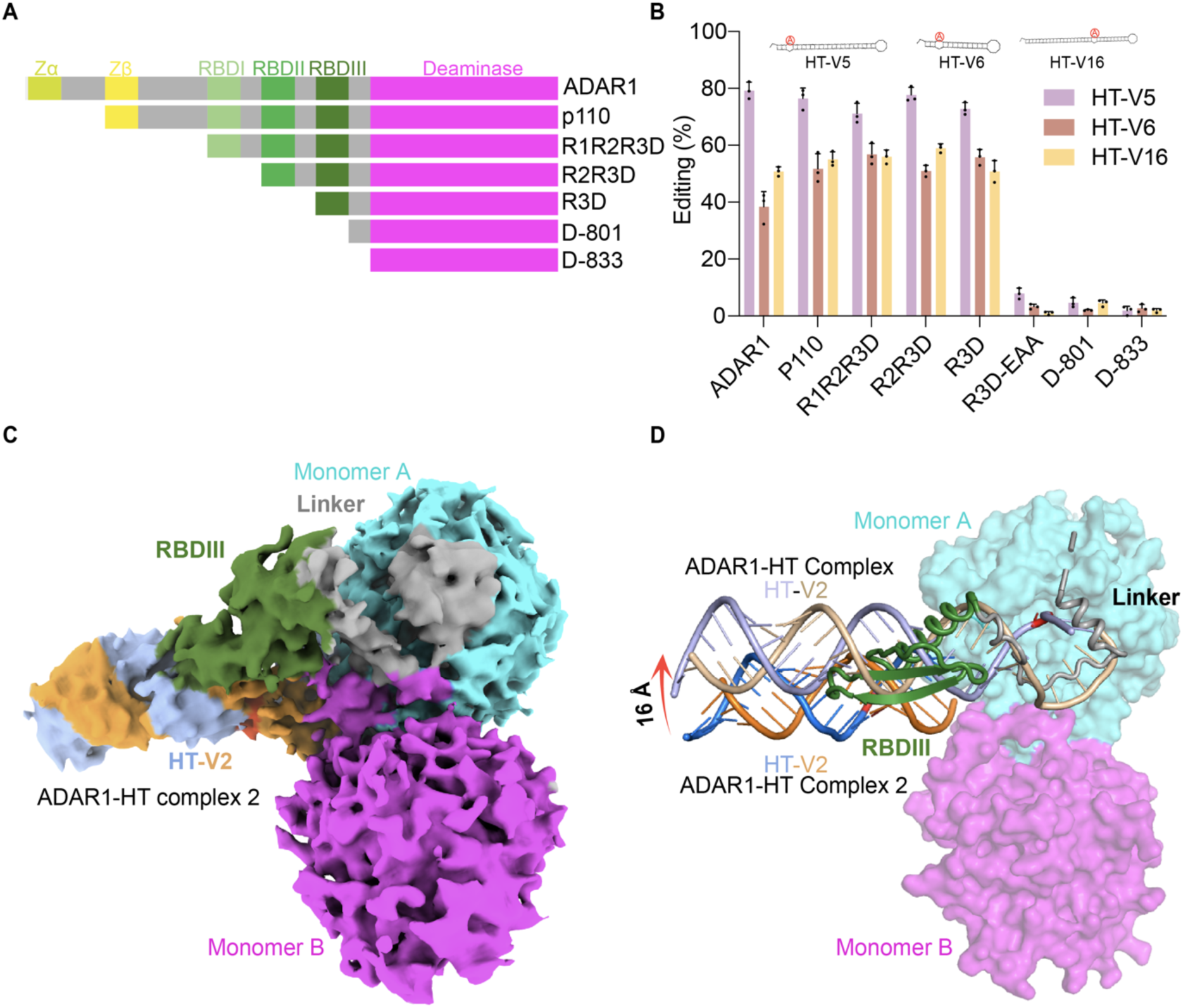
Biochemical characterization of ADAR1 truncation mutations. (A) Diagram of ADAR1 truncation mutants. (B) Editing activity of ADAR1 truncation mutants against three HT RNA variants. Secondary structures of RNA substrates are indicated on the top panel, with the editing site A highlighted in red. Values represent the mean of three independent experiments, with error bars representing standard deviation. 100 nM ADAR1 or truncation mutants and 25 nM each RNA were used. (C) The cryo-EM map of the ADAR1-HT complex 2. The structure is color-coded similarly as in Figure 2. (D) Structural comparison of the HT-RNA in ADAR1-HT complex and ADAR1-HT complex 2. Monomer A from the two structures is aligned. In the ADAR1-HT complex, only the HT-RNA is shown for clarity, and colored in light blue (editing strand) and wheat (non-editing strand), respectively. The movement of HT-RNA is indicated with a red arrow. See also Figures S14-S15.

To identify possible interactions between RBD3 and dsRNA, we performed local 3D classification with a focused mask on RBD3 and identified a new ADAR1-HT Complex 2 at 3.83 Å (ADAR1-HT Complex 2, Figures 5C, 5D, S4, S5G-S5I, and Table S1). The RBD3 (residues A742-G794) and the linker (residues E795-S823) from monomer A, as well as 17 bp of HT-V2 dsRNA can be rigid-body docked to the density, although the local cryo-EM map doesn’t permit atomic model building of RBD3 (Figures 5C, 5D, and S6K-S6M). The RBD3 in the ADAR1-HT Complex 2 is located near the monomer A, distinct from the RBD2 in the ADAR2 structure, which is derived from monomer B and resides at the asymmetric dimer interface^19^ (Figure S15A). Compared to the ADAR1-HT complex, the dsRNA in the ADAR1-HT Complex 2 undergoes a large movement (16 Å) away from the active site (Figure 5D), with the linker between RBD3 and deaminase domain partially blocking the RNA binding site for editing. The ADAR1-HT Complex 2 may represent a pre-editing state, and conformational changes of both RBD3 and the linker are required to permit the dsRNA binding to the active site for efficient editing.

## Discussion

Adenosine deamination represents one of the most abundant post-transcriptional modifications of eukaryotic RNA^58,59^. However, the sequence and structural features of RNA for efficient ADAR1 editing have not been well defined. In this study, we have established a well-controlled biochemical system for comprehensively evaluating ADAR1 substrate preference. Through examining nearly a hundred synthetic RNA substrates with sequential variations of either editing or non-editing strands, we found that the ADAR1 editing selectivity is determined by both strands, with a strong sequence preference for the non-editing strand (Figure 1). These findings expand our understanding of the 5′ and 3′ neighboring sequence preference, which was previously thought to be mainly determined by the editing strands^31–33,36^. In addition, different from ADAR2, which requires an RNA with a long 5’ arm (>11 base pairs) for binding and editing^42^, ADAR1 can efficiently edit RNA with a minimum length of only ∼10 bp RNA duplex, with 3 bp on the 5ʹ arm and 6 bp on the 3ʹ arm. This observation is consistent with our structures that ADAR1 interacts with the dsRNA by contacting a 3-4 base pair duplex on the 5ʹ arm and a 6-7 base pair duplex on the 3ʹ arm in both RNA complexes (Figures 3D and 3E). Moreover, our biochemical data suggested that ADAR1 has a remarkable tolerance for mismatches near the editing site (Figures 1E and 1G), consistent with structural evidence showing that ADAR1 remodels the RNA around the editing site to accommodate these mismatches (Figures S7E, and S7F). This minimal length requirement and mismatch tolerance may explain the widespread ADAR1 editing on endogenous RNA. Moreover, consistent with biochemical analysis, ADAR1 takes the same mode in editing the GLI and HT RNAs, which have different lengths, local sequences, and base pairing patterns (Figure 1I). Our data thus provided clues for the effective mining of ADAR1 substrate and application of ADAR1 RNA editing for biotechnological applications^14^.

ADAR1 contains six functional domains, which all possibly bind duplex RNA. In our ADAR1 structures, the deaminase domain forms an asymmetric dimer with residues from both monomers interacting with dsRNA (Figures 3B-3E). Disrupting the asymmetric dimerization interface abolished ADAR1 editing, suggesting the importance of the asymmetric dimer for RNA binding and editing (Figure S9). Interestingly, although ADAR1 and ADAR2 share a conserved dimerization interface and a 59% sequence similarity of the deaminase domain^43^, only residues from monomer A in ADAR2 appear to contribute to dsRNA binding^19,42^. Besides the deaminase domain, our studies also underscore the pivotal role of RBD3 in ADAR1 RNA binding and editing (Figure 5). The structure and biochemical data indicated the RBD3 from monomer A may help first recruit the substrate dsRNA for subsequent editing, as revealed in our ADAR1-HT Complex 2 structure (Figures 5C and 5D). In addition, RBD3 can be docked to the position of RBD2 from monomer B of ADAR2^19^ without steric clashes (Figure S15B), suggesting RBD3 may also have a direct role in promoting ADAR1 editing^56^. Interestingly, previous reports have also indicated that RBD3 is important for ADAR1 dimerization in the absence of RNA^24,38,39^. This might explain why mutants W1022A or D1023A at the deaminase dimer interface migrate similarly as the WT protein in size-exclusion chromatography analysis (Figure S16F). Although our truncations of ADAR1 suggested that RBD1 and RBD2 are not important for ADAR1 editing of our model RNA substrates (Figure 5B), these domains may contribute to substrate recognition and RNA binding on long duplex RNA or Alu RNA^22,60,61^. In addition, deletion or mutation of ZBDs does not alter ADAR1 editing, possibly due to the lack of Z-RNA in our model substrates.

Furthermore, ADAR1 play pivotal regulatory roles in innate immune response, and mutations in ADAR1 have been correlated to human autoimmune disease AGS^2^. It has been reported that ADAR1 editing is essential for antagonizing MDA5 activation by editing 3′-untranslated region (3′-UTR) dsRNAs primarily comprising long inverted Alu RNA repeats^17,50,62^, whereas ADAR1 RNA binding to dsRNA or Z-RNA is important for masking the activation of ZBP1 and PKR^10,63^. Although previous studies of AGS mutations suggested that the mutations may have different effects on ADAR1^4,50^, the exact substrate preference of these mutations is not well defined. Our biochemical characterizations and RNA-seq analysis of AGS mutations suggest that they have varied effects on ADAR1 editing, but most of them have more pronounced defects on short duplex RNA editing (Figure 4). As inferred from our structural analysis, the AGS mutations may affect ADAR1 deaminase domain RNA binding, dimerization, and stability. However, we found that AGS mutations do not have an apparent effect on RNA-binding affinities or homodimerization of ADAR1 (Figures S11 and S16). This is likely because the RNA-binding domains may play a compensatory role by maintaining RNA-binding affinity and protein dimerization (Figures 5B and S14), potentially compensating for the effects of the AGS mutations. On the other hand, the AGS mutants may be defective in properly localizing the RNA substrate onto the deaminase domain for efficient RNA editing. For example, G1007R eliminates ADAR1 editing on all RNA substrates in our biochemical assays. The dominant negative effect of G1007R is possibly due to the steric clash of the G1007R side chain in monomer A with RNA and the R1030 (Figure S12A), as well as the potential conflict between the G1007R sidechain in monomer B and the residues of V1019, T1021, I1025, and E1029 from monomer A at the dimerization interface (Figure S12B). Interestingly, all other mutants on the deaminase domain preferentially influence ADAR1 editing of RNA with a short dsRNA (Figure 4), which is unlikely related to the long duplex RNA or Alu RNA editing. However, the precise physiological implications of this RNA preference by AGS mutations remain elusive, warranting further investigations. Furthermore, the P193A mutation may contribute to ADAR1 binding and editing of Z-RNA, as previously described^51^. In our study, P193A has nearly no effect on ADAR1 editing, possibly due to the lack of Z-RNA in our substrates.

In summary, our comprehensive biochemical and structural characterizations provide a deep understanding of the molecular basis of ADAR1-mediated RNA editing and provide insights for understanding immunodeficiency associated with ADAR1 dysfunction. Furthermore, the high-resolution structures of ADAR1 and the biochemical rule of ADAR1 editing will guide rational drug discovery and bioengineering of ADAR1.

## Acknowledgments

We acknowledge the Welch Foundation C-2033-20200401 and the Cancer Prevention & Research Institute of Texas (CPRIT) Award RR190046 and Rice University Startup fund to Y.G., the National Institutes of Health (NIH) R01ES031511 to Y-L. W., and the NIH R01CA268518 and the CPRIT RP220480 to J.W. We thank Dr. Gaya P. Yadav at the Laboratory for Biomolecular Structure and Dynamics (LBSD) of Texas A&M University and the core facility at Stanford-SLAC Cryo-EM Center for cryoEM data collection. We thank Dr. Yizhi Tao for the critical reading of the manuscript.

## Author contributions

Y.G., Y.-L.W., and J.W. conceived the project. X.D. performed the cloning, protein, RNA purification, biochemical and structural studies. X.D., L.S., M.Z., and R.B. performed RNA-Seq and Sanger sequencing studies. Y.G. and X.D. analyzed the results and drafted the manuscript. All authors contribute to the editing and proofreading of the manuscript.

## Supplemental information

Figure S1-S16 and Table S1-S6

## Supplemental figures and legends

**Figure S1.**
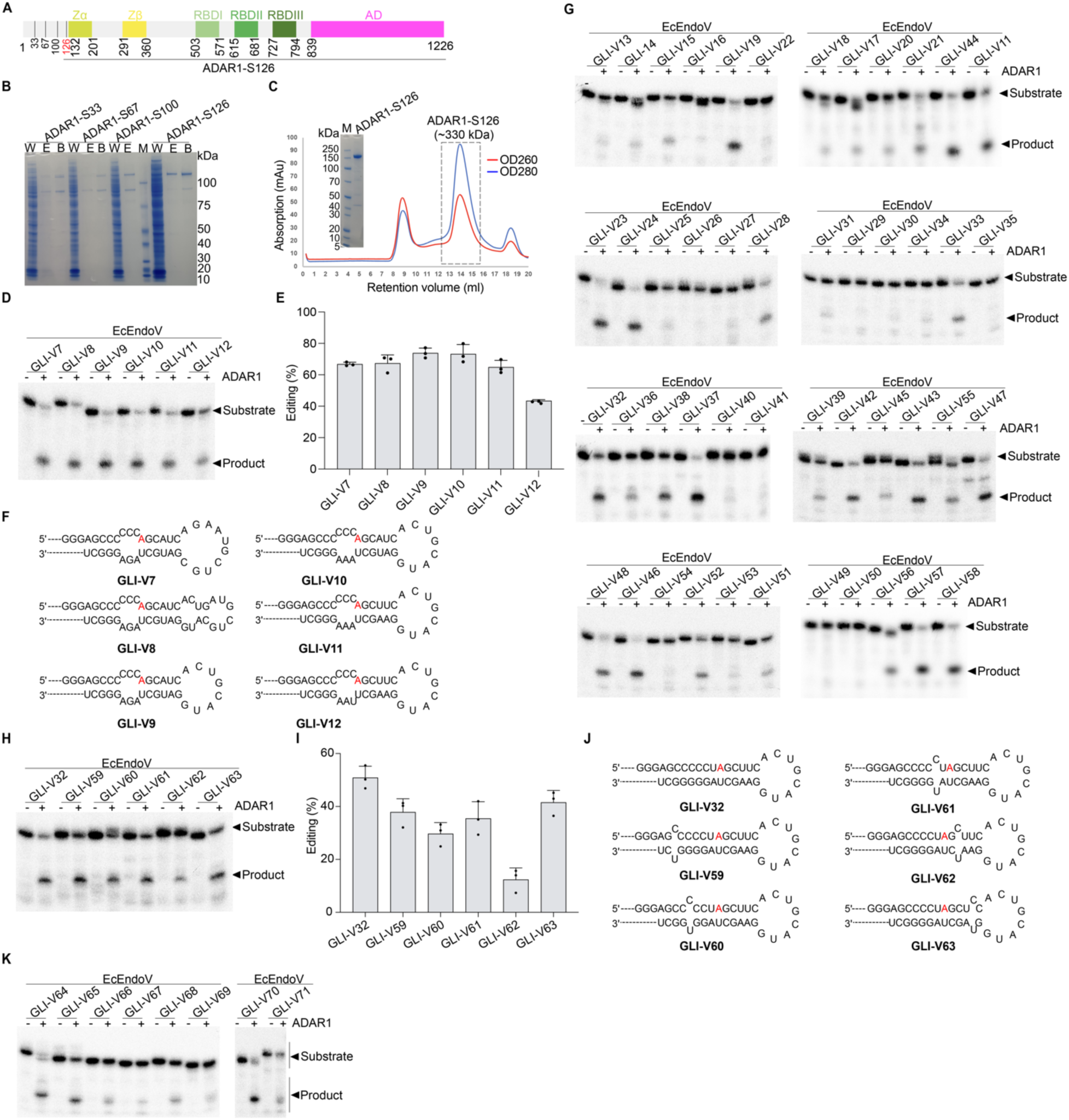
ADAR1 purification and biochemical characterization of ADAR1 editing, related to Figure 1. (A) Domain structures of ADAR1. (B-C) Expression of human ADAR1 N-terminal truncation mutants (B) and purification of ADAR1-S126 indicated by a dashed box in the chromatogram, the estimated molecular weight of MBP tagged ADAR1-S126 homodimer has been indicated, and purity of the protein was checked with SDS-PAGE gel (C). W: Washed, including proteins that do not bind to the amylose resin. E: Elution, including proteins that bind to the amylose resin and can be eluted with the buffer containing maltose. B: Bound, including proteins that bind to the amylose resin but cannot be eluted. (D-F) Representative gels showing RNA editing activities of ADAR1 against GLI RNA variants (V7-V12). (E) Normalized editing activity of (D). (F) The secondary structures of RNA variants were tested in (D), with the editable A highlighted in red. (G-H) Representative gels showing RNA editing activities of ADAR1 against GLI RNA variants (V13-V58) (G) and (V59-V63) (H). (I) Normalized editing activity of (H). (J) The secondary structures of RNA variants were tested in (H), with the editable A highlighted in red. (E) and (I). Values represent the mean of three independent experiments, with error bars indicating standard deviation. 120 nM ADAR1 and 25 nM each RNA were used.

**Figure S2.**
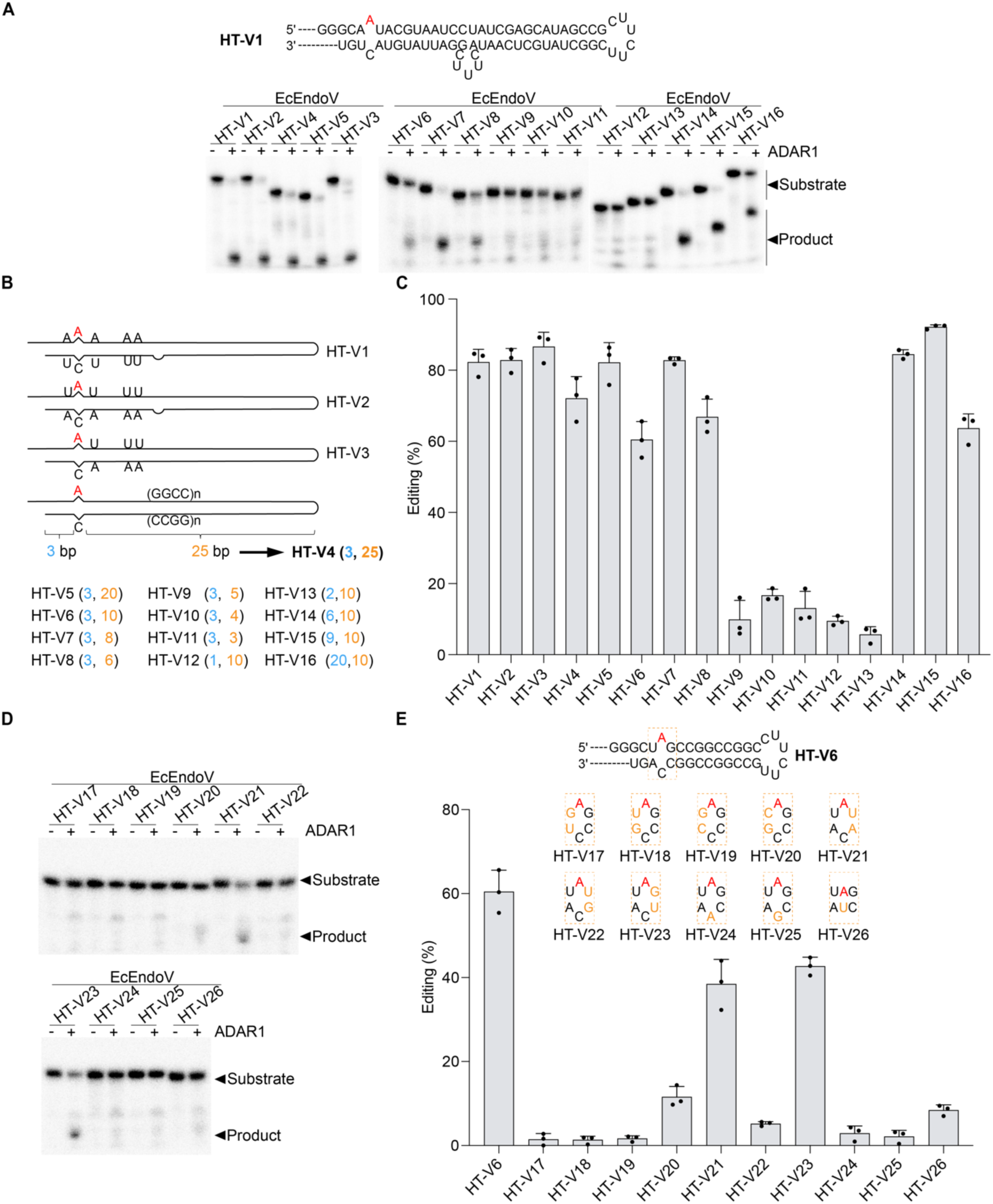
Biochemical characterization of ADAR1 editing on HT RNA variants, related to Figure 1. (A-C) Representative gels showing RNA editing activities of ADAR1 against HT RNA variants (V1-V16) (A). The secondary structures of HT-V1 to V4 are shown in (A) and (B), with the editable A highlighted in red. The lengths of the 5ʹ or 3ʹ arms of the RNA backbones are indicated, with mutated RNA variants of different lengths denoted in parentheses in (B). (C) Normalized editing activity of (A). (D) Representative gels showing RNA editing activities of ADAR1 against HT RNA variants (V17-V26). (E) Normalized editing activity of (D). HT-V6 RNA backbone is shown as a secondary structure. The editable A sites are highlighted in red, while mutated bases are enclosed in dashed boxes and labeled in orange in (E). Values represent the mean of three independent experiments, with error bars indicating standard deviation. 120 nM ADAR1 and 25 nM each RNA were used.

**Figure S3.**
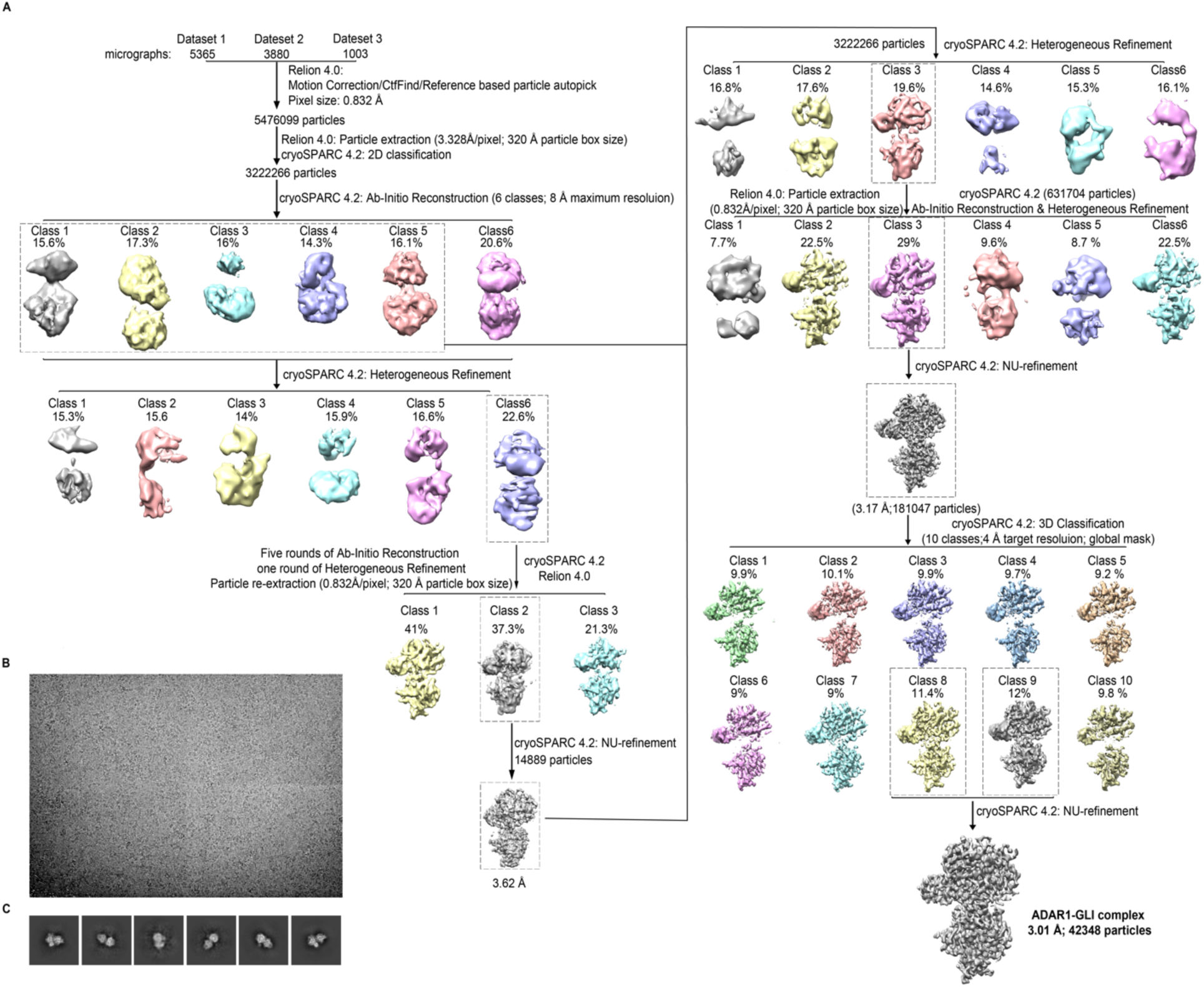
Cryo-EM reconstruction of ADAR1-GLI complex, related to Figure 2. (A) Overview of image processing and refinement strategy. (B) Representative cryo-EM micrograph. (C) Representative 2D class averages.

**Figure S4.**
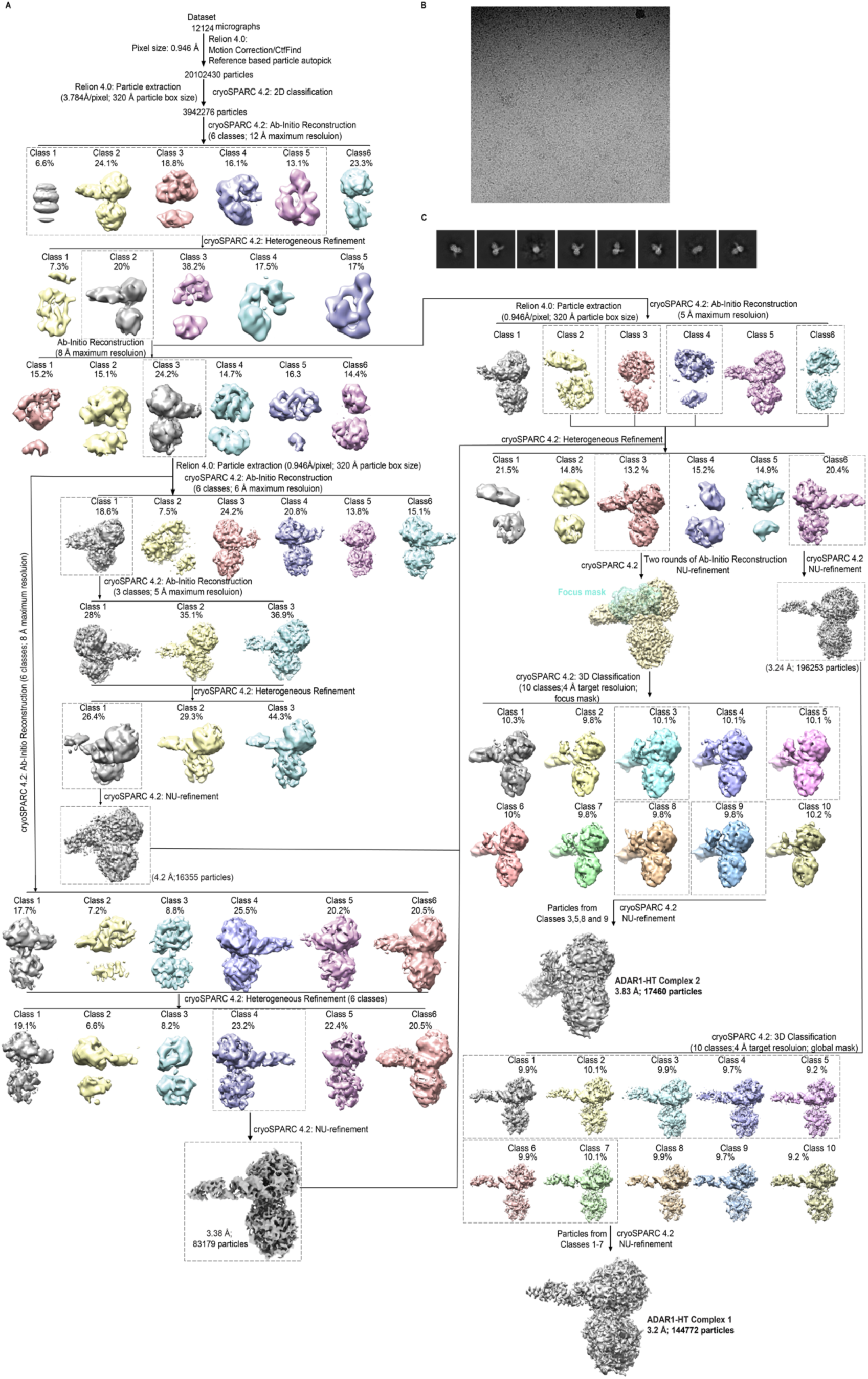
Cryo-EM reconstruction of ADAR1-HT complex and ADAR1-HT complex 2, related to Figure 2. (A) Overview of image processing and refinement strategy. (B) Representative cryo-EM micrograph. (C) Representative 2D class averages.

**Figure S5.**
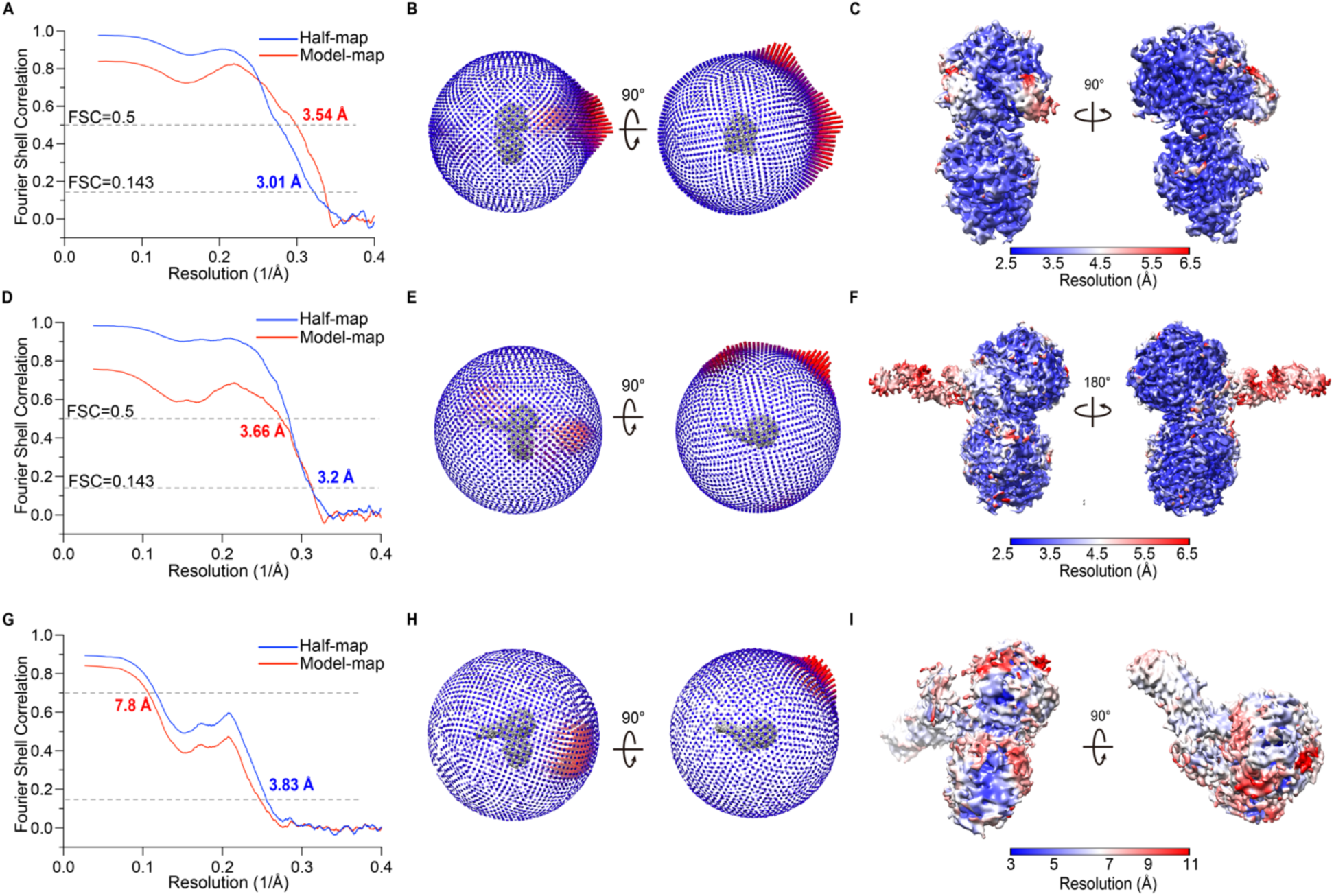
Cryo-EM analysis of ADAR1-RNA complexes, related to Figure 2. (A), (D), and (G) Gold-standard Fourier shell correlation (FSC) curves of ADAR1-GLI complex (A), ADAR1-HT complex (D), and ADAR1-HT complex 2 (G) between the two half maps, with indicated resolution at FSC = 0.143 shown in blue. FSC curves between the refined model and the cryo-EM map, with indicated resolution at FSC = 0.5, are shown in red, respectively. (B), (E), and (H) The angular distribution of particles used in the final 3D reconstruction of ADAR1-GLI complex (B), ADAR1-HT complex (E), and ADAR1-HT complex 2 (H). (C), (F), and (I) Local resolution estimation of the cryo-EM density map of ADAR1-GLI complex (C), ADAR1-HT complex (F), and ADAR1-HT complex 2 (I).

**Figure S6.**
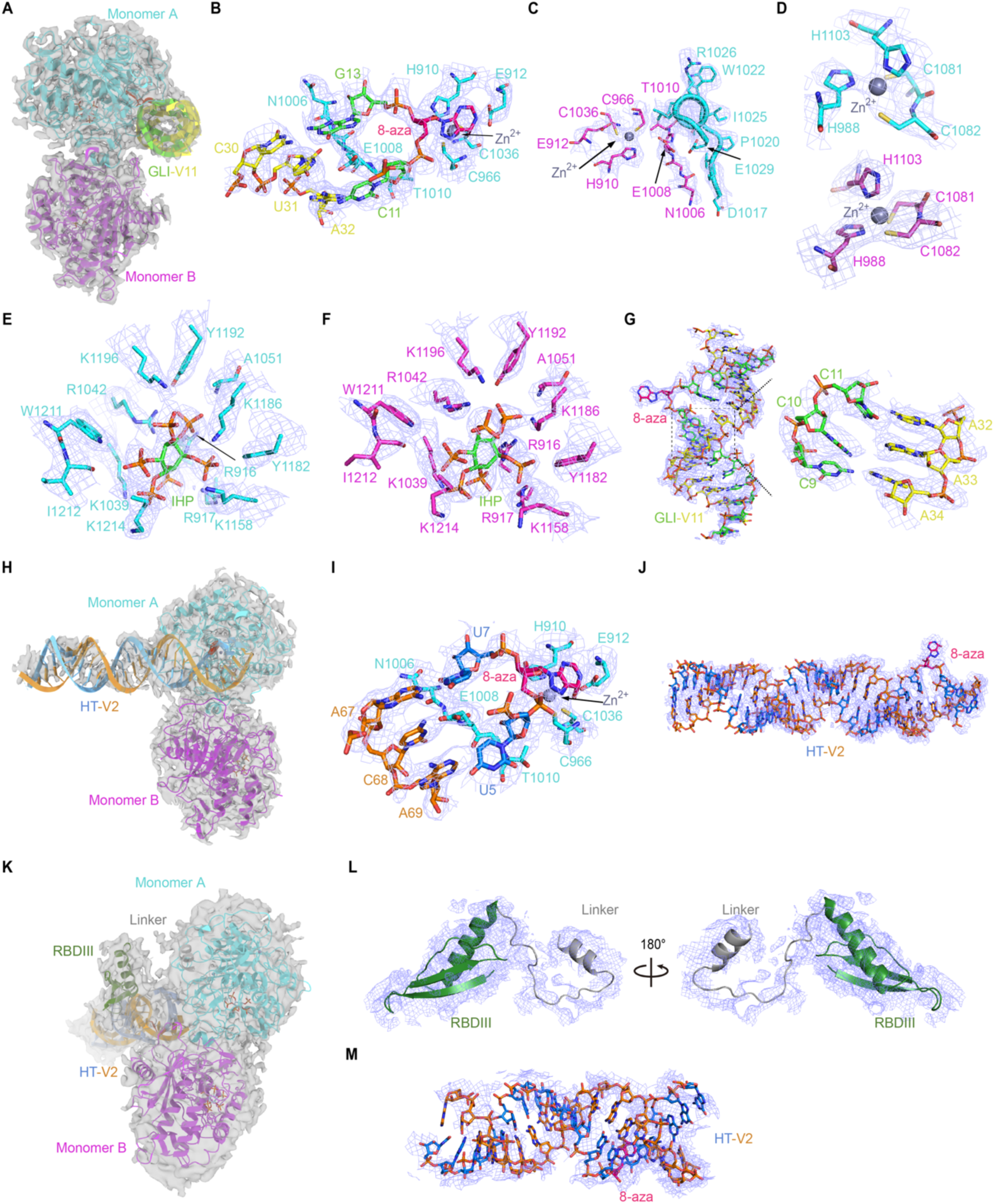
Structure models of ADAR-RNA complexes, related to Figures 2 and 3. (A) Cryo-EM density maps of the ADAR1-GLI complex. (B) The local density map of the catalytic sites in the ADAR1-GLI complex. (C) The local density map of the dimerization interface of ADAR1-GLI complex. (D) The local density map of the two Zn^2+^ binding motifs in the ADAR1-GLI complex. (E-F) The local density map of IHP and its contacted residues in the monomer A (E) and monomer B (F) in the ADAR1-GLI complex. (G) The cryo-EM density map of GLI-V11 RNA, with a zoom-in view of the 3 bp CCC:AAA bubble is shown in the right panel. (H) Cryo-EM density maps of the ADAR1-HT complex. (I) The local density map of the catalytic sites in the ADAR1-HT complex. (J) Cryo-EM density map of HT-V2 RNA in ADAR1-HT complex. (K) Cryo-EM density maps of the ADAR1-HT complex 2. (I) The local density map of the RBD3 and linker. (M) Cryo-EM density map of HT-V2 RNA in ADAR1-HT complex 2.

**Figure S7.**
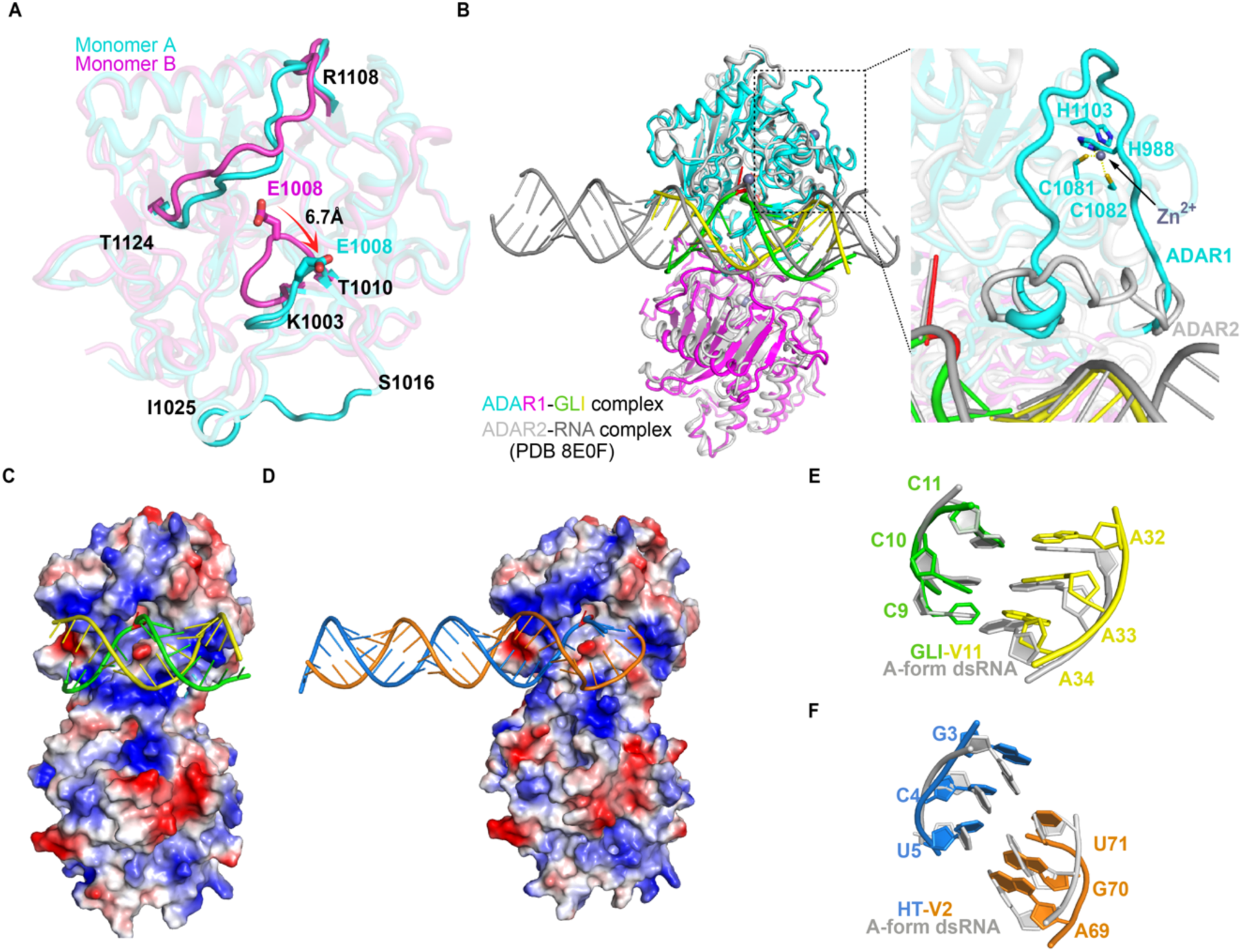
Structural comparison of ADAR1-RNA and ADAR2-RNA complexes, related to Figures 2 and 3. (A) Structural comparison of the monomer A and B from ADAR1-GLI complex. The shift of E1008 residue is indicated with a red arrow. (B) Structural comparison of the ADAR1-GLI and ADAR2-RNA complex (PDB 8E0F), the monomer A from two structures aligned. Only the deaminase domain and RNA in the ADAR2-RNA complex are shown for clarity and colored in grey. The zoom-in view of the flexible loop in ADAR1 and ADAR2 is shown on the right panel. (C-D) The electrostatic potential surface of ADAR1-GLI (C) and ADAR1-HT complex (D). (E-F) Structural comparison of the 3-bp "CCC:AAA" bubble on the 5’ arm of the GLI RNA (E) and the short 3-bp “GCU:UGA” 5’ duplex of the HT RNA with regular A-form dsRNA (F).

**Figure S8.**
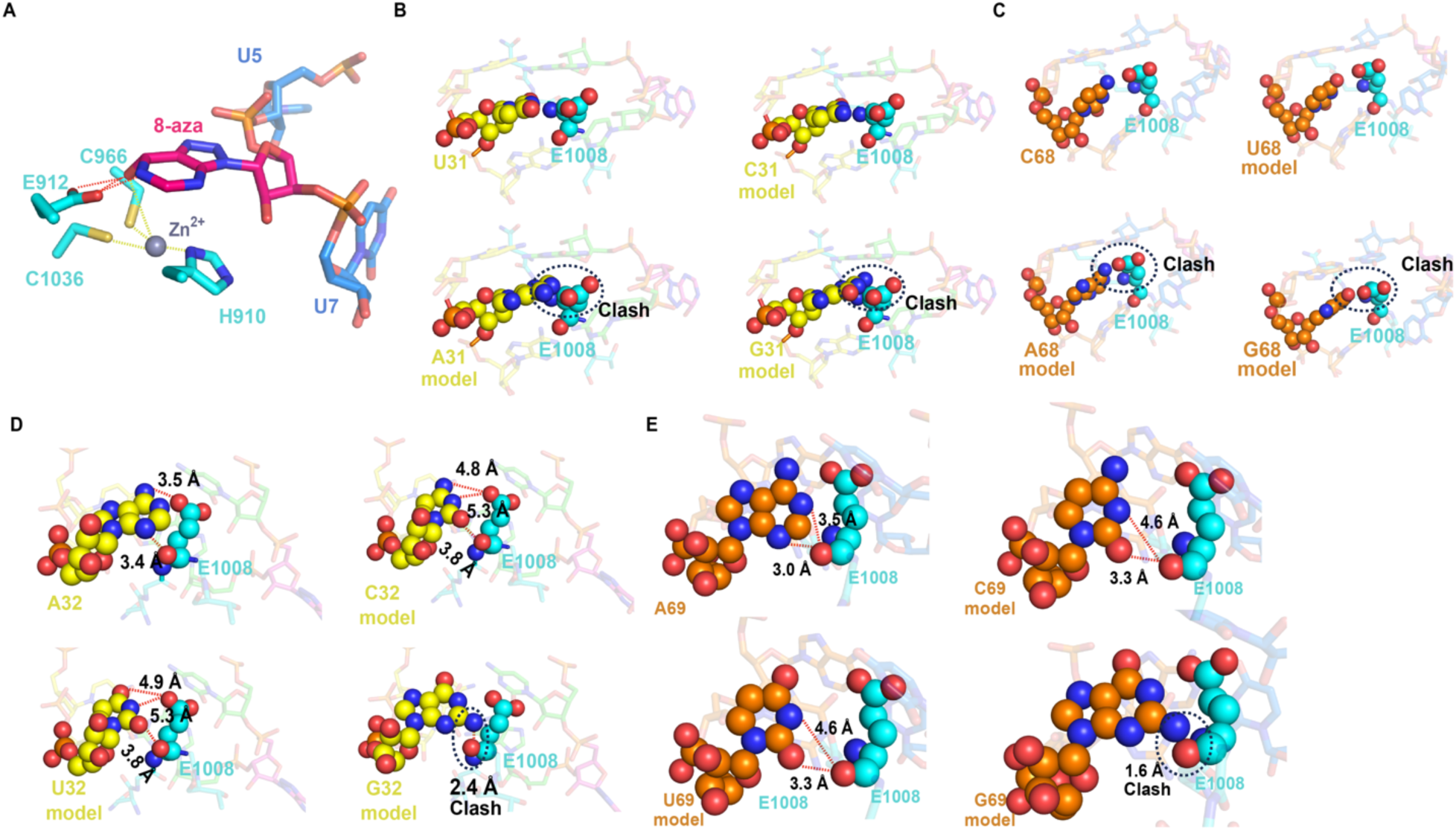
Recognition of the flipped 8-aza and structural modeling of nucleotides near the editing site, related to Figure 3. (A) Recognition of the flipped 8-aza in the active site, the hydrogen bond interactions are shown as red dashed lines. (B-C) Structural modeling of replacing the orphan base in ADAR1 GLI (B) or HT (C) complex. (D-E) Structural modeling of replacing 5ʹ-adjacent “A” on the non-editing strand in ADAR1 GLI (D) or HT (E) complex. Clashes were defined as the hydrogen bond interaction shorter than 2.5 Å in (B-E).

**Figure S9.**
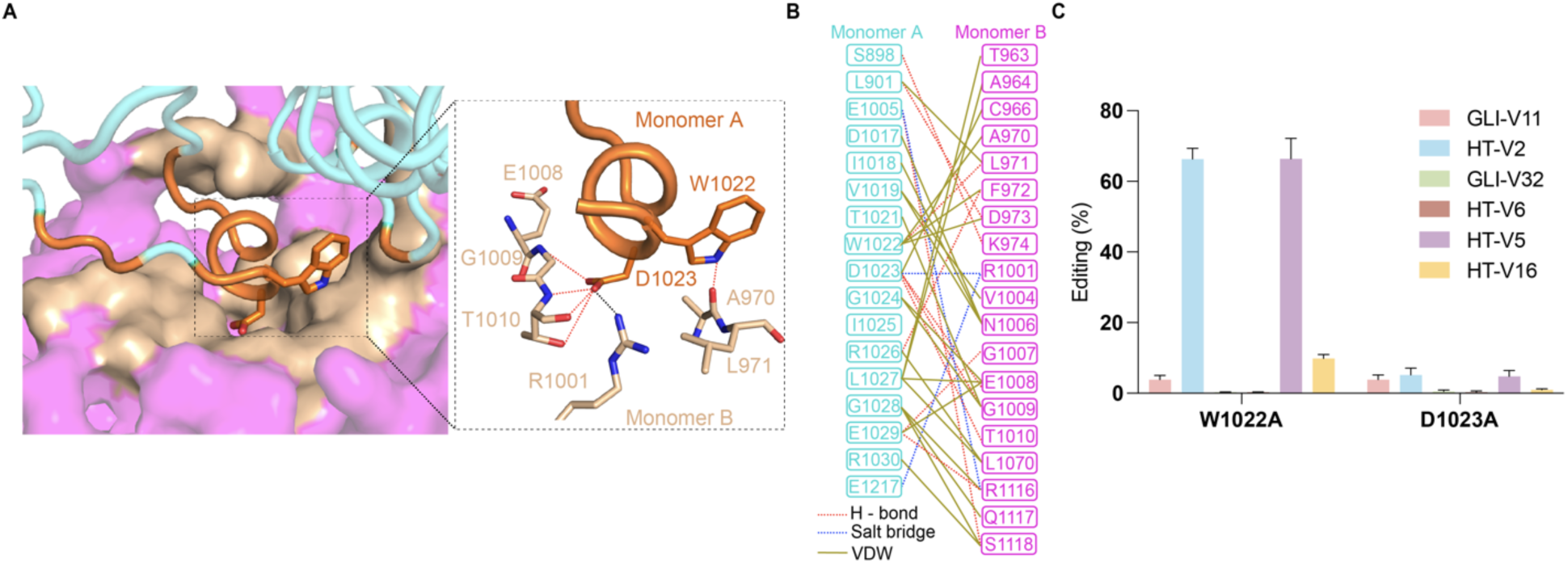
The dimerization interface of ADAR1, related to Figure 3. (A) The dimerization interface between monomer A and monomer B in the ADAR1-GLI complex, the monomer A and monomer B are color-coded as in Figure 2C, while the interfaces on monomer A and B are colored in orange and wheat. The zoom-in view of the interactions between the two key residues (W1022 and D1023) from monomer A and their interacting residues in monomer B are shown on the right panel. (B) The molecular contacts between monomer A and monomer B. VDW: van der Waals interactions. (C) Editing activity of W1022A and D1023A against different RNA substrates. Values represent the mean of three independent experiments, with error bars representing the standard deviation.

**Figure S10.**
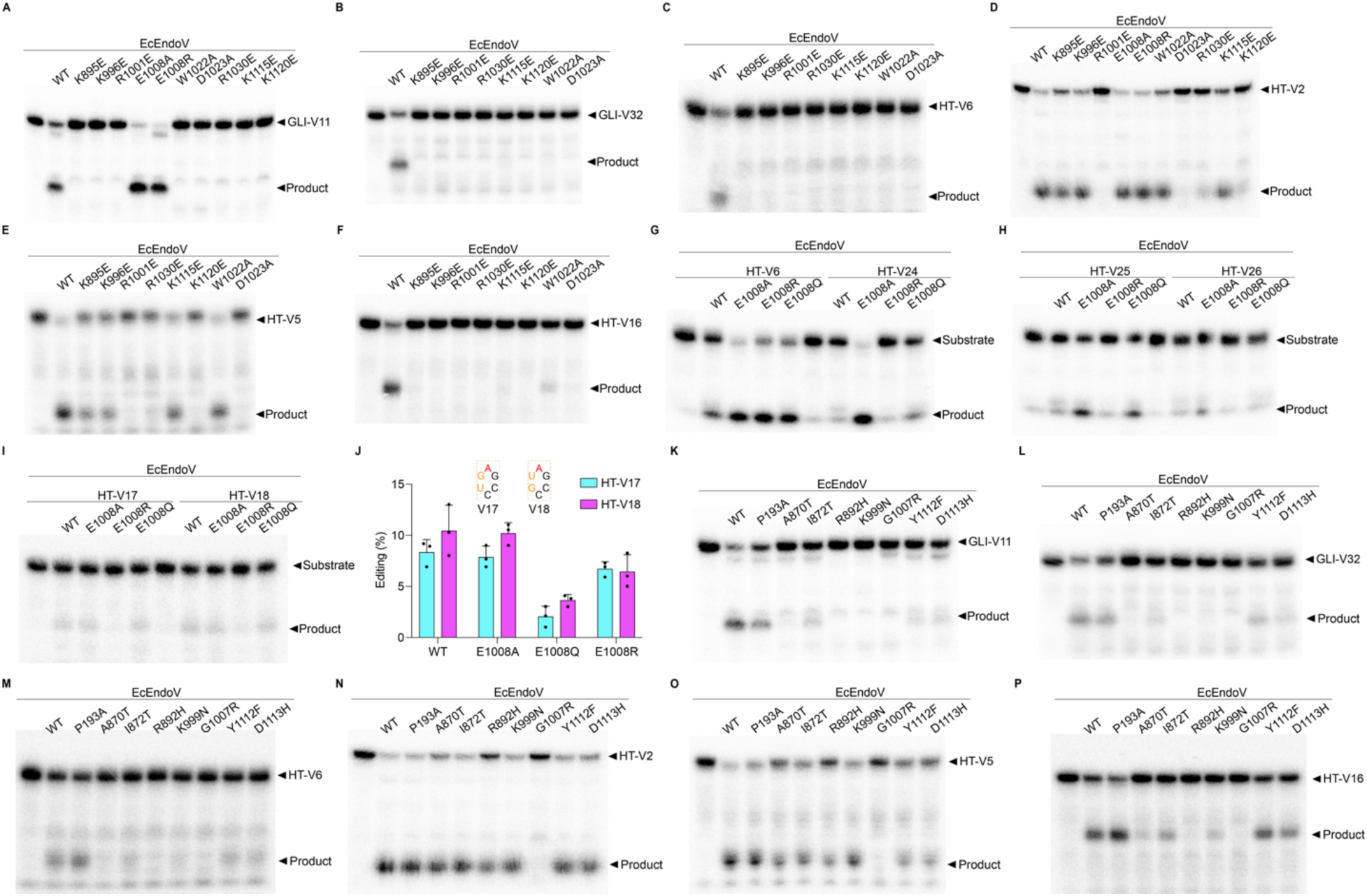
Mutagenesis studies of ADAR1, related to Figures 3 and 4. (A-F) Representative gels of the RNA editing activities of ADAR1 RNA binding or dimerization-related mutants against GLI-V11 (A), GLI-V32 (B), HT-V6 (C), HT-V2 (D), HT-V5 (E), and HT-V16 (F) RNAs. (G-I) Representative gels of the RNA editing activities of ADAR1 E1008 mutants against HT-RNA variants. (J) Normalized editing activity as shown in (I). (K-P) Representative gels of the RNA editing activities of AGS-associated mutants against GLI-V11 (K), GLI-V32 (I), HT-V6 (M), HT-V2 (N), HT-V5 (O), and HT-V16 (P) RNAs. 100 nM ADAR1 or mutants and 25 nM each RNA were used. The values represent the mean of three independent experiments, with the error bars representing the standard deviation.

**Figure S11.**
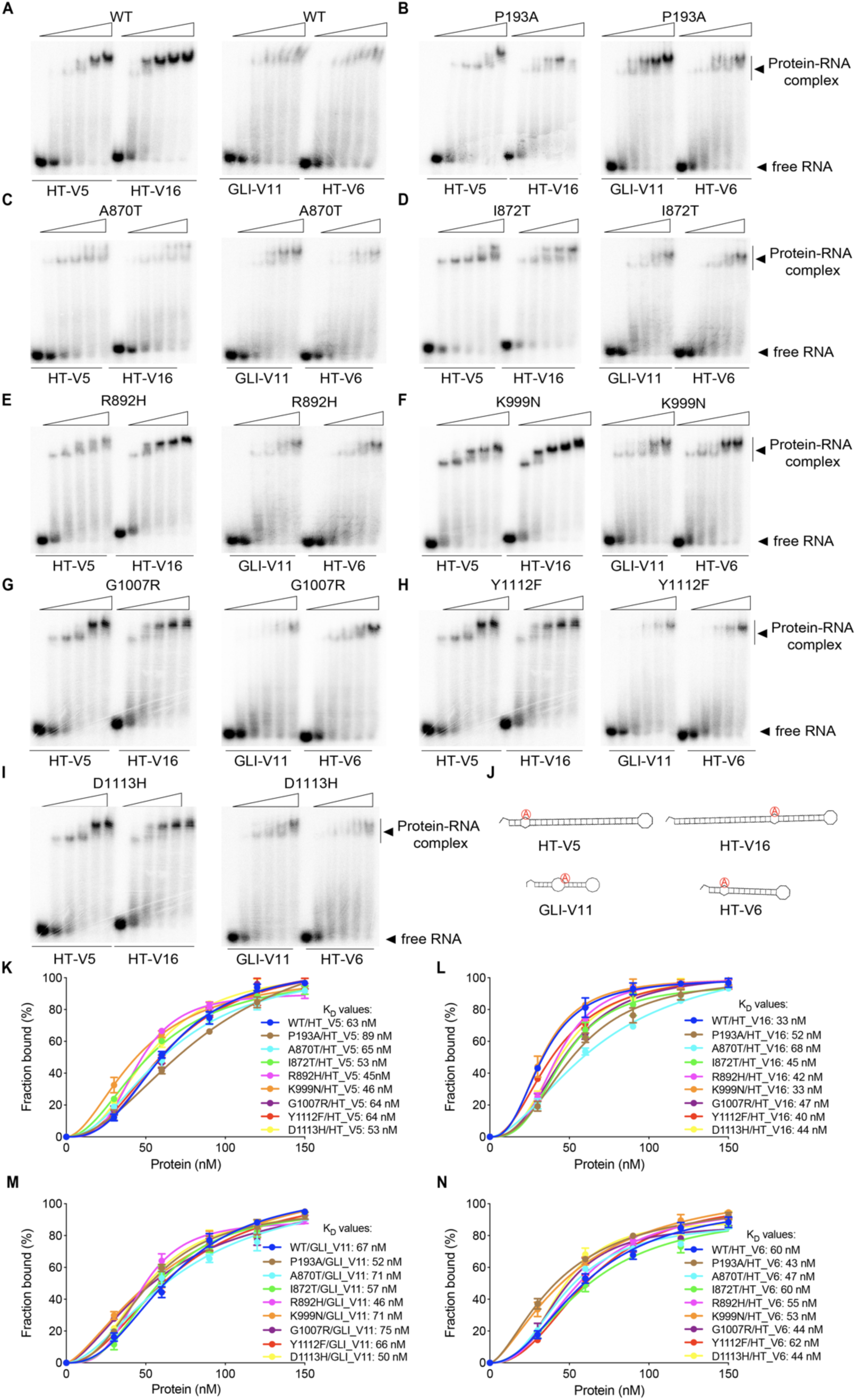
RNA binding affinity characterization of ADAR1 AGS-associated mutations, related to Figure 4. (A-I) Representative EMSA gel images of WT and ADAR1-associated mutants complex with GLI-V11, HT-V5, HT-V6, or HT-V16. (J) The secondary structures of GLI-V11, HT-V5, HT-V6, and HT-V16. (K-N) Calculation of binding affinity between WT or ADAR1-associated mutant with HT-V5 (K), HT-V16 (L), GLI-V11 (M), and HT-V6 (N) based on data from (A-I). The concentration of protein is serially diluted to 150, 120, 90, 60, and 30 nM. The concentration of each RNA variant is 5 nM. Bound and unbound fractions were quantified by densitometry. At least two times each experiment was repeated independently with similar results (A-I).

**Figure S12.**
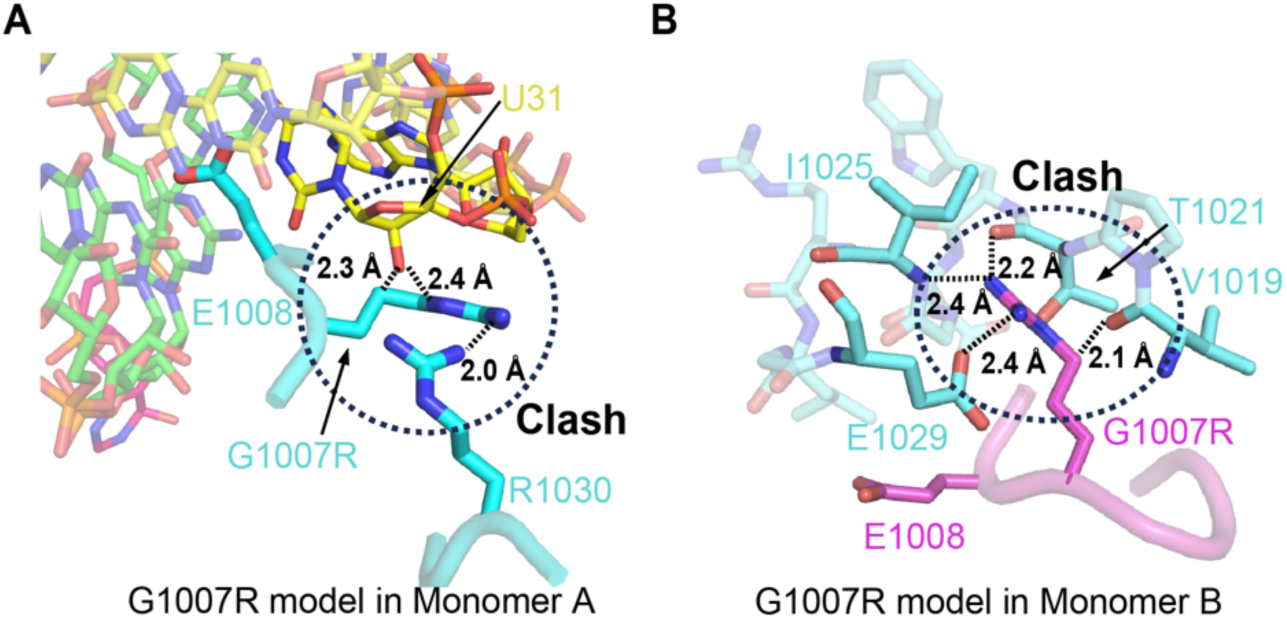
Structural modeling of G1007R mutation in ADAR1-GLI complex, related to Figure 4. (A) In monomer A, the G1007R mutation introduces steric clashes with both the orphan base U31 and residue R1030. (B) In monomer B, the G1007R mutation disrupts the dimerization interface through steric clashes with residues V1019, T1021, I1025, and E1029 from monomer A. Clashes were defined as the hydrogen bond interaction shorter than 2.5 Å.

**Figure S13.**
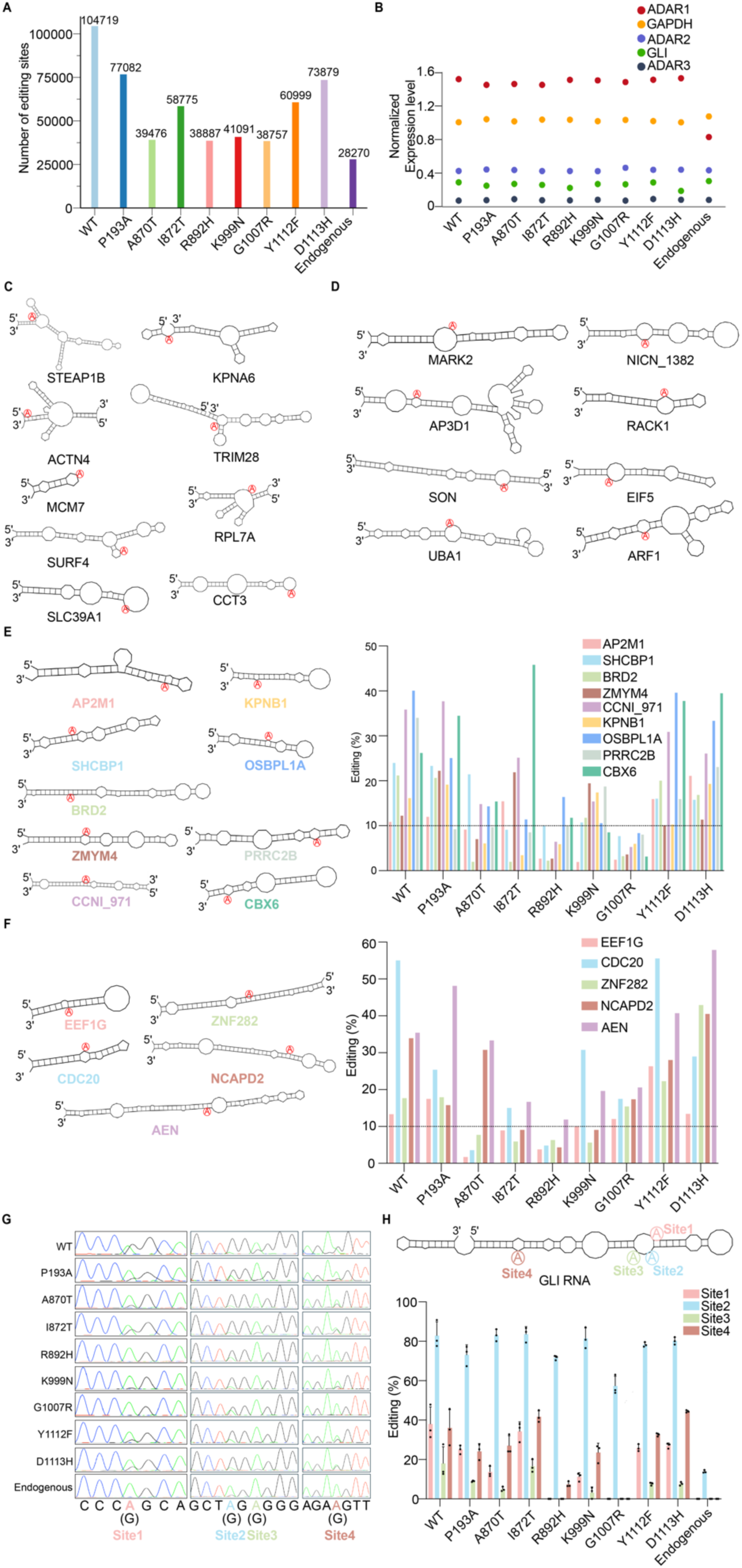
Impaired RNA editing activity of ADAR1 AGS mutants. Related to Figure 4. (A) Overall RNA editing events observed in WT ADAR1, AGS-associated mutants, and the empty vector control groups from RNA-Seq analysis. Editing sites were selected based on a minimum of 5 reads and an editing rate exceeding 1%. (B) RNA expression levels of ADAR1, ADAR2, ADAR3, and GLI in each transfection group. Expression levels were normalized to GAPDH RNA. (C–D) Secondary structures of RNAs which were excluded from activity comparisons as the editing site was located outside dsRNA regions (B) or positioned on large bulges or near branching structures (D). The editing site A is highlighted in red. (E-F) The editing activity of WT ADAR1 and AGS-associated mutants was evaluated on the remaining dsRNAs (E) and with the G1007R mutant showing high background editing (>10%) for five RNA substrates (F) identified from RNA-Seq. Secondary structures of the RNAs are shown, with the editing site adenosine A highlighted in red. The black dotted line indicates 10% editing activity as a reference. (G) Validation of A-to-G RNA editing in GLI RNA by Sanger sequencing. The editing site is labeled, and the difference in peak change from adenosine (A) to guanosine (G, editing product) is related to the editing activity of each transfection group. (H) Normalized editing activity, as shown in (G). Values represent the mean of three independent experiments, with error bars indicating the standard deviation.

**Figure S14.**
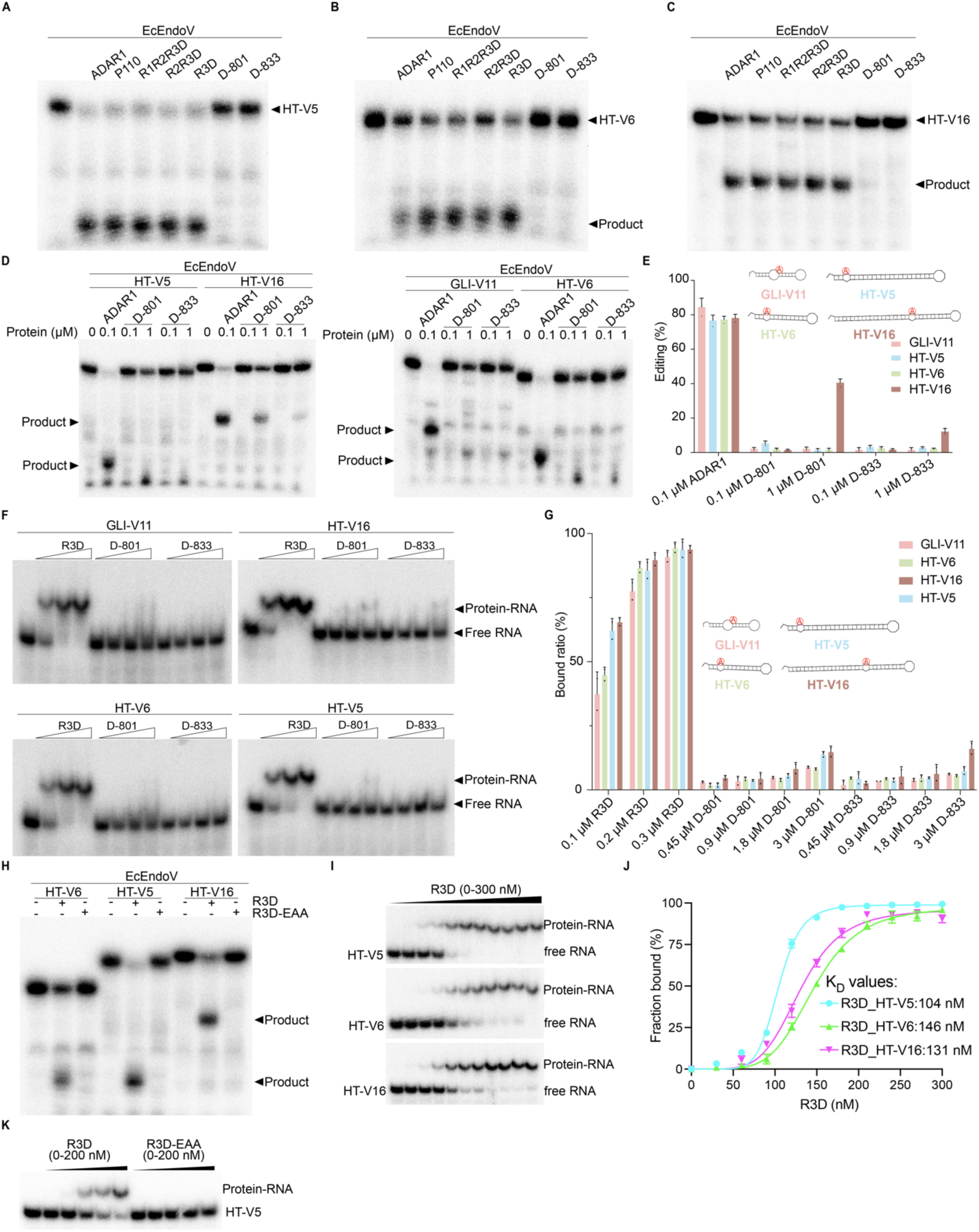
Roles of RBDs in ADAR1 editing, related to Figure 5. (A-C) Representative gels of the RNA editing activities of ADAR1 domain truncation mutants against HT-V5 (A), HT-V6 (B), and HT-V16 (C). 100 nM ADAR1 or mutants and 25 nM each RNA were used. (D) Representative gels of the RNA editing activities of ADAR1 or deaminase domain alone (D-801 and D-833) against GLI-V11, HT-V5, HT-V6, and HT-V16. The concentration of protein was indicated, the concentration of each RNA is 25 nM. (E) Normalized editing activity as shown in (D). (F) Representative EMSA gel images of R3D and deaminase domain alone (D-801 and D-833) complex with GLI-V11, HT-V5, HT-V6, or HT-V16. The concentration of R3D is serially diluted to 300, 200 and 100 nM, while the concentration of D-801 or D-833 is serially diluted to 3000, 1800, 900, and 450 nM. The concentration of each RNA variant is 25 nM. (G) Normalized RNA binding affinity as shown in (F). (H) Representative gels of the RNA editing activities of R3D and R3D-EAA mutants against the indicated RNAs. (I) Representative EMSA gel images of R3D complex with HT-V5, HT-V6, or HT-V16. The concentration of R3D is serially diluted to 300, 270, 240, 210, 180, 150, 120, 90, 60, and 30 nM. The concentration of each RNA variant is 10 nM. (J) Calculation of binding affinity between R3D and HT-V5, HT-V6, or HT-V16. Bound and unbound fractions were quantified by densitometry. (K) Representative EMSA gel images of R3D or R3D-EAA mutant complex with HT-V5. The protein concentration is serially diluted to 200, 160, 120, 80, and 40 nM. The concentration of HT-V5 is 10 nM. In (A-C), the deaminase domain alone (D-801 and D-833) with the MBP tag was used, while in (D) and (F), the MBP tag was removed. At least three times each experiment was repeated independently with similar results (A-D, and H). At least two times each experiment was repeated independently with similar results (F, I, and K).

**Figure S15.**
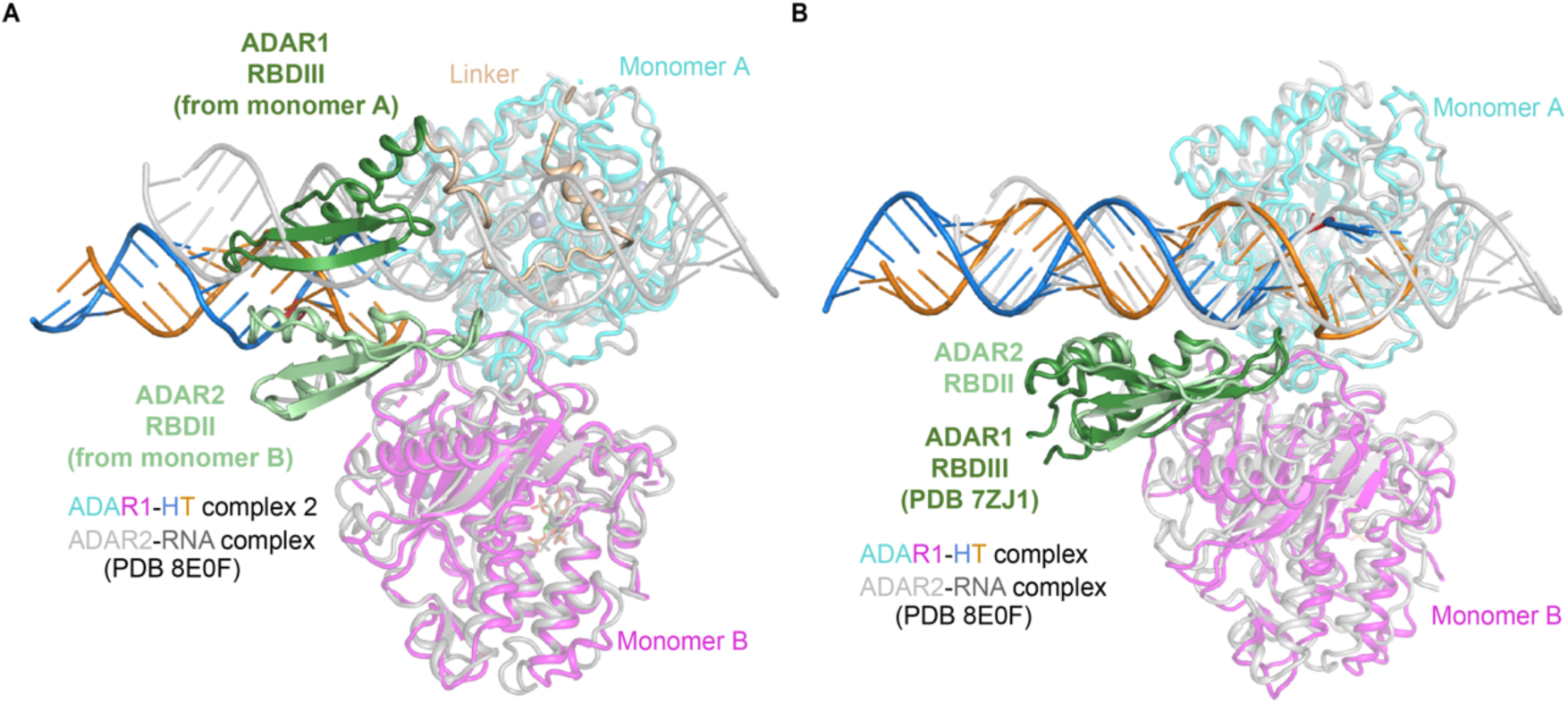
Roles of RBD3 in ADAR1 editing, related to Figure 5. (A) Structural comparison of ADAR1-RNA complex 2 and ADAR2-RNA complex (PDB 8E0F), with the monomer A from two structures aligned. The RBD3 from monomer A of ADAR1-RNA complex 2 was colored in dark green, while the RBD2 from monomer B of ADAR2-RNA complex was colored in light green. (B) Structural comparison of ADAR1-RNA complex and ADAR2-RNA complex (PDB 8E0F), with the monomer A from two structures aligned. The structure of RBD3 of ADAR1 (PDB 7ZJ1) (colored in dark green) was aligned with the RBD2 (colored in light green) from monomer B of ADAR2-RNA complex.

**Figure S16.**
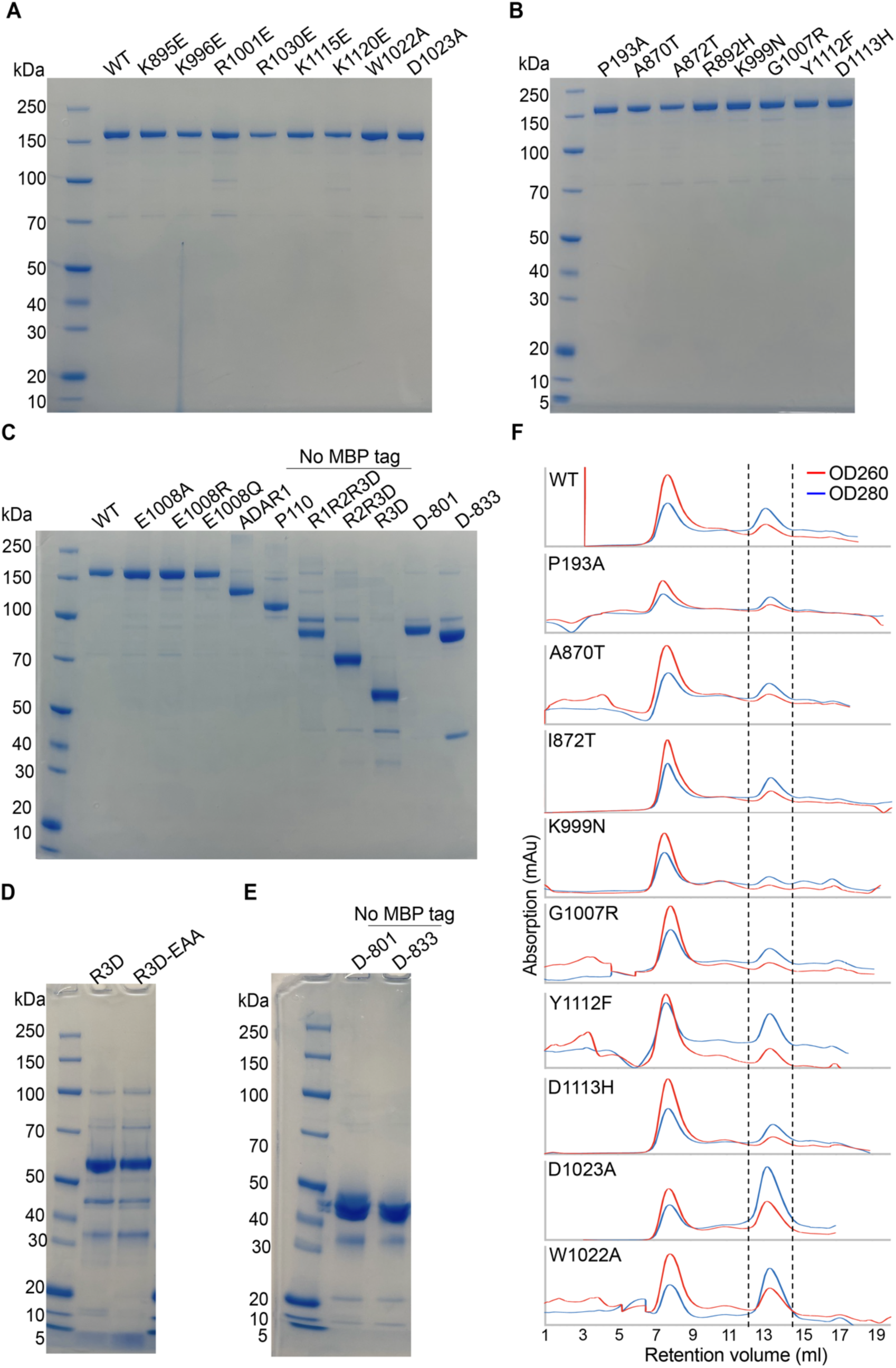
SDS-PAGE and size-exclusion chromatography profiles of purified ADAR1 proteins, related to Figures 4 and 5. (A-D) SDS-PAGE analysis of purified ADAR1 proteins after size-exclusion column purification, the truncation proteins without MBP tag were indicated. (F) Size-exclusion chromatography profiles of purified ADAR1 AGS-associated mutants as well as dimerization defect mutants with W1022A or D1023A mutation. Dashed lines indicate the elution peaks corresponding to the protein fractions.

**Table S1.**
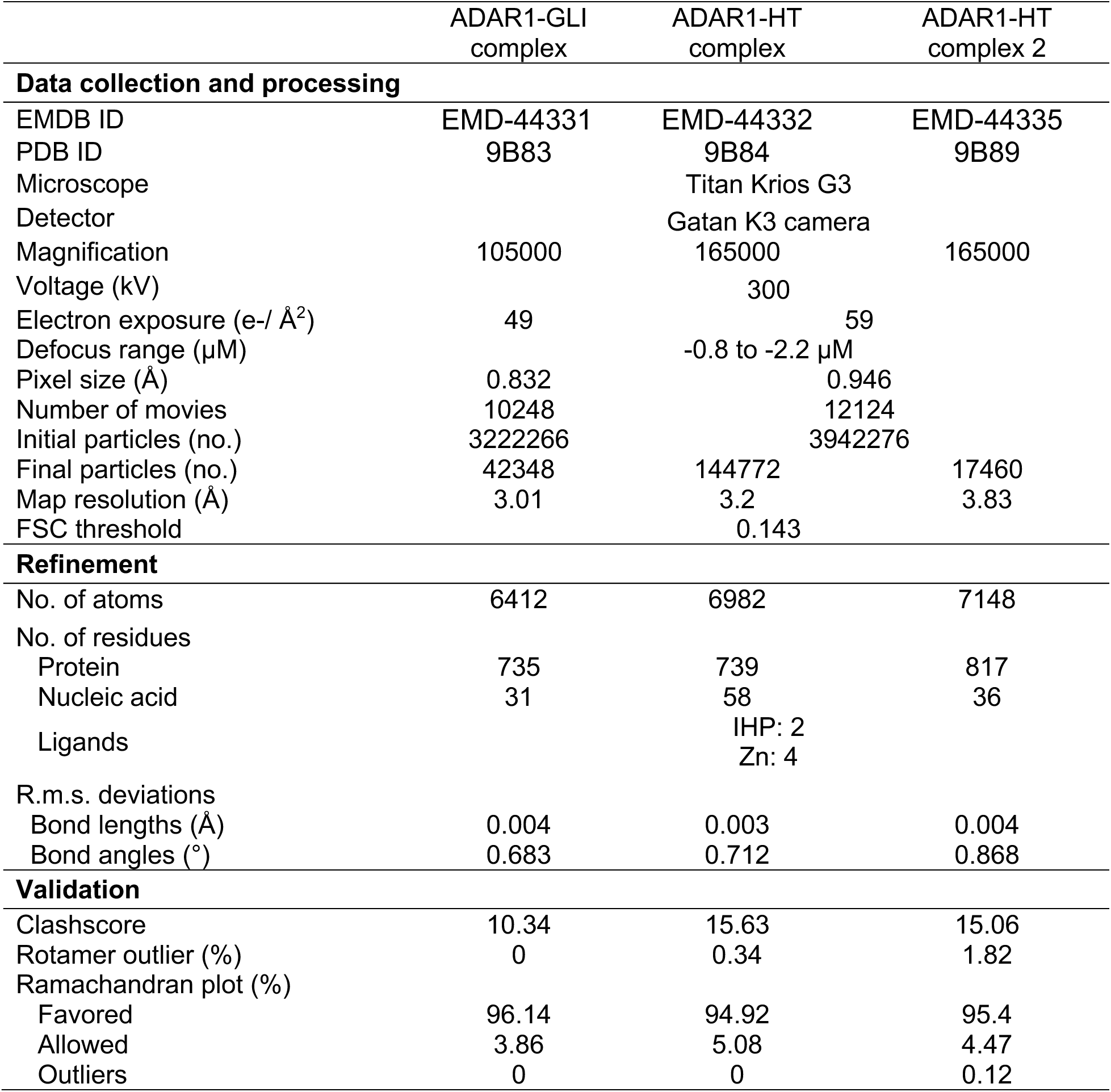
Cryo-EM data collection, model refinement, and validation.

Cryo-EM data collection, model refinement, and validation
|  | ADAR1-GLI<br>complex | ADAR1-HT<br>complex | ADAR1-HT<br>complex 2 |
| --- | --- | --- | --- |
| **Data collection and processing** |  |  |  |
| EMDB ID | EMD-44331 | EMD-44332 | EMD-44335 |
| PDB ID | 9B83 | 9B84 | 9B89 |
| Microscope |  | Titan Krios G3 |  |
| Detector |  | Gatan K3 camera |  |
| Magnification | 105000 | 165000 | 165000 |
| Voltage (kV) |  | 300 |  |
| Electron exposure (e-/Å<sup>2</sup>) | 49 | 59 |  |
| Defocus range (μM) |  | -0.8 to -2.2 μM |  |
| Pixel size (Å) | 0.832 | 0.946 |  |
| Number of movies | 10248 | 12124 |  |
| Initial particles (no.) | 3222266 | 3942276 |  |
| Final particles (no.) | 42348 | 144772 | 17460 |
| Map resolution (Å) | 3.01 | 3.2 | 3.83 |
| FSC threshold |  | 0.143 |  |
| **Refinement** |  |  |  |
| No. of atoms | 6412 | 6982 | 7148 |
| No. of residues |  |  |  |
| Protein | 735 | 739 | 817 |
| Nucleic acid | 31 | 58 | 36 |
| Ligands |  | IHP: 2<br>Zn: 4 |  |
| R.m.s. deviations |  |  |  |
| Bond lengths (Å) | 0.004 | 0.003 | 0.004 |
| Bond angles (°) | 0.683 | 0.712 | 0.868 |
| **Validation** |  |  |  |
| Clashscore | 10.34 | 15.63 | 15.06 |
| Rotamer outlier (%) | 0 | 0.34 | 1.82 |
| Ramachandran plot (%) |  |  |  |
| Favored | 96.14 | 94.92 | 95.4 |
| Allowed | 3.86 | 5.08 | 4.47 |
| Outliers | 0 | 0 | 0.12 |

**Table S2.**
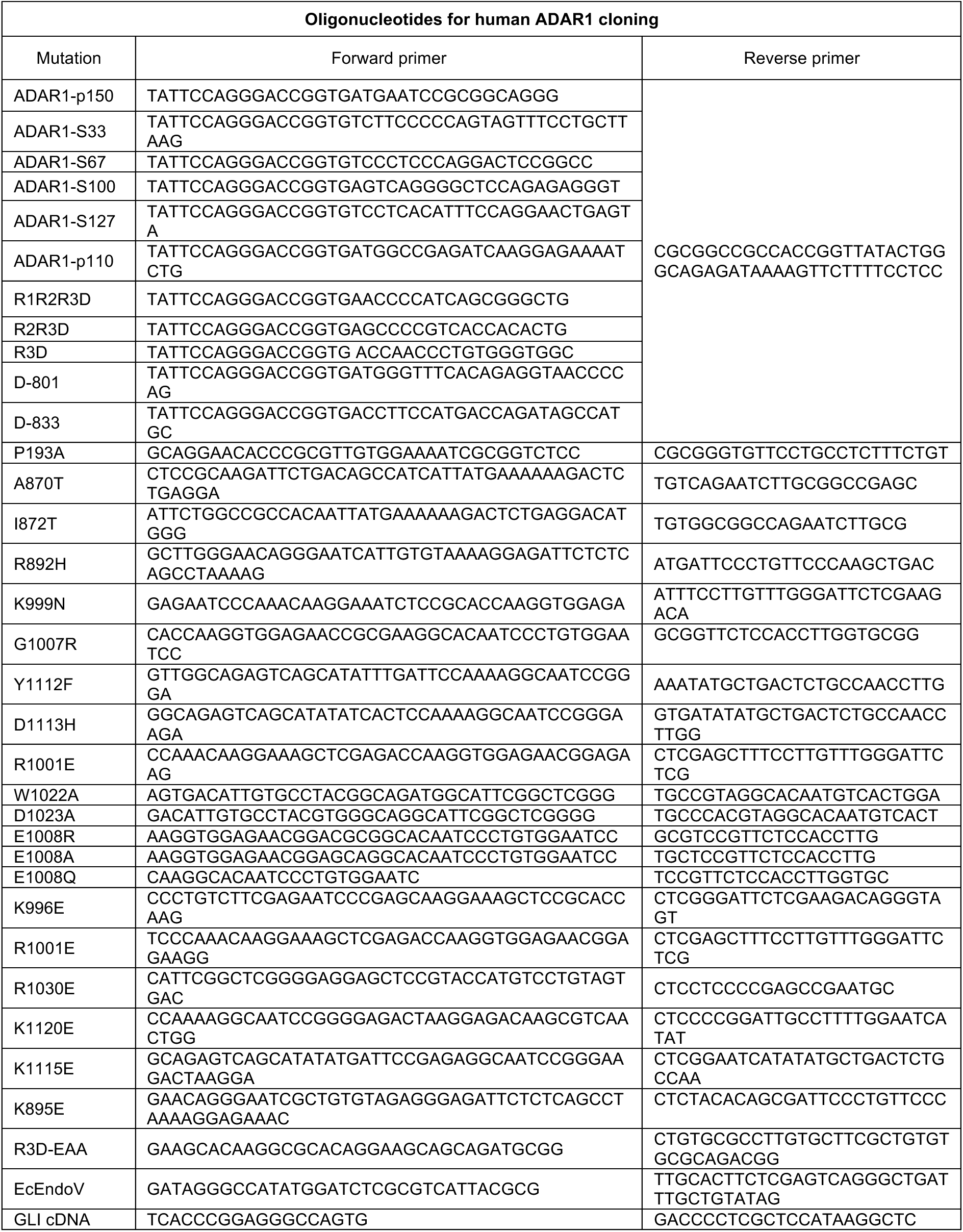

**Table S3.**
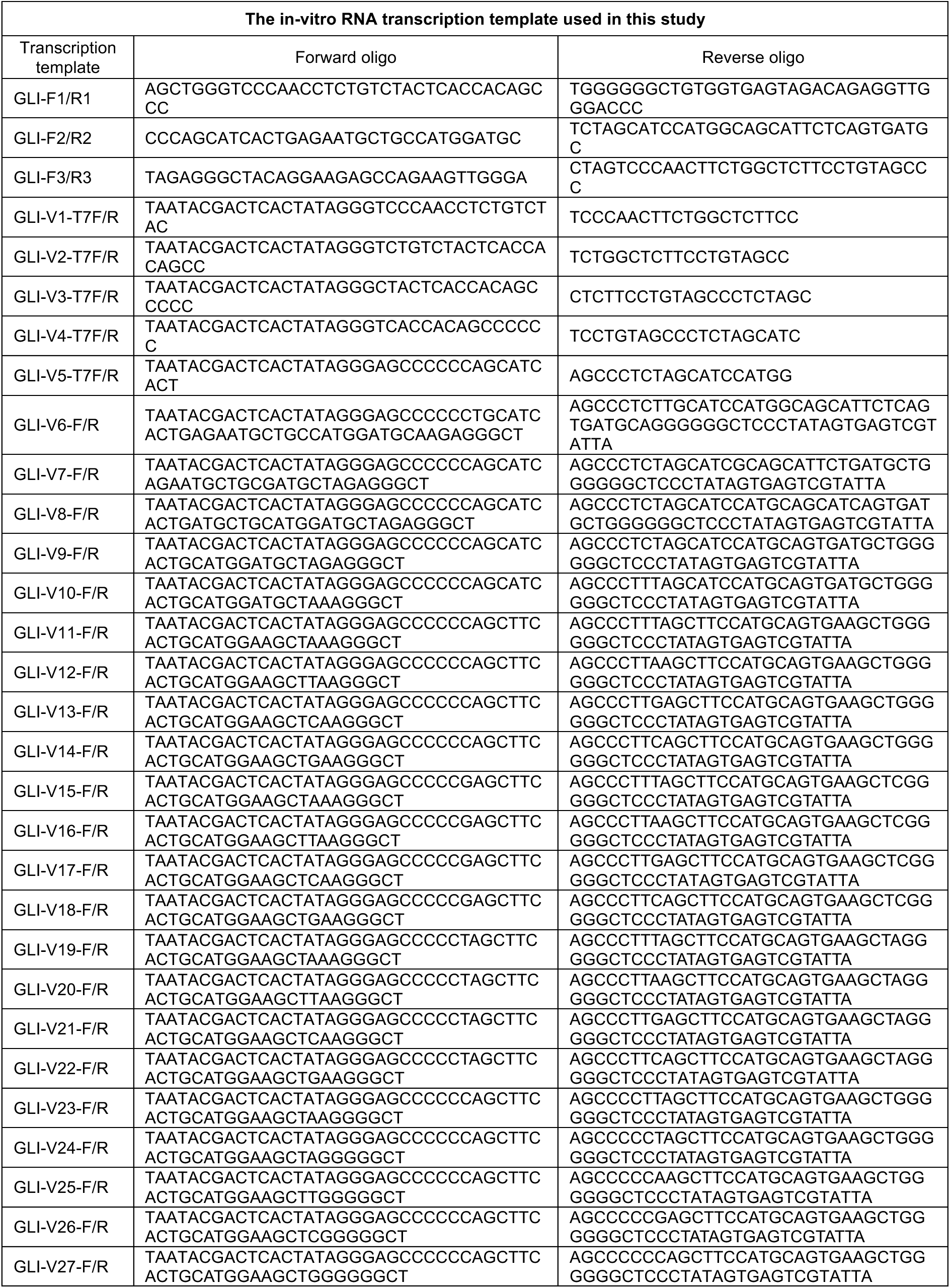
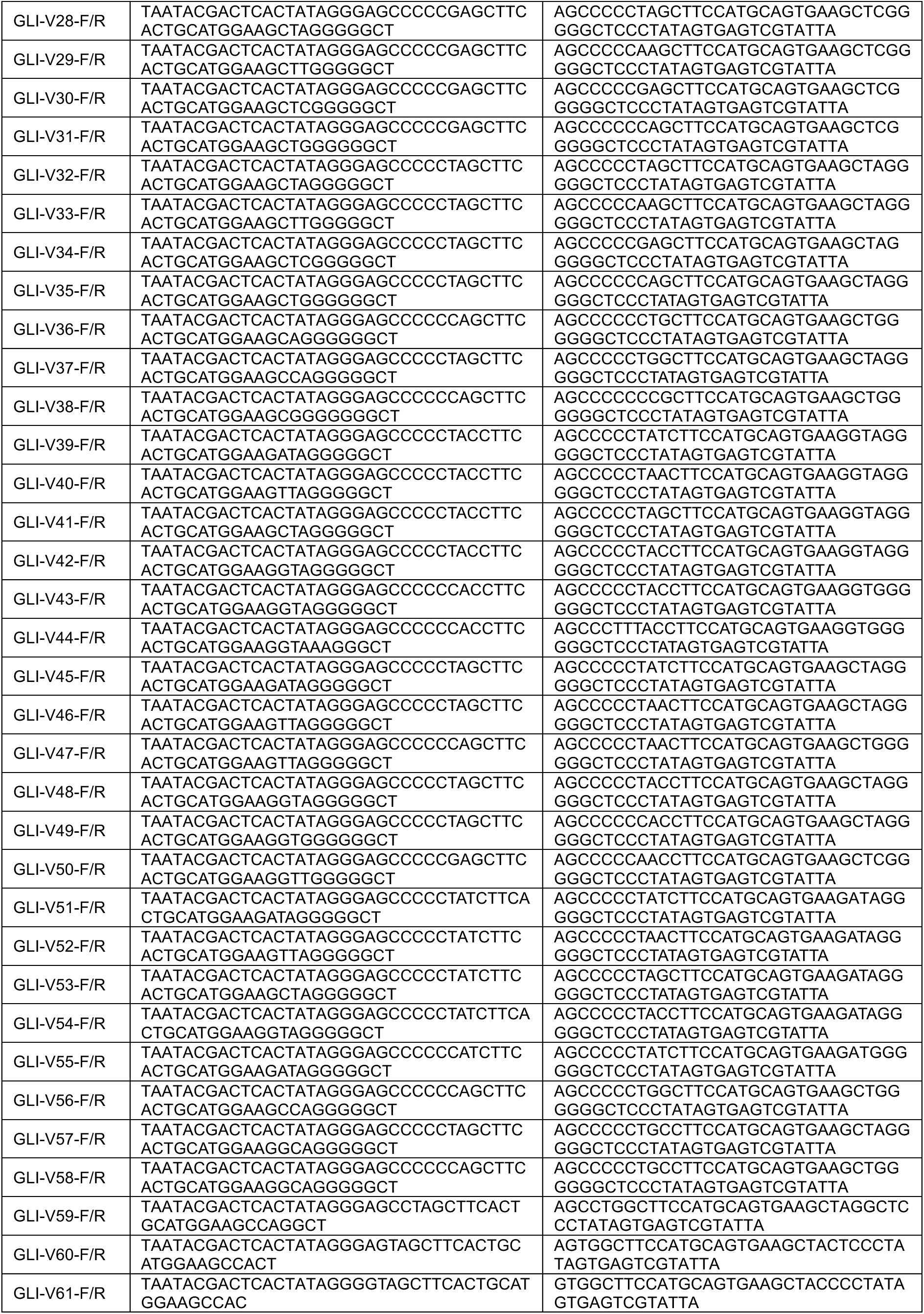
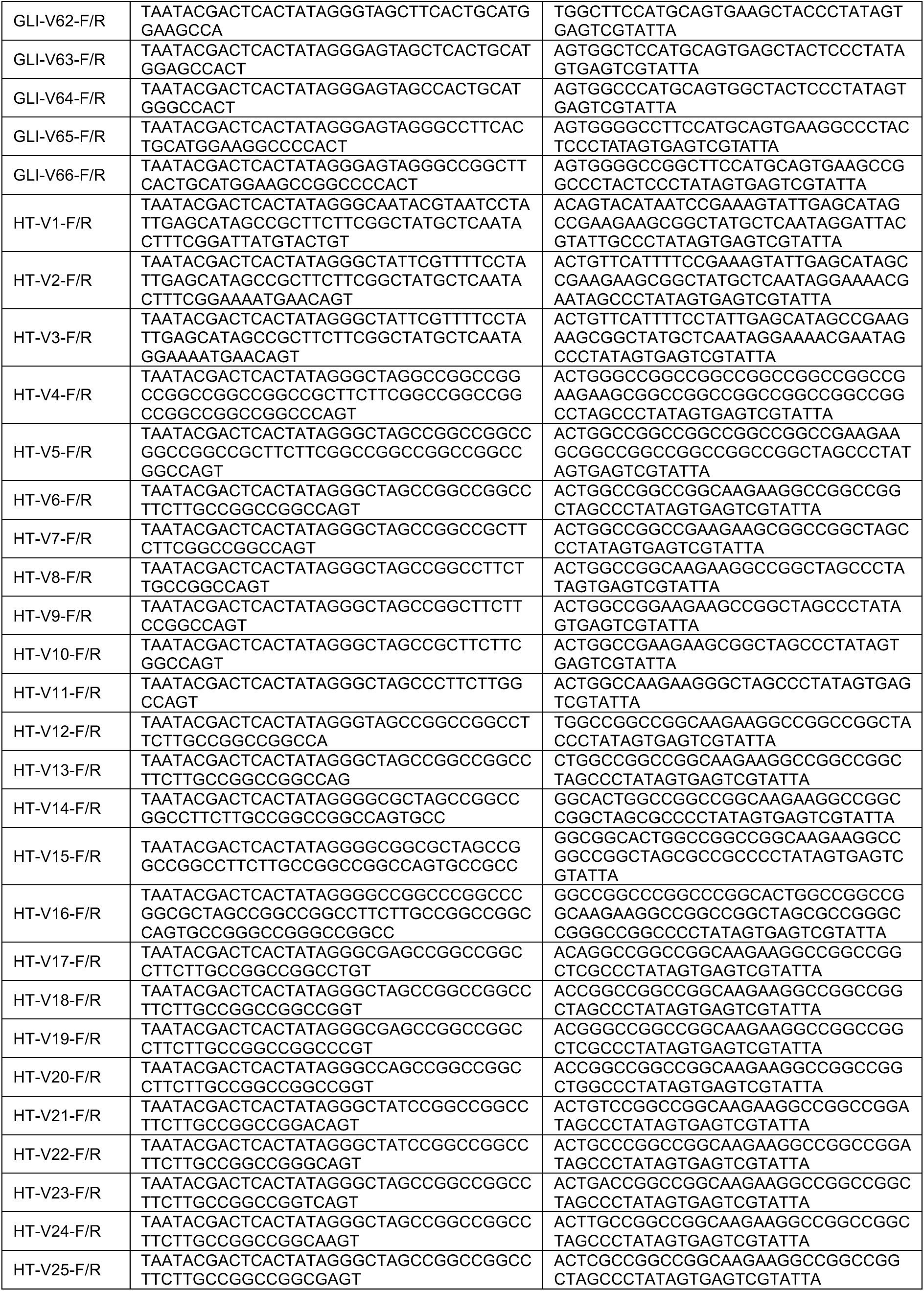
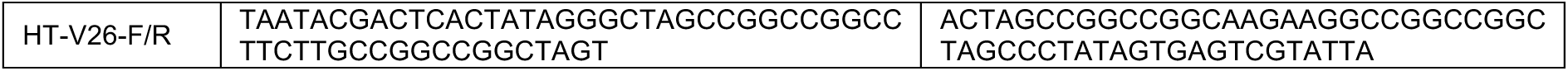

**Table S4.**
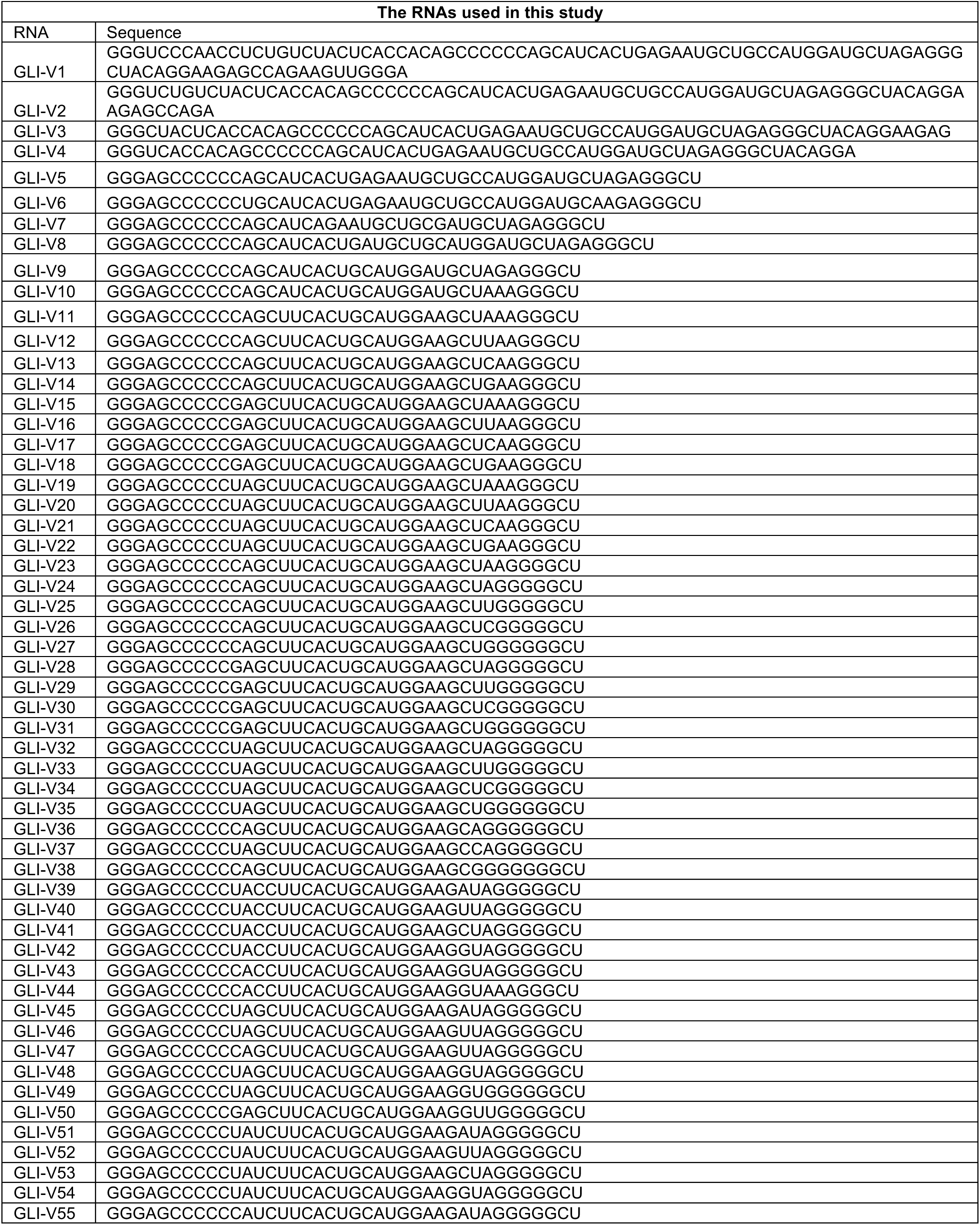
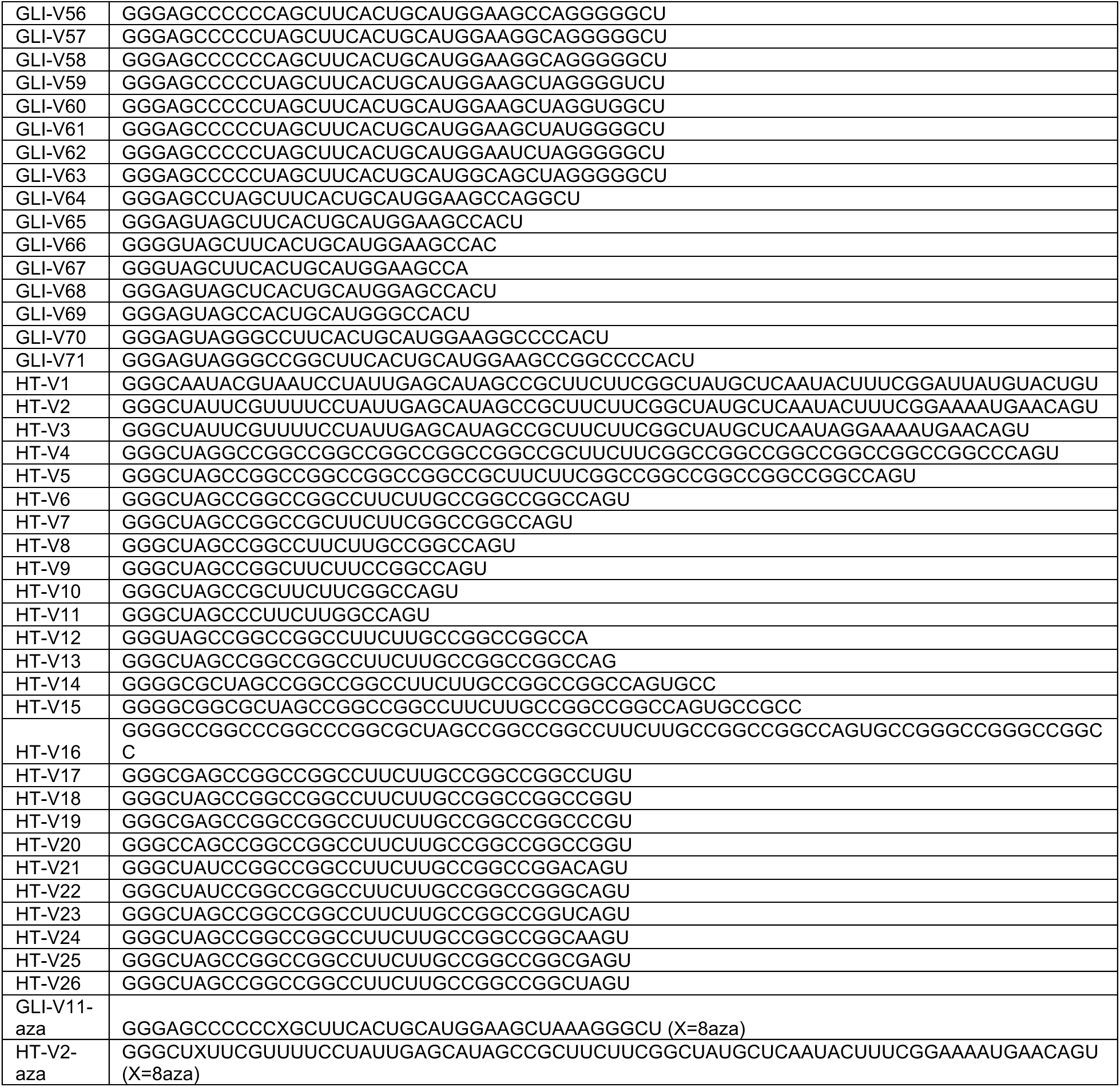

## STAR Methods

### Key resources table

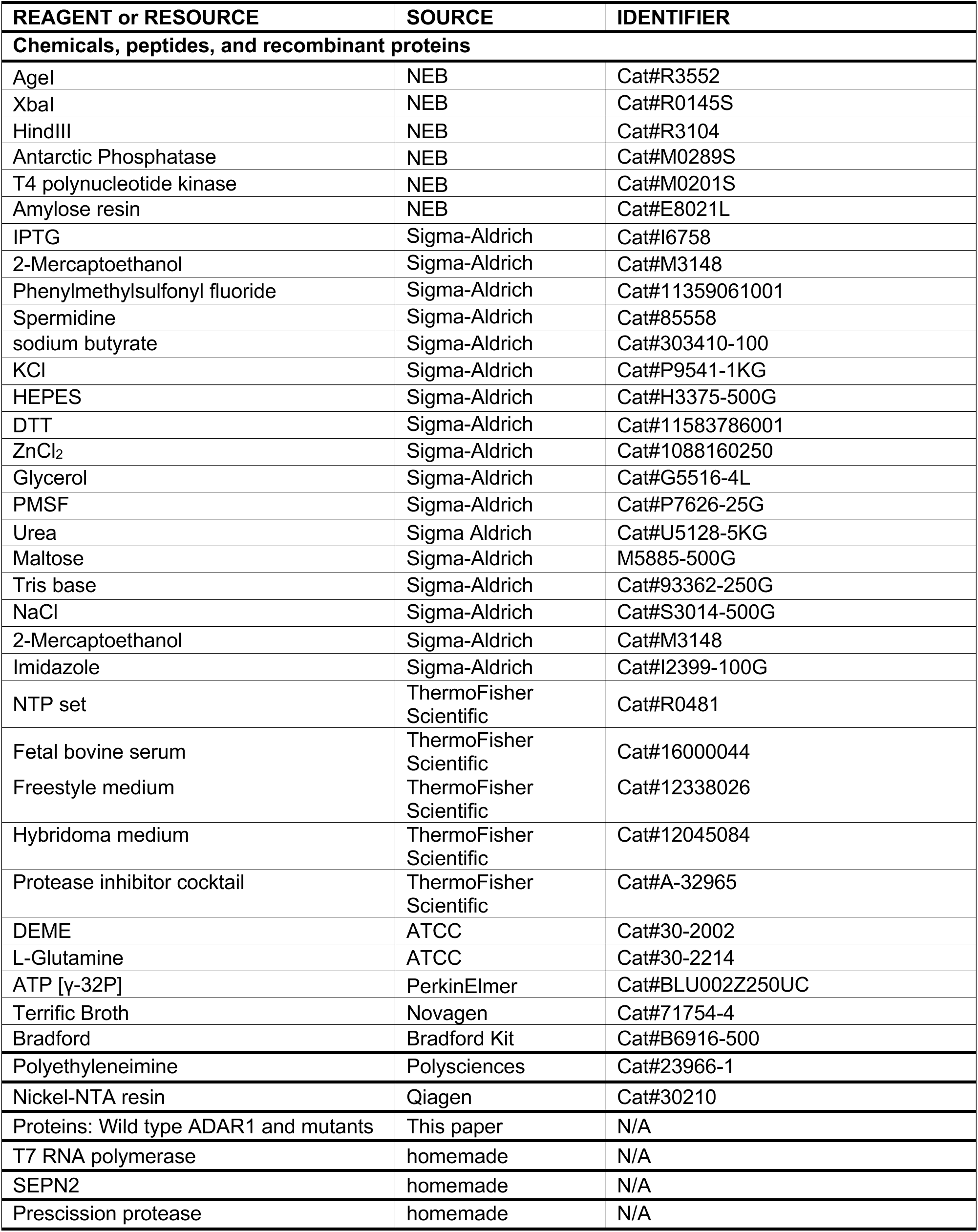
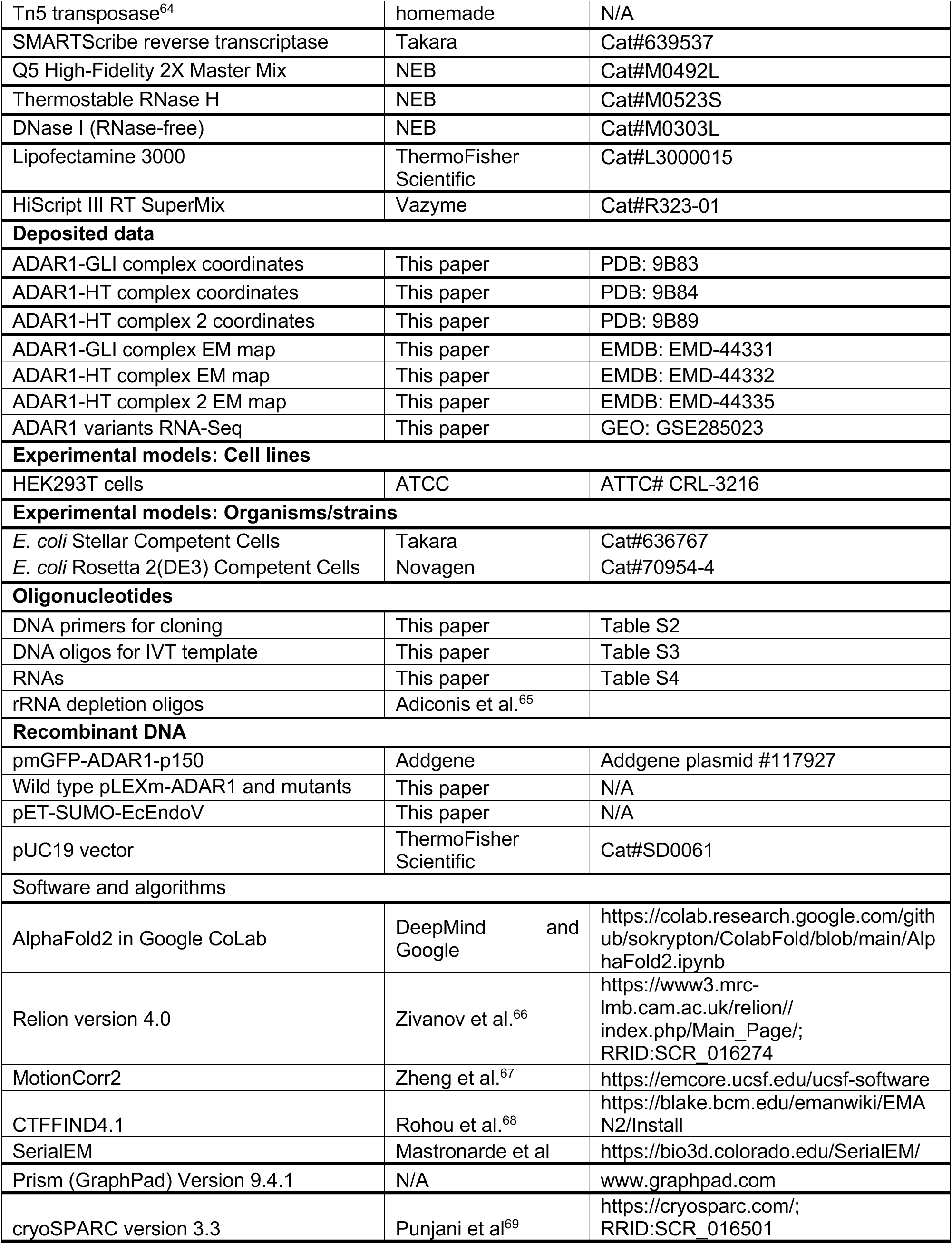
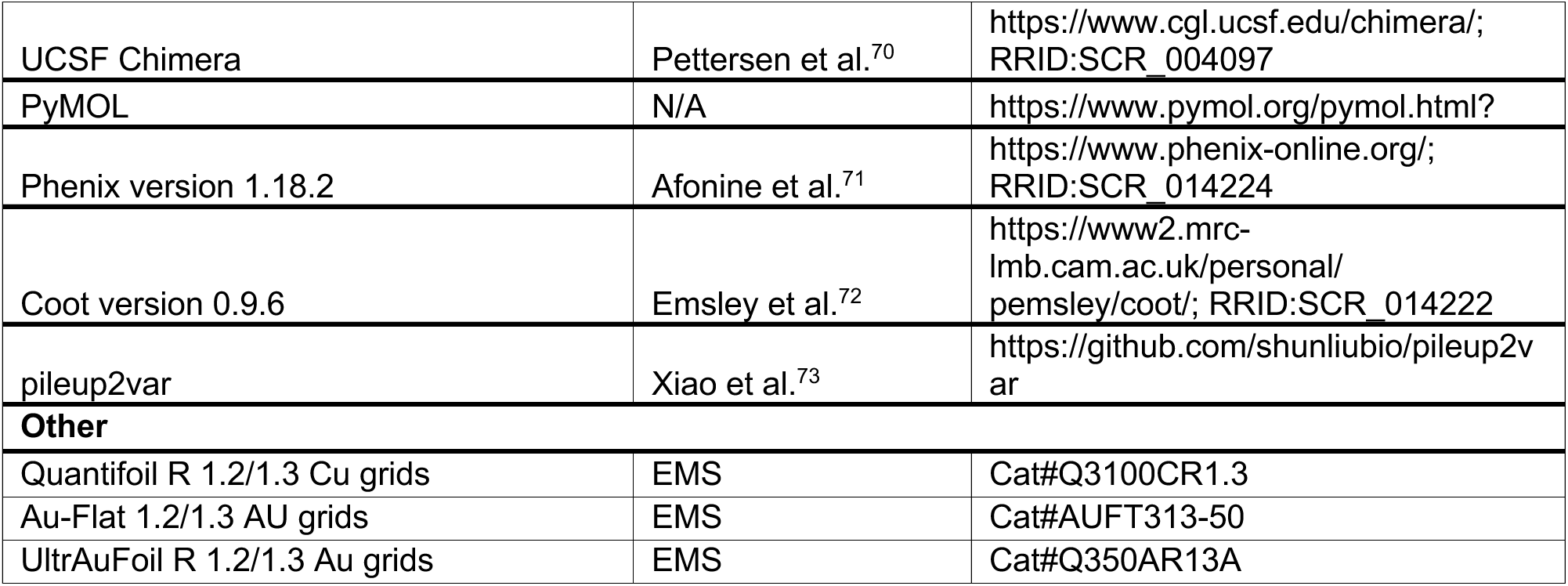

## Methods details

### 1, Protein cloning

The gene of human ADAR1-p150 (pmGFP-ADAR1-p150) was ordered from Addgene (plasmid #117927). The expression plasmids of ADAR1-p150, truncation, or point mutations were constructed by the In-Fusion Snap Assembly method (TaKaRa) into a modified pLEXm mammalian expression vector^74^ with an N-terminal MBP tag following a Prescission cleavage site. The gene of EcEndoV was amplified from the *Escherichia coli* Rosetta (DE3) strain and cloned into a modified pET-SUMO vector (Novage) with an N-terminal His6-SUMO tag following a Sentrin/small ubiquitin-like modifier (SUMO)-specific protease 2 (SEPN2) cleavage site. All the constructions were confirmed by Sanger sequencing (Genewiz). The DNA primers for cloning are listed in Table S2.

### 2, Cell Culture, transfection, and protein expression

For ADAR1 transfection and expression, the HEK293T cell (American Type Culture Collection, catalog CRL-3216) was established as an adherent cell line with the culture medium containing DMEM (ATCC, catalog 30-2002), 10% FBS (Thermo Fisher, catalog 16000044), and 1X L-Glutamine (ATCC, catalog 30-2214) in an incubator set at 37 °C and 5% CO_2_, and the cells were then adapted to the suspension growth with suspension culture containing Freestyle medium (Thermo Fisher, catalog 12338026) and 1% FBS when the cells reached about 50% confluency. The cells were incubated in a shaker set at 37 °C and 5% CO_2_ with 160 rpm/min for suspension growth.

For one liter cells (in a 3L flask) transfection, 1 mg of plasmid and 40 mg fresh-made Polyethyleneimine (PEI, Polysciences, catalog 23966-1) solution (1 mg/ml) were separately mixed with Hybridoma medium (Thermo Fisher, catalog 12045084) to a final volume of 20 ml, then the two solutions were separately filtered with 0.2 μM filter and combined as 40 ml transfection mix, and further incubated at room temperature for 30 minutes before adding it to the suspension cells with density 1.8-2.0 million cells/ml. After 10 hours of transfection, 10 mM sodium butyrate (Sigma, catalog 303410-100) was added to the medium to boost protein expression.

For EcEndoV expression, the recombinant protein was overexpressed in *Escherichia coli* Rosetta (DE3) in the Luria-Bertani (LB) broth medium. The cells were grown at 37 °C until OD600 reached 0.8, then the cells were cooled on ice for 20 min and switched to 25°C and induced with 0.25 mM isopropyl b-D-1-thiogalac-topyranoside (IPTG) for 16-18 hr.

### 3, Protein purification

For ADAR1 purification, the cells were harvested after 60 hours of transfection with centrifugation at 5000xg at 4°C. The cell pellets were resuspended in 60 ml resuspension buffer containing 1M KCl, 50 mM HEPES pH7.5, 2 mM DDT, 0.1 mM ZnCl_2_, 10 % Glycerol, 1 mM PMSF, and 1 tablet protease inhibitor cocktail (Thermo Fisher, catalog J61852.XF). The cells were then lysed by sonication, and centrifuged at 23,000xg for 1 hr at 4 °C. The supernatant was then mixed with 2 ml amylose resin (NEB, catalog E8021L) and incubated for 2 hours in the cold room. After incubation, the supernatant was passed through a gravity flow column at 4 ℃, and the amylose resin was washed with 30 column volumes of resuspension buffer, then the protein was eluted from the column with elution buffer containing 1M KCl, 50 mM HEPES pH7.5, 2 mM DTT, 0.1 mM ZnCl_2_, 5 % Glycerol, 1 mM PMSF and 300 mM maltose (Sigma, catalog M5885-500G), all the buffer was kept cold during purification. The eluted protein was concentrated to 0.5-1 ml and purified by Superose S6 column (GE Healthcare) with gel filtration buffer containing 1M KCl, 50 mM HEPES pH7.5, 2 mM DTT, 0.1 mM ZnCl_2_ and 5 % Glycerol. The ADAR1 mutants were purified by the same method as described above. To study the role of RBDs in ADAR1, the MBP tag was cleaved overnight on ice using homemade Prescission protease and subsequently purified using the same method. ADAR1 and P110 were purified using a Superose S6 column, while the remaining truncation mutants were purified using a Superdex 200 column (GE Healthcare) with the same gel filtration buffer. The purity of ADAR1 proteins after size-exclusion chromatography was assessed by SDS-PAGE (Figures S16A–S16E).

To purify EcEndoV, the cell pellets were harvested through centrifugation at 5000xg and resuspended in buffer A (40 mM Tris-HCl, pH 8.0, 500 mM NaCl, 5 mM beta-mercaptoethanol, 1 mM phenylmethylsulfonyl fluoride). Then cells were lysed by sonication, and centrifuged at 23,000xg for 1 hour at 4°C. The supernatant was then passed through 2 ml Nickel-NTA resin (Qiagen, pre-equilibrated with buffer A), and the resin was washed with 20 column volumes of buffer A. The protein was eluted from the column with buffer B (40 mM Tris-HCl, pH 8.0, 500 mM NaCl, 5 mM beta-mercaptoethanol, 1 mM phenylmethylsulfonyl fluoride, 250 mM imidazole). Next, the SUMO tag was cleaved overnight on ice using homemade SEPN2 protease. After cleavage, the protein was loaded onto a 5 ml heparin column (GE Healthcare) that was pre-equilibrated with buffer C (20 mM Tris-HCl, pH 8.0, 250 mM NaCl, 1 mM DTT). The protein was eluted using a gradient NaCl concentration with buffer D (20 mM Tris-HCl, pH 8.0, 1 M NaCl, 1 mM DTT).

Both purified ADAR1 and EcEndoV proteins were pooled, concentrated, supplemented with 30% glycerol, aliquoted, flash-froze, and stored at -80°C for further use.

### 4, In vitro transcription (IVT) and purification of RNAs

The 96 bp transcription template for GLI-V1 was ordered as six DNA oligos (GLI-F1 to F3 and GLI-R1 to R3, IDT). The oligos were phosphorylated with T4 polynucleotide kinase (10 units/μl; Thermo Fisher), annealed, and ligated into the pUC19 vector (Thermo Fisher) with *HindIII* and *XbaI* restriction sites. The construction was confirmed by Sanger sequencing (Genewiz), and the transcription template was obtained by PCR amplification with primers of GLI-V1-T7F and GLI-V1-R. The PCR product was purified by phenol, followed by ethanol precipitation, air-dried, and dissolved with RNase-free H_2_O. The transcription template of GLI-V2 to GLI-V5 was prepared by PCR amplification using the GLI-V1 construction as the template, with primers of GLI-V2-T7F/R to GLI-V5-T7F/R, respectively. The transcription template for other GLI or HT RNAs was ordered as oligonucleotides from IDT, and the double-stranded transcription template was obtained by annealing the forward and reverse primers at a molar ratio of 1:1. The DNA oligos for preparing RNA transcription templates and corresponding RNA sequences were listed in Table S3 and S4.

The *in vitro* transcription was carried out at 37 °C for 4 hours in a buffer containing 100 mM HEPES potassium (pH7.5), 30 mM DTT, with 20 mm MgCl_2_, 2 mM spermidine, 2.5 mM each NTP, 100 ng/μl DNA template and 2 μM T7 RNA polymerase (homemade). The RNAs were purified by 15% denaturing (7 M urea) polyacrylamide gel electrophoresis (PAGE), extracted, precipitated by ethanol, air-dried, and dissolved with RNase-free H_2_O. The RNA concentration was measured with Nanodrop (DeNovix DS-11), aliquoted, and stored at -20°C for further use.5, 5ʹ-end radiolabeling of RNAs The phosphorylated 5ʹ end of RNAs from IVT was removed by phosphatase (NEB) and purified with Illustra MicroSpin G-50 columns (GE Healthcare), then 2 μM of de-phosphorylated RNA was incubated in a 20 μl reaction mixture containing 1 X T4 polynucleotide kinase buffer, 0.5 μl of [γ-^32^P] ATP (6,000 Ci/mmol; PerkinElmer Life Sciences), and 1 μl of T4 polynucleotide kinase (10 units/μl; Thermo Fisher). The reaction was incubated at 37 °C for 30 min. The labeled product was purified with the RNA Clean-up Kit (NEB, T2030L), and the concentration of 5ʹ-end radiolabeled RNAs was measured with Nanodrop. The RNA was annealed by heating to 85 °C for 5 min and slowly cooling to 4 °C with a PCR machine.

### 6, In vitro editing assay

The in vitro editing assay was carried out by incubating the indicated concentration of 120 nM ADAR1 or mutants with 25 nM of each RNA in buffer containing 10 mM HEPES potassium (pH7.5), 70 mM KCl, 5 % Glycerol, and 1 mM DTT at 37 °C for 45 mins, and the editing reaction was quenched by heating at 85 °C for 3 mins. Then the reaction was further incubated with 100 nM EcEndoV and 2 mM MnCl_2_ at 37 °C for 30 mins and terminated by adding 2 x loading buffer (93.5% formamide, 0.025% xylene cyanol FF, and 50 mM EDTA, pH 8.0) and quenched at 85 °C for 5 mins. Quenched reactions were resolved on 15 % denaturing polyacrylamide gels (Urea-PAGE). The gels were followed by exposure to a phosphorimager plate for 1 hour and imaged using phosphorimaging by the Sapphire Biomolecular Imager (Azure Biosystems). Assays were performed in at least three independent replicates, and the RNA editing percentage was calculated by analyzing the relative band intensities of RNA substrates and products from the same reaction. Specifically, the RNA editing percentage was determined as the ratio of RNA product to the total RNA (substrate + product) in each reaction. The band intensity of the substrates and products were analyzed using Azure Spot (Azure Biosystem) and plotted through GraphPad Prism.

### 7, Electrophoretic mobility shift assay

To quantify the RNA binding affinity of ADAR1 and AGS-associated mutants, assays were carried out in a binding buffer containing 10 mM HEPES potassium (pH 7.5), 70 mM KCl, 5 % Glycerol, and 1 mM DTT. The indicated concentration of ADAR1 or mutants was mixed with 5 nM 5ʹ-end radiolabeled GLI-V11, HT-V5, HT-V6, or HT-V16 in the binding buffer on the ice for 30 min, then each sample was resolved on a 4% native polyacrylamide gel containing 1 x TG buffer (25 mM Tris, 192 mM glycine). The gel was pre-run at 4 °C for 45 min at 150 V with 1 x TG buffer.

To quantify the RNA binding affinity of R3D and D-801, assays were carried out in a binding buffer containing 10 mM HEPES potassium (pH 7.5), 70 mM KCl, 5 % Glycerol, and 1 mM DTT. The indicated concentration of R3D or D-801 was mixed with 10 nM 5ʹ-end radiolabeled HT-V5, HT-V6, or HT-V16 in the binding buffer on the ice for 30 min, then each sample was resolved on a 10% native polyacrylamide gel containing 20 mM HEPES (pH 8.0). The gel was pre-run at 4 °C for 45 min at 150 V with 20 mM HEPES (pH 8.0).

The gels were imaged using phosphorimaging by the Sapphire Biomolecular Imager (Azure Biosystems). The bound and unbound fraction of RNA was quantified by using Azure Spot (Azure Biosystem) and plotted in GraphPad Prism.

### 8, Transfection, RNA extraction, Reverse Transcription PCR, and Sanger sequencing

To transiently transfect HEK293T cells, 1 × 10⁶ cells were seeded in 2 ml of DMEM in a single well of a 6-well plate. After overnight incubation, cells were transfected with 4 µg of plasmids, including wild-type (WT) and mutant ADAR1 variants (P193A, A870T, I872T, R892H, K999N, G1007R, Y1112F, D1113H), as well as an empty vector control, using Lipofectamine 3000 (Thermo Fisher Scientific, Cat. No. L3000015).

Following a 48-hour transfection, cells were harvested, and RNA was extracted using the FastPure Cell/Tissue Total RNA Isolation Kit V2 (Vazyme, Cat. No. RC112-01). 1 µg of the isolated RNA was treated to remove DNA contamination and reverse-transcribed using the HiScript III RT SuperMix for qPCR (+ gDNA wiper) (Vazyme, Cat. No. R323-01).

PCR amplification was performed with 4 µl of cDNA in a total reaction volume of 20 µl using NEB Q5 High-Fidelity DNA Polymerase (New England Biolabs). The PCR program consisted of an initial denaturation at 98°C for 30 seconds followed by 30 cycles of denaturation at 98°C for 10 seconds, annealing at 60°C for 30 seconds, and extension at 72°C for 30 seconds, with a final extension at 72°C for 2 minutes PCR products were purified using the FastPure Gel DNA Extraction Mini Kit (Vazyme, Cat. No. DC301-01) and sent for Sanger sequencing at the Precision Environmental Health DNA Sequencing Core. Editing efficiency was calculated by determining the relative peak height of G/(A+G) in the sequencing chromatograms, expressed as a percentage. The DNA primers used for amplifying GLI cDNA are listed in Table S2.

### 9, RNA library preparation

To prepare rRNA-depleted RNA for library construction, the following procedure was employed. Initially, 12 µl of purified total RNA (1 µg) was combined with an rRNA depletion solution comprising 2 µl of probe hybridization buffer and rRNA depletion solution. The mixture was heated to 95°C for 2 minutes, then gradually cooled to 22°C at a rate of 0.1°C per second and maintained at 22°C for 5 minutes using a pre-heated thermal cycler. Subsequently, an RNase H digestion reaction was prepared by adding 2 µl of RNase H reaction buffer, 2 µl of Thermostable RNase H, and 1 µl of nuclease-free water to the mixture. This reaction was incubated at 37°C for 30 minutes to effectively deplete rRNA. Following this, a DNase I digestion reaction was conducted by adding 5 µl of DNase I reaction buffer, 2.5 µl of RNase-free DNase I, and 22.5 µl of nuclease-free water to the RNase H-treated mixture. This reaction was incubated at 37°C for 30 minutes to eliminate any residual oligonucleotides. Finally, Ampure Beads (1.8x) were employed to purify the rRNA-depleted RNA. The rRNA-depleted RNA was then reverse transcribed into cDNA using SMARTScribe reverse transcriptase and random hexamers (10 µM) at 42°C for 60 minutes. The resulting cDNA was immediately subjected to Tn5 tagmentation following the SMART-seq protocol^75^. Libraries were PCR-amplified using Nextera primer indices and purified with Ampure Beads (0.8x). Finally, the samples were pooled and sequenced on the Illumina NextSeq 550 platform.

### 10, Transcriptome-wide RNA editing analysis

A-to-I RNA editing analysis High-confidence A-to-I RNA editing sites were identified as previously described^76^. Briefly, sequencing reads were trimmed and aligned to the human reference genome (GRCh38) using the STAR aligner. To enable a comparative analysis of editing events among ADAR variants, sequence depth was normalized across all samples. A-to-G mismatches, indicative of A-to-I editing events, were identified using the pileup2var^73^. Editing events mapped to common genomic variants in dbSNP (v151) were discarded. Editing sites were defined with a minimum of five mapped reads at the site and an editing ratio ≥1%.

### 11, Cryo-EM sample preparation

The complex was first assembled by mixing the purified ADAR1 and GLI-V11-8aza or HT-H2-8aza (ordered from Keck Biotechnology Resource Laboratory, Yale University) at a molar ratio of 1:3 in buffer containing 20 mM HEPES, pH 7.5, 500 mM KCl, 1 mM DTT. The complex was incubated on ice for 1 hour, and then slowly diluted KCl from 500 mM to 60 mM with dilution buffer containing 20 mM HEPES, pH 7.5, 1 mM DTT. The complex was further incubated on ice for 30 mins, and then concentrated to 0.7 mg/ml (Amicon Ultra centrifugal filter), as determined by the Bradford assay kit (Bio-Rad). 4 μl freshly purified ADAR1-RNA complex were spotted onto freshly glow-discharged indicated grids (Quantifoil R 1.2/1.3 Cu 300 mesh grids, GF 1.2/1.3 AU 300 mesh grids (Protochips), or Quantifoil R 1.2/1.3 Gold foil on Gold 300 mesh grid). Excess samples were blotted using the Vitrobot Mark IV (FEI) with the standard Vitrobot filter paper (Ø55/20 mm (Ted Pella), the blotting time was set to 2 s, the blotting force was set to 4 and the blotting was done under 100% humidity at 20°C. The grids were flash-frozen in liquid ethane and stored in liquid nitrogen.

### 12, Cryo-EM data collection and processing

The dataset of ADAR1-GLI complex was collected from BCBP Cryo-EM Research Center (Texas A&M University) (5365 micrographs were collected from UltrAuFoil R 1.2/1.3 Au 300 mesh grid, 3880 micrographs were collected from Quantifoil R 1.2/1.3 Cu 300 mesh grids, and 1003 micrographs were collected from GF 1.2/1.3 AU 300 mesh grids). The dataset of the ADAR1-HT complex was collected from Stanford-SLAC Cryo-EM Center (12124 micrographs were collected from the UltrAuFoil R 1.2/1.3 Au 300 mesh grid). Both datasets were recorded on a Titan Krios electron microscope operated at 300 kV, and detailed parameters for data collection were summarized in Table 1. The micrographs were motion-corrected by MotionCor2^67^, and defocus values were estimated on non-dose-weighted micrographs with Gctf. For the processing of the ADAR1-GLI complex dataset, the reference-free auto-picking (Laplacian-of-Gaussian picking) from 100 micrographs and 2D classification was used to obtain good classes as picking templates for reference-base picking through RELION-4.0^77,78^. Around 5476099 particles were picked and extracted to a pixel size of 3.328 Å/pixel and imported to cryoSPARC-4.2 for 2D classification. Around 3222266 particles were selected, and these particles were separated into six classes by ab initials reconstruction. Then the six classes were subjected to heterogeneous refinement, and one class with intact and clear protein and RNA features was selected for subsequent processing. The particles from the good class were extracted (0.832 Å/pixel) in RELION-4.0 and reimported to cryoSPARC-4.2 for iterative ab initials reconstructions and heterogeneous refinement, which yielded a 3.62 Å map after non-uniform refinement. To improve the quality of the map, the 3.62 Å map was used to replace the reference map (the class with clear protein and RNA features) from the first round of ab initials reconstructions, together with the other five classes for new rounds of iterative heterogeneous refinement and ab initials reconstructions (Figure S3), which yielded a map for global resolutions of 3.17 Å with total 181047 particles. The particles were then subjected to global 3D classification in CryoSPARC-4.2 with a global mask and a target resolution of 4 Å, followed by non-uniform refinement of 42,348 particles, which improved the resolution to 3.01 Å. Detailed processing steps for the ADAR1-GLI complex are summarized in Figure S3.

For the processing of the ADAR1-HT complex dataset, a similar method was used to obtain picking templates as the ADAR1-GLI complex dataset, and 20102430 particles were picked and extracted to a pixel size of 3.784 Å/pixel and imported to cryoSPARC-4.2 for 2D classification. Around 3942276 particles were selected, and these particles were subjected to 2 rounds of ab initials reconstructions and one round of heterogeneous refinements to remove junk particles. Then the particles from the good class were extracted (0.832 Å/pixel) in RELION-4.0 and reimported to cryoSPARC-4.2 and further processed with ab initials reconstructions and heterogeneous refinement to remove junk particles, the class with clear protein and RNA features from the heterogeneous refinement was subjected to non-uniform refinement, which yielded a map for global resolutions of 3.38 Å with total 83179 particles. To enhance the map quality, a similar processing strategy as used for the ADAR1-GLI complex was applied, replacing the reference map during heterogeneous refinement. This approach enabled the assignment of more particles to the final high-quality class, characterized by clear protein and RNA features from the initial ab initio reconstructions. A total of 196,253 particles were then subjected to global 3D classification in CryoSPARC-4.2 using a global mask and a target resolution of 4 Å. Subsequent non-uniform refinement of 144,772 particles improved the resolution to 3.2 Å. The processing details for the ADAR1-HT complex are summarized in Figure S4.

For further processing of the ADAR1-HT complex dataset to get the ADAR1-HT complex 2 map, the extracted particles (0.946 Å/pixel) from the first round of ab initials reconstructions and heterogeneous refinement were further separated into six classes by ab initials reconstruction (setting the maximum reconstruction resolution to 6 Å). The class with both protein and RNA features but contains additional density compared to the map of the ADAR1-HT complex was selected to be further separated into three classes by ab initials reconstruction (setting the maximum reconstruction resolution to 5 Å), and followed by heterogeneous refinement, and one class with clear protein and RNA features but without preferred orientation issue was selected for non-uniform refinement which yielded a map of 4.2 Å. To improve the quality of the map, the particles from the class with both protein and RNA features of the first round heterogeneous refinement were extracted to 0.946 Å/pixel and were further separated into six classes by ab initials reconstruction (setting the maximum reconstruction resolution to 5 Å). Then the 3.38 Å map of the ADAR1-HT complex and 4.2 Å map of the ADAR1-HT complex 2 were used to replace the two classes (class 1 and class 5, with clear protein and RNA features) from the ab initials reconstruction for heterogeneous refinement (Figure S4), then the class contains additional density compared to the map of the ADAR1-HT complex was further subjected to non-uniform refinement, followed by local 3D classification using a focused mask for RBD3, which identified four classes with intact RBD3 density out of a total of ten classes. The final non-uniform refinement of 17,460 particles yielded a map at 3.87 Å resolution. The processing details for ADAR1-HT complex 2 are summarized in Figure S4.

### 13, Model building and refinement

For the ADAR1-GLI complex, the model of full-length ADAR1 was downloaded from the AlphaFold Protein Structure Database (ID: AF-P55265-F1). The model of the base-paired GLI-V11 RNA or HT-V2 RNA was generated from Coot-0.9.6 modeling tools^72^. Each monomer of the deaminase domain and RNA was manually docked into the cryo-EM density maps in Chimera^70^. The model was further manually rebuilt in COOT based on electron density and refined in Phenix with real-space refinement and secondary structure and geometry restraints^71^. For the ADAR1-HT complex, the deaminase dimer from the model of the ADAR1-GLI complex and the HT-V2 RNA were separately docked into the ADAR1-HT cryo-EM density map in Chimera, and the model was further manually rebuilt in COOT based on electron density and refined in Phenix. For the ADAR1-HT complex 2, the model of deaminase dimer was from the model of the ADAR1-GLI complex, while the RBD3 and linker were obtained from the model of full-length ADAR1. Each deaminase dimer, RBD3, linker, and HT-V2 RNA was rigid body docked into the 3.87 Å cryo-EM density map and refined in Phenix. All figures were generated by UCSF Chimera and PyMol (http://www.pymol.org). Statistics of all cryo-EM data collection and structure refinement are shown in Table S1.

## References

1. Nishikura, K. (2016). A-to-I editing of coding and non-coding RNAs by ADARs. Nat Rev Mol Cell Biol 17, 83–96. 10.1038/nrm.2015.4.

2. Song, B., Shiromoto, Y., Minakuchi, M., and Nishikura, K. (2022). The role of <SCP>RNA</SCP> editing enzyme ADAR1 in human disease. WIREs RNA 13. 10.1002/wrna.1665.

3. Quin, J., Sedmík, J., Vukić, D., Khan, A., Keegan, L.P., and O’Connell, M.A. (2021). ADAR RNA Modifications, the Epitranscriptome and Innate Immunity. Trends Biochem Sci 46, 758–771. 10.1016/j.tibs.2021.02.002.

4. Rice, G.I., Kasher, P.R., Forte, G.M.A., Mannion, N.M., Greenwood, S.M., Szynkiewicz, M., Dickerson, J.E., Bhaskar, S.S., Zampini, M., Briggs, T.A., et al. (2012). Mutations in ADAR1 cause Aicardi-Goutières syndrome associated with a type I interferon signature. Nat Genet 44, 1243–1248. 10.1038/ng.2414.

5. Tang, Q., Rigby, R.E., Young, G.R., Hvidt, A.K., Davis, T., Tan, T.K., Bridgeman, A., Townsend, A.R., Kassiotis, G., and Rehwinkel, J. (2021). Adenosine-to-inosine editing of endogenous Z-form RNA by the deaminase ADAR1 prevents spontaneous MAVS-dependent type I interferon responses. Immunity 54, 1961–1975.e5. 10.1016/j.immuni.2021.08.011.

6. Nakahama, T., Kato, Y., Shibuya, T., Inoue, M., Kim, J.I., Vongpipatana, T., Todo, H., Xing, Y., and Kawahara, Y. (2021). Mutations in the adenosine deaminase ADAR1 that prevent endogenous Z-RNA binding induce Aicardi-Goutières-syndrome-like encephalopathy. Immunity 54, 1976–1988.e7. 10.1016/j.immuni.2021.08.022.

7. Hartner, J.C., Schmittwolf, C., Kispert, A., Müller, A.M., Higuchi, M., and Seeburg, P.H. (2004). Liver Disintegration in the Mouse Embryo Caused by Deficiency in the RNA-editing Enzyme ADAR1. Journal of Biological Chemistry 279, 4894–4902. 10.1074/jbc.M311347200.

8. Wang, Q., Miyakoda, M., Yang, W., Khillan, J., Stachura, D.L., Weiss, M.J., and Nishikura, K. (2004). Stress-induced Apoptosis Associated with Null Mutation of ADAR1 RNA Editing Deaminase Gene. Journal of Biological Chemistry 279, 4952–4961. 10.1074/jbc.M310162200.

9. Chung, H., Calis, J.J.A., Wu, X., Sun, T., Yu, Y., Sarbanes, S.L., Dao Thi, V.L., Shilvock, A.R., Hoffmann, H.-H., Rosenberg, B.R., et al. (2018). Human ADAR1 Prevents Endogenous RNA from Triggering Translational Shutdown. Cell 172, 811–824.e14. 10.1016/j.cell.2017.12.038.

10. Zhang, T., Yin, C., Fedorov, A., Qiao, L., Bao, H., Beknazarov, N., Wang, S., Gautam, A., Williams, R.M., Crawford, J.C., et al. (2022). ADAR1 masks the cancer immunotherapeutic promise of ZBP1-driven necroptosis. Nature 606, 594–602. 10.1038/s41586-022-04753-7.

11. Ishizuka, J.J., Manguso, R.T., Cheruiyot, C.K., Bi, K., Panda, A., Iracheta-Vellve, A., Miller, B.C., Du, P.P., Yates, K.B., Dubrot, J., et al. (2019). Loss of ADAR1 in tumours overcomes resistance to immune checkpoint blockade. Nature 565, 43–48. 10.1038/s41586-018-0768-9.

12. Booth, B.J., Nourreddine, S., Katrekar, D., Savva, Y., Bose, D., Long, T.J., Huss, D.J., and Mali, P. (2023). RNA editing: Expanding the potential of RNA therapeutics. Molecular Therapy 31, 1533–1549. 10.1016/j.ymthe.2023.01.005.

13. Kaseniit, K.E., Katz, N., Kolber, N.S., Call, C.C., Wengier, D.L., Cody, W.B., Sattely, E.S., and Gao, X.J. (2023). Modular, programmable RNA sensing using ADAR editing in living cells. Nat Biotechnol 41, 482–487. 10.1038/s41587-022-01493-x.

14. Aquino-Jarquin, G. (2020). Novel Engineered Programmable Systems for ADAR-Mediated RNA Editing. Mol Ther Nucleic Acids 19, 1065–1072. 10.1016/j.omtn.2019.12.042.

15. Bass, B.L. (2002). RNA Editing by Adenosine Deaminases That Act on RNA. Annu Rev Biochem 71, 817–846. 10.1146/annurev.biochem.71.110601.135501.

16. Li, Q., Gloudemans, M.J., Geisinger, J.M., Fan, B., Aguet, F., Sun, T., Ramaswami, G., Li, Y.I., Ma, J.-B., Pritchard, J.K., et al. (2022). RNA editing underlies genetic risk of common inflammatory diseases. Nature 608, 569–577. 10.1038/s41586-022-05052-x.

17. Liddicoat, B.J., Piskol, R., Chalk, A.M., Ramaswami, G., Higuchi, M., Hartner, J.C., Li, J.B., Seeburg, P.H., and Walkley, C.R. (2015). RNA editing by ADAR1 prevents MDA5 sensing of endogenous dsRNA as nonself. Science (1979) 349, 1115–1120. 10.1126/science.aac7049.

18. Wong, S.K., Sato, S., and Lazinski, D.W. (2001). Substrate recognition by ADAR1 and ADAR2. RNA 7, S135583820101007X. 10.1017/S135583820101007X.

19. Thuy-Boun, A.S., Thomas, J.M., Grajo, H.L., Palumbo, C.M., Park, S., Nguyen, L.T., Fisher, A.J., and Beal, P.A. (2020). Asymmetric dimerization of adenosine deaminase acting on RNA facilitates substrate recognition. Nucleic Acids Res 48, 7958–7972. 10.1093/nar/gkaa532.

20. Kleinova, R., Rajendra, V., Leuchtenberger, A.F., Lo Giudice, C., Vesely, C., Kapoor, U., Tanzer, A., Derdak, S., Picardi, E., and Jantsch, M.F. (2023). The ADAR1 editome reveals drivers of editing-specificity for ADAR1-isoforms. Nucleic Acids Res 51, 4191– 4207. 10.1093/nar/gkad265.

21. Wang, Y., Park, S., and Beal, P.A. (2018). Selective Recognition of RNA Substrates by ADAR Deaminase Domains. Biochemistry 57, 1640–1651. 10.1021/acs.biochem.7b01100.

22. Zambrano-Mila, M.S., Witzenberger, M., Rosenwasser, Z., Uzonyi, A., Nir, R., Ben-Aroya, S., Levanon, E.Y., and Schwartz, S. (2023). Dissecting the basis for differential substrate specificity of ADAR1 and ADAR2. Nat Commun 14, 8212. 10.1038/s41467-023-43633-0.

23. Uzonyi, A., Nir, R., Shliefer, O., Stern-Ginossar, N., Antebi, Y., Stelzer, Y., Levanon, E.Y., and Schwartz, S. (2021). Deciphering the principles of the RNA editing code via large-scale systematic probing. Mol Cell 81, 2374–2387.e3. 10.1016/j.molcel.2021.03.024.

24. Mboukou, A., Rajendra, V., Messmer, S., Mandl, T.C., Catala, M., Tisné, C., Jantsch, M.F., and Barraud, P. (2024). Dimerization of ADAR1 modulates site-specificity of RNA editing. Nat Commun 15, 10051. 10.1038/s41467-024-53777-2.

25. Ramaswami, G., and Li, J.B. (2014). RADAR: a rigorously annotated database of A-to-I RNA editing. Nucleic Acids Res 42, D109–D113. 10.1093/nar/gkt996.

26. Porath, H.T., Carmi, S., and Levanon, E.Y. (2014). A genome-wide map of hyper-edited RNA reveals numerous new sites. Nat Commun 5, 4726. 10.1038/ncomms5726.

27. Levanon, E.Y., Eisenberg, E., Yelin, R., Nemzer, S., Hallegger, M., Shemesh, R., Fligelman, Z.Y., Shoshan, A., Pollock, S.R., Sztybel, D., et al. (2004). Systematic identification of abundant A-to-I editing sites in the human transcriptome. Nat Biotechnol 22, 1001–1005. 10.1038/nbt996.

28. Hundley, H.A., and Bass, B.L. (2010). ADAR editing in double-stranded UTRs and other noncoding RNA sequences. Trends Biochem Sci 35, 377–383. 10.1016/j.tibs.2010.02.008.

29. Morse, D.P., Aruscavage, P.J., and Bass, B.L. (2002). RNA hairpins in noncoding regions of human brain and Caenorhabditis elegans mRNA are edited by adenosine deaminases that act on RNA. Proceedings of the National Academy of Sciences 99, 7906–7911. 10.1073/pnas.112704299.

30. Kiran, A.M., O’Mahony, J.J., Sanjeev, K., and Baranov, P. V. (2012). Darned in 2013: inclusion of model organisms and linking with Wikipedia. Nucleic Acids Res 41, D258– D261. 10.1093/nar/gks961.

31. Eggington, J.M., Greene, T., and Bass, B.L. (2011). Predicting sites of ADAR editing in double-stranded RNA. Nat Commun 2, 319. 10.1038/ncomms1324.

32. Polson, A.G., and Bass, B.L. (1994). Preferential selection of adenosines for modification by double-stranded RNA adenosine deaminase. EMBO J 13, 5701–5711. 10.1002/j.1460-2075.1994.tb06908.x.

33. Lehmann, K.A., and Bass, B.L. (2000). Double-Stranded RNA Adenosine Deaminases ADAR1 and ADAR2 Have Overlapping Specificities. Biochemistry 39, 12875–12884. 10.1021/bi001383g.

34. Zhang, R., Deng, P., Jacobson, D., and Li, J.B. (2017). Evolutionary analysis reveals regulatory and functional landscape of coding and non-coding RNA editing. PLoS Genet 13, e1006563. 10.1371/journal.pgen.1006563.

35. Sapiro, A.L., Deng, P., Zhang, R., and Li, J.B. (2015). Cis Regulatory Effects on A-to-I RNA Editing in Related Drosophila Species. Cell Rep 11, 697–703. 10.1016/j.celrep.2015.04.005.

36. Liu, X., Sun, T., Shcherbina, A., Li, Q., Jarmoskaite, I., Kappel, K., Ramaswami, G., Das, R., Kundaje, A., and Li, J.B. (2021). Learning cis-regulatory principles of ADAR-based RNA editing from CRISPR-mediated mutagenesis. Nat Commun 12, 2165. 10.1038/s41467-021-22489-2.

37. Jumper, J., Evans, R., Pritzel, A., Green, T., Figurnov, M., Ronneberger, O., Tunyasuvunakool, K., Bates, R., Žídek, A., Potapenko, A., et al. (2021). Highly accurate protein structure prediction with AlphaFold. Nature 596, 583–589. 10.1038/s41586-021-03819-2.

38. Cho, D.-S.C., Yang, W., Lee, J.T., Shiekhattar, R., Murray, J.M., and Nishikura, K. (2003). Requirement of Dimerization for RNA Editing Activity of Adenosine Deaminases Acting on RNA. Journal of Biological Chemistry 278, 17093–17102. 10.1074/jbc.M213127200.

39. Ota, H., Sakurai, M., Gupta, R., Valente, L., Wulff, B.-E., Ariyoshi, K., Iizasa, H., Davuluri, R.V., and Nishikura, K. (2013). ADAR1 Forms a Complex with Dicer to Promote MicroRNA Processing and RNA-Induced Gene Silencing. Cell 153, 575–589. 10.1016/j.cell.2013.03.024.

40. Sebastian Vik, E., Sameen Nawaz, M., Strøm Andersen, P., Fladeby, C., Bjørås, M., Dalhus, B., and Alseth, I. (2013). Endonuclease V cleaves at inosines in RNA. Nat Commun 4, 2271. 10.1038/ncomms3271.

41. Shimokawa, T., Rahman, M.F.-U., Tostar, U., Sonkoly, E., Ståhle, M., Pivarcsi, A., Palaniswamy, R., and Zaphiropoulos, P.G. (2013). RNA editing of the GLI1 transcription factor modulates the output of Hedgehog signaling. RNA Biol 10, 321–333. 10.4161/rna.23343.

42. Matthews, M.M., Thomas, J.M., Zheng, Y., Tran, K., Phelps, K.J., Scott, A.I., Havel, J., Fisher, A.J., and Beal, P.A. (2016). Structures of human ADAR2 bound to dsRNA reveal base-flipping mechanism and basis for site selectivity. Nat Struct Mol Biol 23, 426–433. 10.1038/nsmb.3203.

43. Park, S., Doherty, E.E., Xie, Y., Padyana, A.K., Fang, F., Zhang, Y., Karki, A., Lebrilla, C.B., Siegel, J.B., and Beal, P.A. (2020). High-throughput mutagenesis reveals unique structural features of human ADAR1. Nat Commun 11, 5130. 10.1038/s41467-020-18862-2.

44. Macbeth, M.R., Schubert, H.L., VanDemark, A.P., Lingam, A.T., Hill, C.P., and Bass, B.L. (2005). Inositol Hexakisphosphate Is Bound in the ADAR2 Core and Required for RNA Editing. Science (1979) 309, 1534–1539. 10.1126/science.1113150.

45. Kuttan, A., and Bass, B.L. (2012). Mechanistic insights into editing-site specificity of ADARs. Proceedings of the National Academy of Sciences 109. 10.1073/pnas.1212548109.

46. Wang, Y., Havel, J., and Beal, P.A. (2015). A Phenotypic Screen for Functional Mutants of Human Adenosine Deaminase Acting on RNA 1. ACS Chem Biol 10, 2512–2519. 10.1021/acschembio.5b00711.

47. Karki, A., Campbell, K.B., Mozumder, S., Fisher, A.J., and Beal, P.A. (2024). Impact of Disease-Associated Mutations on the Deaminase Activity of ADAR1. Biochemistry 63, 282–293. 10.1021/acs.biochem.3c00405.

48. Guo, X., Wiley, C.A., Steinman, R.A., Sheng, Y., Ji, B., Wang, J., Zhang, L., Wang, T., Zenatai, M., Billiar, T.R., et al. (2021). Aicardi-Goutières syndrome-associated mutation at ADAR1 gene locus activates innate immune response in mouse brain. J Neuroinflammation 18, 169. 10.1186/s12974-021-02217-9.

49. Guo, X., Steinman, R.A., Sheng, Y., Cao, G., Wiley, C.A., and Wang, Q. (2022). An AGS-associated mutation in ADAR1 catalytic domain results in early-onset and MDA5-dependent encephalopathy with IFN pathway activation in the brain. J Neuroinflammation 19, 285. 10.1186/s12974-022-02646-0.

50. Mannion, N.M., Greenwood, S.M., Young, R., Cox, S., Brindle, J., Read, D., Nellåker, C., Vesely, C., Ponting, C.P., McLaughlin, P.J., et al. (2014). The RNA-Editing Enzyme ADAR1 Controls Innate Immune Responses to RNA. Cell Rep 9, 1482–1494. 10.1016/j.celrep.2014.10.041.

51. Ha, S.C., Choi, J., Hwang, H.-Y., Rich, A., Kim, Y.-G., and Kim, K.K. (2009). The structures of non-CG-repeat Z-DNAs co-crystallized with the Z-DNA-binding domain, hZα ADAR1. Nucleic Acids Res 37, 629–637. 10.1093/nar/gkn976.

52. Lee, A.-R., Kim, N.-H., Seo, Y.-J., Choi, S.-R., and Lee, J.-H. (2018). Thermodynamic Model for B-Z Transition of DNA Induced by Z-DNA Binding Proteins. Molecules 23, 2748. 10.3390/molecules23112748.

53. Lorenz, R., Bernhart, S.H., Höner zu Siederdissen, C., Tafer, H., Flamm, C., Stadler, P.F., and Hofacker, I.L. (2011). ViennaRNA Package 2.0. Algorithms for Molecular Biology 6, 26. 10.1186/1748-7188-6-26.

54. Tay, D.J.T., Song, Y., Peng, B., Toh, T.B., Hooi, L., Toh, D.-F.K., Hong, H., Tang, S.J., Han, J., Gan, W.L., et al. (2021). Targeting RNA editing of antizyme inhibitor 1: A potential oligonucleotide-based antisense therapy for cancer. Molecular Therapy 29, 3258–3273. 10.1016/j.ymthe.2021.05.008.

55. Gabay, O., Shoshan, Y., Kopel, E., Ben-Zvi, U., Mann, T.D., Bressler, N., Cohen-Fultheim, R., Schaffer, A.A., Roth, S.H., Tzur, Z., et al. (2022). Landscape of adenosine-to-inosine RNA recoding across human tissues. Nat Commun 13, 1184. 10.1038/s41467-022-28841-4.

56. Valente, L., and Nishikura, K. (2007). RNA Binding-independent Dimerization of Adenosine Deaminases Acting on RNA and Dominant Negative Effects of Nonfunctional Subunits on Dimer Functions. Journal of Biological Chemistry 282, 16054–16061. 10.1074/jbc.M611392200.

57. Shiromoto, Y., Sakurai, M., Minakuchi, M., Ariyoshi, K., and Nishikura, K. (2021). ADAR1 RNA editing enzyme regulates R-loop formation and genome stability at telomeres in cancer cells. Nat Commun 12, 1654. 10.1038/s41467-021-21921-x.

58. Nishikura, K. (2010). Functions and Regulation of RNA Editing by ADAR Deaminases. Annu Rev Biochem 79, 321–349. 10.1146/annurev-biochem-060208-105251.

59. Bazak, L., Haviv, A., Barak, M., Jacob-Hirsch, J., Deng, P., Zhang, R., Isaacs, F.J., Rechavi, G., Li, J.B., Eisenberg, E., et al. (2014). A-to-I RNA editing occurs at over a hundred million genomic sites, located in a majority of human genes. Genome Res 24, 365–376. 10.1101/gr.164749.113.

60. Kallman, A.M. (2003). ADAR2 A-&gt;I editing: site selectivity and editing efficiency are separate events. Nucleic Acids Res 31, 4874–4881. 10.1093/nar/gkg681.

61. Stefl, R., Oberstrass, F.C., Hood, J.L., Jourdan, M., Zimmermann, M., Skrisovska, L., Maris, C., Peng, L., Hofr, C., Emeson, R.B., et al. (2010). The Solution Structure of the ADAR2 dsRBM-RNA Complex Reveals a Sequence-Specific Readout of the Minor Groove. Cell 143, 225–237. 10.1016/j.cell.2010.09.026.

62. Pestal, K., Funk, C.C., Snyder, J.M., Price, N.D., Treuting, P.M., and Stetson, D.B. (2015). Isoforms of RNA-Editing Enzyme ADAR1 Independently Control Nucleic Acid Sensor MDA5-Driven Autoimmunity and Multi-organ Development. Immunity 43, 933– 944. 10.1016/j.immuni.2015.11.001.

63. Hu, S.-B., Heraud-Farlow, J., Sun, T., Liang, Z., Goradia, A., Taylor, S., Walkley, C.R., and Li, J.B. (2023). ADAR1p150 prevents MDA5 and PKR activation via distinct mechanisms to avert fatal autoinflammation. Mol Cell 83, 3869–3884.e7. 10.1016/j.molcel.2023.09.018.

64. Picelli, S., Björklund, Å.K., Reinius, B., Sagasser, S., Winberg, G., and Sandberg, R. (2014). Tn5 transposase and tagmentation procedures for massively scaled sequencing projects. Genome Res 24, 2033–2040. 10.1101/gr.177881.114.

65. Adiconis, X., Borges-Rivera, D., Satija, R., DeLuca, D.S., Busby, M.A., Berlin, A.M., Sivachenko, A., Thompson, D.A., Wysoker, A., Fennell, T., et al. (2013). Comparative analysis of RNA sequencing methods for degraded or low-input samples. Nat Methods 10, 623–629. 10.1038/nmeth.2483.

66. Zivanov, J., Nakane, T., and Scheres, S.H.W. (2020). Estimation of high-order aberrations and anisotropic magnification from cryo-EM data sets in RELION-3.1. IUCrJ 7, 253–267. 10.1107/S2052252520000081.

67. Zheng, S.Q., Palovcak, E., Armache, J.-P., Verba, K.A., Cheng, Y., and Agard, D.A. (2017). MotionCor2: anisotropic correction of beam-induced motion for improved cryo-electron microscopy. Nat Methods 14, 331–332. 10.1038/nmeth.4193.

68. Rohou, A., and Grigorieff, N. (2015). CTFFIND4: Fast and accurate defocus estimation from electron micrographs. J Struct Biol 192, 216–221. 10.1016/j.jsb.2015.08.008.

69. Punjani, A., Rubinstein, J.L., Fleet, D.J., and Brubaker, M.A. (2017). cryoSPARC: algorithms for rapid unsupervised cryo-EM structure determination. Nat Methods 14, 290–296. 10.1038/nmeth.4169.

70. Pettersen, E.F., Goddard, T.D., Huang, C.C., Meng, E.C., Couch, G.S., Croll, T.I., Morris, J.H., and Ferrin, T.E. (2021). <SCP>UCSF ChimeraX</SCP> : Structure visualization for researchers, educators, and developers. Protein Science 30, 70–82. 10.1002/pro.3943.

71. Afonine, P. V., Poon, B.K., Read, R.J., Sobolev, O. V., Terwilliger, T.C., Urzhumtsev, A., and Adams, P.D. (2018). Real-space refinement in *PHENIX* for cryo-EM and crystallography. Acta Crystallogr D Struct Biol 74, 531–544. 10.1107/S2059798318006551.

72. Emsley, P., and Cowtan, K. (2004). Coot: Model-building tools for molecular graphics. Acta Crystallogr D Biol Crystallogr 60, 2126–2132. 10.1107/S0907444904019158.

73. Xiao, Y.-L., Liu, S., Ge, R., Wu, Y., He, C., Chen, M., and Tang, W. (2023). Transcriptome-wide profiling and quantification of N6-methyladenosine by enzyme-assisted adenosine deamination. Nat Biotechnol 41, 993–1003. 10.1038/s41587-022-01587-6.

74. Aricescu, A.R., Lu, W., and Jones, E.Y. (2006). A time- and cost-efficient system for high-level protein production in mammalian cells. Acta Crystallogr D Biol Crystallogr 62, 1243–1250. 10.1107/S0907444906029799.

75. Picelli, S., Faridani, O.R., Björklund, Å.K., Winberg, G., Sagasser, S., and Sandberg, R. (2014). Full-length RNA-seq from single cells using Smart-seq2. Nat Protoc 9, 171–181. 10.1038/nprot.2014.006.

76. Xiang, J.-F., Yang, Q., Liu, C.-X., Wu, M., Chen, L.-L., and Yang, L. (2018). N6-Methyladenosines Modulate A-to-I RNA Editing. Mol Cell 69, 126–135.e6. 10.1016/j.molcel.2017.12.006.

77. Kimanius, D., Dong, L., Sharov, G., Nakane, T., and Scheres, S.H.W. (2021). New tools for automated cryo-EM single-particle analysis in RELION-4.0. Biochemical Journal 478, 4169–4185. 10.1042/BCJ20210708.

78. Zivanov, J., Nakane, T., and Scheres, S.H.W. (2020). Estimation of high-order aberrations and anisotropic magnification from cryo-EM data sets in RELION-3.1. IUCrJ 7, 253–267. 10.1107/S2052252520000081.

